# Extra-lineage tissue programs define the transcription states of human pancreatic cancer

**DOI:** 10.64898/2026.04.29.721655

**Authors:** Sabrina Ge, Paul Tonon, Jingxiong Xu, Gun Ho Jang, Ferris Nowlan, Jimin Min, Karen Ng, Eugenia Flores Figueroa, Amy Zhang, Michelle Chan-Seng-Yue, Yuanchang Fang, Adriana Migliorini, Julie M. Wilson, Anna Dodd, Sebastian Arcila-Barrera, Ayah Elqaderi, Ilinca Lungu, Yu Zhang, Stephanie Ramotar, Shawn Hutchinson, Daniela Bevacqua, Ayelet Borgida, Spring Holter, Pathum Kossinna, Ruth Isserlin, Veronique Voisin, Andreas Mund, Maria Cristina Nostro, Erica S. Tsang, Robert C. Grant, Grainne M. O’Kane, David A. Tuveson, Oren Parnas, Federico Gaiti, Nina Steele, Jennifer J. Knox, Anirban Maitra, Hartland W. Jackson, Steven Gallinger, Faiyaz Notta

## Abstract

Cancers acquire alternate transcriptional states as they evolve, but the origins, timing and determinants of this plasticity are poorly understood in many tumours. We investigated the transcriptional states of pancreatic cancer by integrating ∼1000 tumour-enriched genomes and transcriptomes from 464 patients combined with scRNA-seq, multiome profiling, and spatial proteomics. Four epithelial states covering the spectrum of lineage plasticity were identified (Classical-1, Classical-2, Basal-1, Basal-2). Comparing these states to normal and pan-cancer human single cell atlases showed each state reflects distinct tissue programs found in other malignancies. Single cell analysis uncovered that the main transcription state of this disease (Classical-1) emerges before KRAS mutations. Spatial proteomics from patients and cancer-free donors showed that the Classical-1 program emerges during acinar-to-ductal metaplasia, and also unexpectedly, in normal ducts without disrupting their morphology. Overall, these findings link the extensive lineage plasticity potential of this organ to the origins of the transcriptional states.

---

Although pancreatic ductal adenocarcinoma (herein pancreatic cancer) is defined by a uniform set of driver mutations (*KRAS, CDKN2A, TP53, SMAD4*), it evolves into a transcriptionally complex disease. The transcriptional framework of these tumours is described by two major programs, Basal and Classical (Moffitt et al., 2015), which are widely used to classify tumours and predict clinical outcome. However, their biological basis is not fully understood. One possibility is that these programs reflect differences in the pancreatic cell-of-origin, and this is supported by mouse models that show that duct versus acinar cells give rise to these divergent transcriptional programs (Flowers et al., 2021; Lee et al., 2019). By contrast, studies of human tumours suggest that these states are not fixed but instead that the Classical program transitions into Basal with progression (Chan-Seng-Yue et al., 2020; Hayashi et al., 2020). Importantly, pancreatic cells harbour extensive lineage plasticity and these transcription programs may reflect both cellular origins and reprogramming events during tumour evolution (Alonso-Curbelo et al., 2021; Burdziak et al., 2023; Miyabayashi et al., 2020; Schlesinger et al., 2020). From human tumours, it remains unclear when and how these transcriptional states are established in tumourigenesis. This has been difficult to address due to technical limitations.

A number of single cell studies have shown that ‘Basal/Classical’ underestimate the plasticity in this disease (Hwang et al., 2022; Loveless et al., 2025; Raghavan et al., 2021). Accurately resolving the transcriptional architecture of pancreatic cancer has been challenging for several reasons. First, these tumours have very low cellularity due to extensive desmoplasia, and the surrounding tissue express extreme levels of enzymatic genes. 90-95% of transcription in acinar cells is dedicated to fewer than 30 proteins (amylases, trypsinogens, lipases, elastases; Harding et al., 1977), and thus, even residual contamination of these cells can influence detection of tumour-related transcription programs (Bailey et al., 2016; Collisson et al., 2011; Raphael et al., 2017). Second, most studies lack adequate representation of metastatic patients due to tissue access. This patient population is critical to profile as they compose the majority of diagnoses and harbour more aggressive tumours. Single-cell approaches can address some cellularity limitations, but high dropout rates can miss transcription signals. Moreover, single cell analysis at the scale of hundreds of patients requires merged of independent studies, which often requires extensive normalization and possible loss of signal. Collectively, these issues have limited our ability to study the transcriptional architecture of this disease. In this study, we assembled a comprehensively annotated patient cohort and employed a number of external mouse and human datasets to validate our findings. Building on our initial study of 240 patients (Chan-Seng-Yue et al., 2020), we doubled the patient cohort of laser-capture microdissected (LCM) tumours with matched RNA-seq and whole-genome sequencing (WGS) (∼1000 total samples). This collection includes both early-stage and metastatic patients representing a decade’s worth of sample collection (Aung et al., 2018; Chan-Seng-Yue et al., 2020; Knox et al., 2025; Notta et al., 2016; O’Kane et al., 2020). These data were integrated with single-cell transcriptomics (47 patients, 161,363 cells), single-nucleus Multiome profiling (8 patients, >7,800 cells), patient organoid profiling (n=164), and spatial proteomics of patients and cancer-free donors.

## Cross-platform identification of tumour transcriptional states

Our cohort is comprised of 490 matched whole genomes and transcriptomes from 464 donors that were laser-captured (**Table S1-2**). The median tumour cellularity of our cohort was 76.9% determined by WGS. To ensure transcriptional programs were present in the tumour, we also generated scRNA-seq on 48 tumours (47 donors), including 32 primary tumours and 16 metastases (15 liver, 1 lung; **Fig. S1A** and **Table S3**). To address low cellularity in single cell analysis, we used a negative selection strategy before scRNA-seq to deplete immune and stromal cells (Methods). We sequenced 161,363 high quality single cells. To distinguish neoplastic from non-neoplastic cells, we used both tumour markers and inference of aneuploidy by tracking copy number burden (**Fig S1A-C**). Copy number profiles were generated using Numbat, a haplotype-aware model that integrates allelic expression imbalance to copy number resolution (**Fig. S1B**). Using this approach, 84,239 of the 161,363 cells were neoplastic. Single nucleus Multiome profiling (snMultiome; ATAC and RNA-seq) on 8 additional patients (**Table S4**) allowed us to analyze regulatory changes related to chromatin accessibility.

A number of transcription schemes have been published for pancreatic cancer (Bailey et al., 2016; Collisson et al., 2011; Maurer et al., 2019; Puleo et al., 2018) and all converge on the Basal/Classical model (Moffitt et al., 2015). The original classification originated from microarray and RNA-seq data from bulk tissue with low cellularity. Although LCM helps to clarify tumour transcriptional signals, residual stromal cells can still dominate the expression profile due to high transcript abundance in particular cell types (e.g. enzymes in acinar cells, collagens in fibroblasts, or cytokines from immune cells). We implemented the following strategy. First, non-negative matrix factorization (NMF) was applied to our LCM RNA-seq data to separate tumour from stromal signals (**Fig. 1A**). Using bulk RNA-seq in the first step of the analysis is critical as the breath of transcript coverage is substantially greater compared to scRNA-seq (bulk - 15,000-25,000 genes per sample; single-cell - 1,000-6,000 genes per cell). Moreover, using 490 patients allows us to capture rarer patient subgroups. However, we did combine bulk RNA-seq analysis later with scRNA-seq to ensure that the extracted RNA signatures originated from tumour cells.

**Figure 1:**
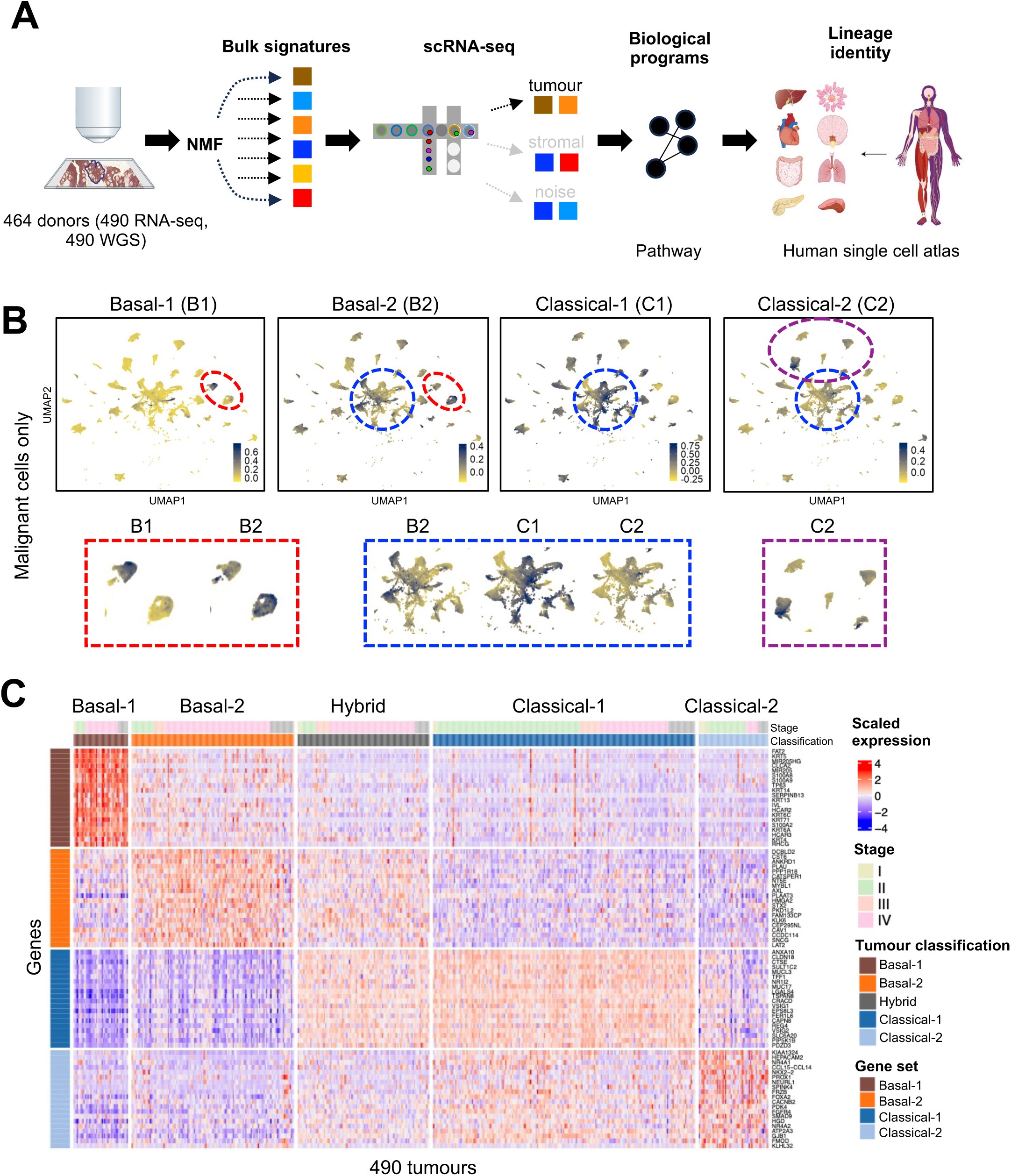
Four robust pancreatic cancer programs identified from 490 tumours. (a) Workflow of analysis of human pancreatic ductal adenocarcinoma samples. Transcriptional data supported by genetic data is deconvoluted using NMF into signatures. Tumour programs derived from these signatures are validated with scRNA-seq and biological meaning is derived through pathway enrichment and comparison with the human single cell atlas. (b) UMAP of malignant cells scored with the tumour programs. Circled sections of the UMAP are shown below and highlight differences in expression of the tumour programs. (c) Heatmap showing expression of tumour program gene sets genes across all 490 bulk tumours. Expression is scaled by z-score. Tumours are annotated above the heatmap by stage and classification.

NMF was applied across a range of factorization ranks (k) to identify the minimal k at which tumour and stromal signatures separated. Multiple considerations went into selecting the rank. For example, at lower k values (e.g., k < 10), the Classical program often fragmented into multiple signatures that all showed strong statistical significance (k=5 **Fig. S1A**). At low rank, the Basal signature was intermixed with the fibroblast signature (k=7, **Fig. S1A**). By k=15, these issues resolved and additional transcriptional structure became evident. Also, the stromal signatures also clearly separated at this rank. Knowing that transcription signatures converge on the Basal/Classical framework, we used our previous work to guide signature identification. Four transcription signatures related to the Basal/Classical framework were identified (Basal-1, Basal-2, Classical-1, Classical-2). The Classical-2 signature retained a considerable number of acinar cell genes at k=15, which was not present at k=16 (**Fig. S1B**). These newly extracted signatures were validated in single cells to confirm that each signature mapped to neoplastic cells (**Fig. S3**). Integrating bulk and single-cell data also helped to confirm the interpretation of each signature. For example, even though Basal-1 and Basal-2 were different RNA signatures, they both aligned to the previous Basal signature (hence their labels Basal-1 and Basal-2; **Fig. S3**). By contrast, the Classical-2 signature is distinct from the previous Classical program based on single cell analysis (**Fig. S3**). This is not unexpected as the Classical-2 signature is in a rare subset of tumours and is often contaminated with acinar cell genes. Importantly, all four signatures were found in distinct malignant cell clusters from independent patients (**Fig. 1B**), supporting these are discrete transcriptional tumour states.

We compared the tumour classification derived from these new signatures to our previously published study (Chan-Seng-Yue et al., 2020; Basal-A, Basal-B, Classical-A, Classical-B). Concordance was quantified using consensus probability where 200 iterations were run to calculate the stability of sample assignment. In the original 2020 classification, there were a number of tumours assigned with low confidence (**Fig. S4A**, 2020). This included Hybrid tumours that had a strong Basal-A or Basal-B signal, and Classical-B tumours with a significant Classical-A component (see arrows). The new signatures significantly improved the overall assignment of tumours (**Fig. S4B**, 2026). The effect was most pronounced for the Classical-2 and Hybrid categories. Hybrid tumours were particularly difficult to resolve in our 2020 classification. Using the new signatures, Hybrid tumours were found to consist mainly of Basal-2 and Classical-1 programs and now stand as a distinct patient subgroup. The patient heatmap using the new signatures now shows well-defined patient subgroups (**Fig. 1C**). Clinically, we confirmed that Basal-1/2 programs are enriched in metastatic tumours, whereas Classical-1/2 are enriched in resectable disease (**Fig. S4C**). However, clinical investigation of these programs will be reported in a separate study.

## Biological identity of each transcription state

We began with an in-depth examination of gene composition, pathways, and lineage features of each transcription signature using a human single-cell atlas (Pan et al., 2023) and pan-cancer single-cell atlas (Gavish et al., 2023).

### i. Classical-1 is a pan-foregut secretory gastrointestinal state and not a ductal progenitor program

The Classical-1 signature is the main transcriptional program of pancreatic cancer and we sought to define its biological basis. The term ‘Classical’ has evolved to describe a common case of pancreatic cancer, and in the literature, has also been described as ‘gastric’ or ‘progenitor-like’ (Bailey et al., 2016; Benitz et al., 2024). We defined this program without using these previous interpretations.

A central tenet of cancer is that tumour transcription states reflect the lineage of the tissue in which they originate. In this regard, the Classical-1 program would be assumed to resemble the normal pancreatic duct since these are ductal adenocarcinomas. To determine this, we collected scRNA-seq datasets from normal human fetal, neonatal, and adult pancreas (Migliorini et al., 2024; Tosti et al., 2021). We identified 160,568 cells spanning endocrine and exocrine lineages and scored them for Classical-1 signature (**Fig. 2A**). The Classical-1 program was absent in the pancreatic lineage regardless of developmental state and was only detected in rare adult ductal cells (0/11119 fetal; 0/2470 neonatal; 40/20835, 0.19% adult ductal cells; **Fig 2A inset**). The observation that Classical-1+ ductal cells are essential absent in early pancreas development and only present in rare adult cells supports this program is not a developmental ductal progenitor state.

**Figure 2:**
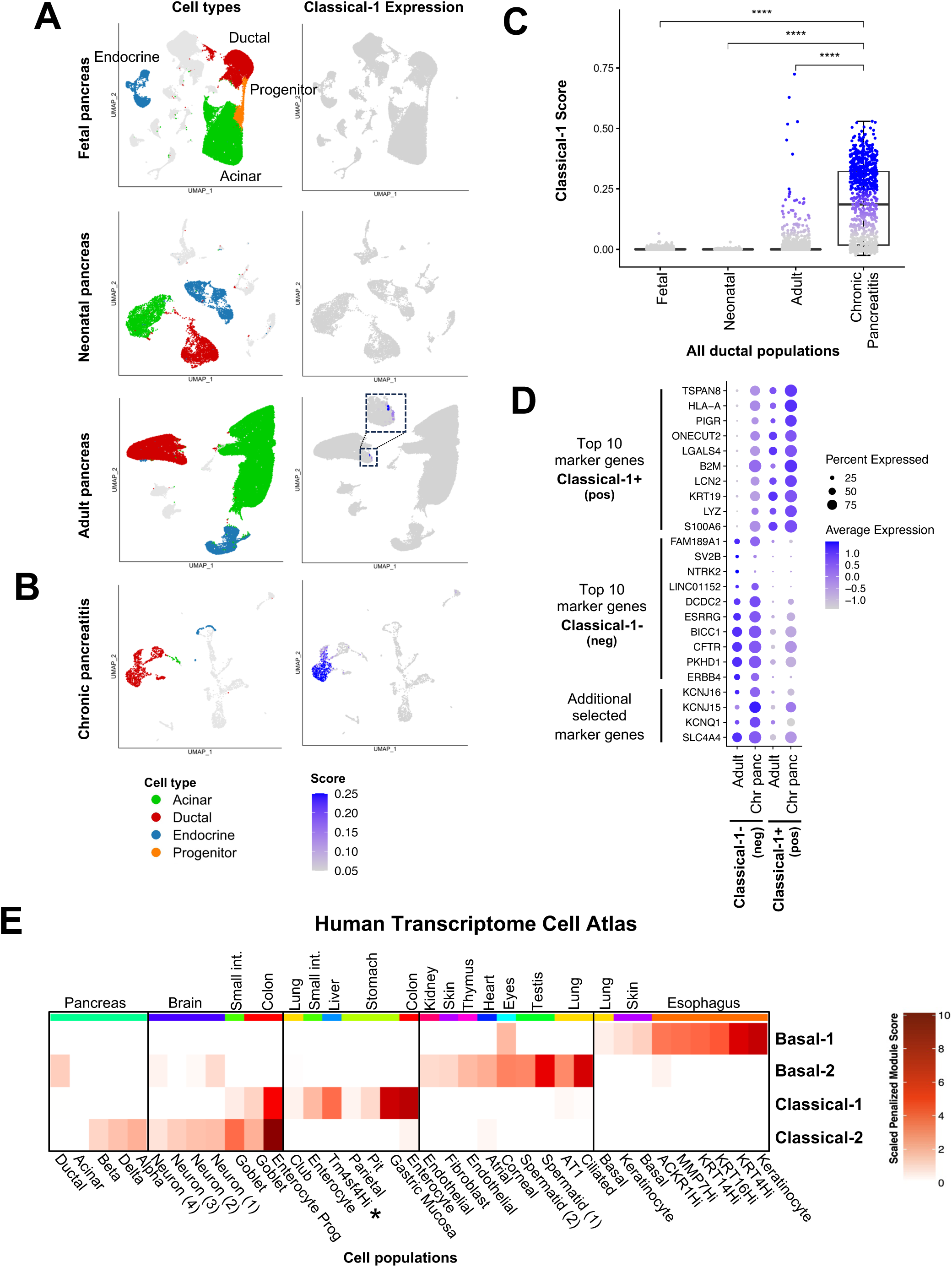
Classical-1 expression in the normal human pancreatic duct. (a) UMAP plots highlighting pancreatic cell types (left) and Classical-1 penalized module score (right) across human pancreas development timepoints. Depicted from top to bottom: fetal pancreas (from Migliorini et al.), neonatal pancreas and adult pancreas (from Tosti et al). Inset highlights rare Classical-1 program expression in the adult pancreas. Grey indicates low expression and blue indicates high expression. (b) UMAP plots highlighting pancreatic cell types (left) and Classical-1 penalized module score (right) in chronic pancreatitis (from Tosti et al.). (c) Expression of Classical-1 penalized module score across ductal populations from fetal pancreas, neonatal pancreas, adult pancreas, and chronic pancreatitis. Grey indicates low program expression and blue indicates high expression as in UMAP plots in (a) and (b). Ductal cells in chronic pancreatitis show significantly higher expression of Classical-1 compared to all other ductal populations (**** < 0.0001, Wilcoxon rank sum test). (d) Dotplot of top marker genes and selected genes across Classical-1-(neg) and Classical-1+(pos) populations in the adult pancreas and in chronic pancreatitis. Dot size indicates percentage of cells expressing each gene while dot colour indicates average expression from grey (lowest) to blue (highest). (e) Heatmap of programs scores across most enriched cell types and organ systems of the Human Transcriptome Cell Atlas dataset. Penalized module scores are utilized for scoring and are scaled across cell types. White indicates low program expression and dark red indicates high program expression.

As the Classical-1 program appeared in the context of rare adult ductal cells, we investigated its relationship to adult cell injury. We examined scRNA-seq data from patients with chronic pancreatitis. Patients with chronic pancreatitis showed marked expansion of Classical-1+ cells (**Fig. 2B-C**, **Fig. S5A**). We performed differential expression analysis between Classical-1+ and Classical-1- ductal cell populations. Notably, Classical-1+ cells showed significant downregulation of key ductal genes related to ion transport, such as *CFTR* and *SLC4A4*, as well as potassium channels such as *KCNQ1* and *KCNJ15/16* (**Fig. 2D**). Importantly, this pattern indicates that the ductal identity in Classical-1+ cells is not entirely lost with the activation of this program (**Fig S5B**). Pathway analysis of the genes of the Classical-1 program were linked to epithelial stress responses, membrane remodelling, and secretory processes consistent with an injury state (**Fig. S5C**).

Since the Classical-1 program did not strongly resemble the normal human pancreatic duct, we performed an unbiased search for this program in the human single cell atlas. Single cell RNA-seq data were obtained from the Human Transcriptome Cell Atlas (HTCA), which consists of more than 2 million single cells. Across the HTCA, the Classical-1 program was highly enriched in a number of foregut tissues including enterocytes of the colon and small intestine and gastric mucosal cells (**Fig. 2E**). These cells expressed a shared set of secretory epithelial genes (ex. *AGR2, ANXA10, CLDN18, TFF1, TFF2, CTSE, MUC5AC, MUC13, MUC17, REG4*) and conserved foregut transcription factors (ex. *GATA6*, *HNF4A*, *HNF1B*, *FOXA3;* **Fig. S6A**). Classical-1 was also enriched in TM4SF4-high epithelial cells in the liver, a population corresponding to reactive cholangiocytes in the biliary tract that emerge with chronic hepatic injury (**Fig. 2E asterisk**). These TM4SF4-high cells similarly expressed foregut secretory and mucin-associated genes, consistent with a reactive duct state rather than hepatocyte identity. These results support that the Classical-1 program represents a pan-foregut epithelial gastrointestinal injury program and is not limited to a ‘gastric’ state.

### ii. Basal-1 and Basal-2 are distinct states and not a single transcription gradient

The Basal program is often referred to as ‘strong’ and ‘weak’ and considered to be a single transcriptional continuum. However, Basal-1 and Basal-2 mapped as discrete malignant cell clusters in single-cell analysis supporting they are multiple states (**Fig. 1B**). This prompted us to further explore these programs. We first compared the gene composition of Basal-1 and Basal-2 programs. The Basal-1 program was characterized by genes associated with cornified epithelium (*IVL, SPRR1B, SPRR2A/B/D/E, S100A7/8/9, CSTA, DSG3, TGM1*) and basal cells (*MIR205HG/MIR205, ZNF750, CLCA2, LY6D, PKP1*), consistent with a basal or squamous state (**Table S5**). *TP63* and basal keratins (*KRT5, KRT6A/C, KRT14, KRT13, KRT4*) were strongly expressed genes in the Basal-1 program and confirm this is a squamous-like state. This program largely accounts for what is referred to as ‘Basal’ in the field. By contrast, the Basal-2 state was strongly enriched for genes involved in ciliogenesis (*CFAP54, CEP295NL, RFX8*; **Table S3**, **Fig. S5C**), consistent with our previous observations (Chan-Seng-Yue et al., 2020). Basal-2 tumours showed a trend toward increased acetylated tubulin staining, a structural marker of cilia, although this did not reach statistical significance (**Fig. S6B**). Neither Basal state corresponded to the normal pancreas lineage (**Fig. S5A**), which lead us to map these programs in the context of human single-cell atlas. Consistent with the above interpretation, the Basal-1 program was strongly reflected in stratified epithelial tissues including the esophagus, corneal epithelium, and basal layers of the skin (**Fig. 2E**). Whereas Basal-2 program was enriched in ciliated cells found in the lung and testis. Importantly, although both Basal-1 and -2 contribute to the original Basal signature, Basal-2 is not related to basal or squamous state.

It was unexpected that neither Basal programs were associated with EMT given the well-established relationship between these programs (Aiello et al., 2018; Bailey et al., 2016; Collisson et al., 2011; Hwang et al., 2022; Martinelli et al., 2017; Moffitt et al., 2015; Raghavan et al., 2021). Gene inspection showed *SNAI2* was upregulated in Basal-1 tumours (**Table S5**). However, *SNAI2* expression alone does not indicate EMT, as it is expressed in keratinocytes during differentiation which does not involve EMT (Kusewitt et al., 2009; Mistry et al., 2014) and is dispensable for EMT-related metastasis (Aiello et al., 2018; Fischer et al., 2015). By contrast, the Basal-2 program showed upregulation of multiple EMT- and invasion-related genes, including *AXL, TGFB2, CDH2, FN1, L1CAM, TGM2, PLAU, CAV1,* and *CCL2* (**Table S5**). EMT typically accompanies a physical change in tumour cells that cannot be observed in the transcriptome. We examined the histological features of Basal-1 and Basal-2 tumours (**Fig. 3A**; please see Flores-Figueroa et al., 2026, BioRxiv for a full analysis). Basal-1 tumours were correlated with squamous morphology (rho=0.296, p=3.04e-6), whereas Basal-2 tumours were correlated with solid tumour architecture, including cell nests, and single-cell invasion patterns (rho=0.378, p=1.46e-9). The latter patterns are consistent with a mesenchymal state. The absence of a clear EMT signal in bulk-derived RNA signatures likely reflect a technical outcome of NMF where tumour-associated EMT cannot be distinguished from leftover fibroblast program. From paired WGS, we confirmed that Basal-2 tumours were not confounded by cellularity (**Fig. S6C**; p=0.23). Although NMF ensures tumour and stromal signatures do not intermix, it does limit analysis of mechanisms that can be present in both tumours cells and stromal cells such as EMT. Given this limitation, we pursued further analysis using single cells.

**Figure 3:**
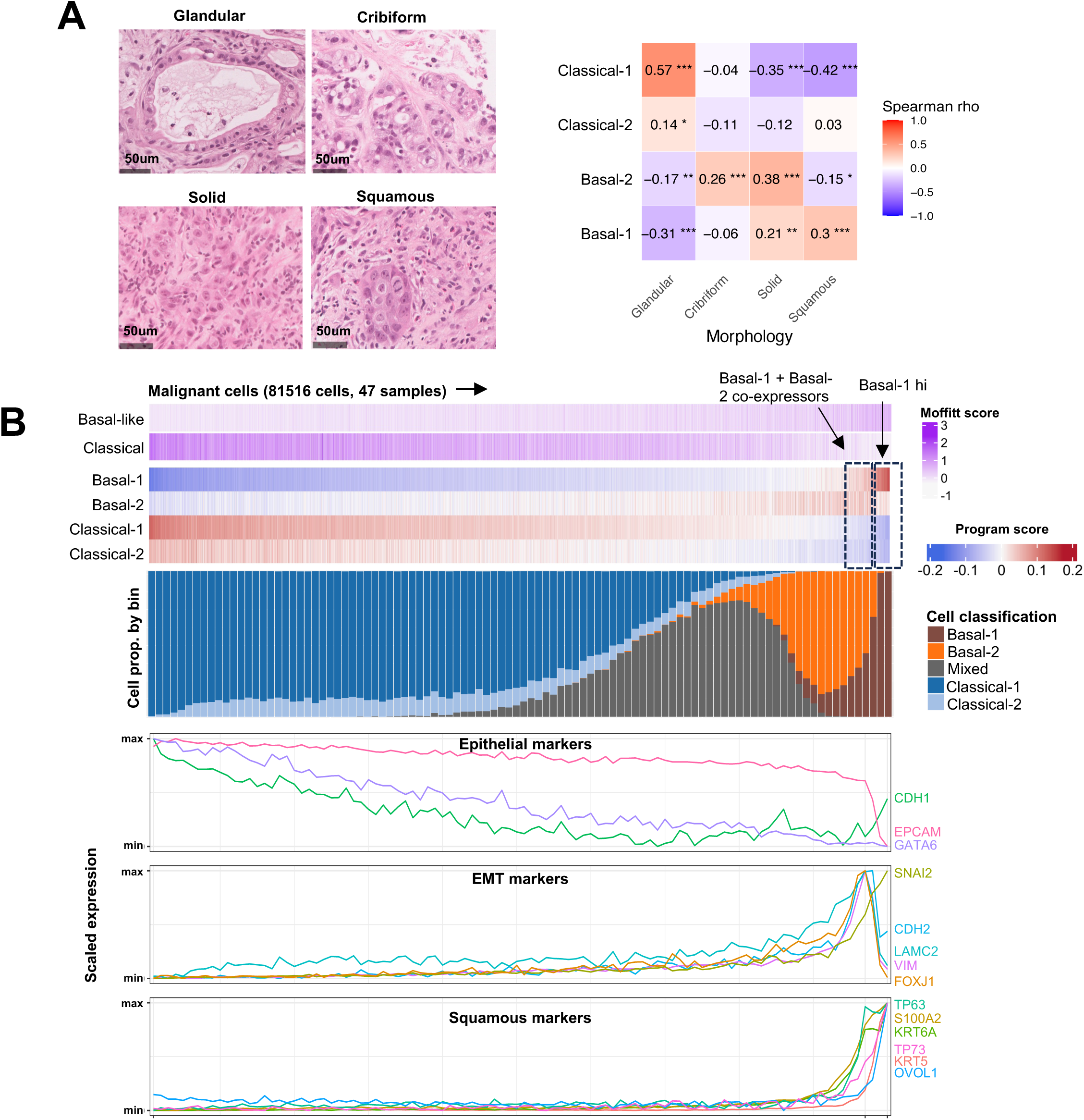
Basal states separate by morphological features and EMT state. (a) Representative histopathologic features of the four morphologic subtypes (left). Scale bars are 50µm. The glandular subtype is characterized by well-formed glands lined by pancreatobiliary-type epithelium exhibiting mild cytologic atypia. The cribriform subtype demonstrates incomplete and fused glandular structures composed of pancreatobiliary-type cells with moderate atypia. The solid subtype shows tumor cells arranged in nests or sheets with complete loss of glandular differentiation. The squamous subtype is composed of polygonal cells with abundant eosinophilic cytoplasm, prominent intercellular bridges, and dyskeratotic cells. Spearman correlation plot of morphology subtype proportion and program score for 240 matched tumours (right). * p < 0.05; ** p < 0.01; *** p < 0.001. (b) Heatmap of Moffitt and program scores of all malignant cells, excluding the neuroendocrine tumour, ordered as a program continuum along PC1 from principal component analysis (top). Bar plot of cell classification proportions across 100 bins along the program continuum (middle). Scaled expression of gene markers of duct epithelial identity, EMT, and squamous epithelium, plotted as line plots across 100 bins of the program continuum (bottom).

To resolve how EMT is related to the Basal program, we examined the scRNA-seq data. Neoplastic single cells (n=84,239) were scored for each transcriptional state and ordered using principal component analysis. In single cells, Basal-1 and Basal-2 programs co-occurred in 10.3% of neoplastic cells (**Fig. S7A**). Notably, Basal-2 frequently occurred as an independent cellular state as 39.1% of Basal-2+ cells lacked the Basal-1 program. By contrast, the majority of Basal-1 cells (10.26/11.22 or 91%) showed partial activation of the Basal-2 program supporting that Basal-2 is a more permissive state. In line with this, 36.4% of single cells co-harboured the Basal-2 and Classical-1 programs corresponding to a Hybrid state. Basal-1 and Classical-1 were mutually exclusive programs, indicating that Hybrid state originates from the Basal-2 and not the Basal-1 program. Accordingly, Basal-2 cells were enriched for the intermediate “co-expressor” program previously described by Raghavan *et al*. (**Fig. S7B**). To validate that two Basal states, we analyzed an independent dataset comprising 229 pancreatic cancer scRNA-seq samples from 12 studies (Loveless et al., 2025). Both Basal-1 and Basal-2 programs were detected in distinct malignant cell clusters further indicating they are distinct states (**Fig. S7C**). Notably, the Basal-2 program represents a key component of transcriptional plasticity in pancreatic cancer.

To characterize this further, we layered EMT signatures onto this analysis. Multiple marker genes and signatures were used. Both *SNAI2* and*VIM* were enriched in both Basal-1 or Basal-2 (**Fig. 3B**). EMT activity rose along the entire continuum but peaked in neoplastic cells that co-expressed both Basal-1 and Basal-2 programs. Notably, EMT signatures were lost once cells transitioned into a terminal Basal-1 state (Basal-1hi in **Fig. 3B and Fig. S7B**). This suggests the EMT program is most active in cells transitioning between Basal-1 and Basal-2 states. Basal-2 cells also showed elevated cell-cycle signatures, suggesting that this state is proliferative in the presence of the EMT program (**Fig. S7D**). Because KRAS signalling is closely linked to the Basal-EMT axis (Miyabayashi et al., 2020), we overlaid KRAS signalling signatures onto this framework. KRAS signalling followed a pattern similar to EMT and cell-cycle activity (**Fig. 3B and Fig. S7B**). However, KRAS signalling signatures were not lost in the terminal Basal-1 state indicating that these cells may remain KRAS-dependent. This may have implications for KRAS inhibitors that show state specific vulnerabilities (Dilly et al., 2024; Singhal et al., 2024). Using our paired WGS data, we also assessed the relationship between transcriptomic state and genetic imbalance in mutant *KRAS*. As previously described, tumours were classified as balanced (mutant=wildtype), minor imbalance (small gains 3 or less mutant alleles), or major imbalance (≥4 copies of mutant *KRAS* typically amplifications). We also separated primary and metastatic tumours as there is significant shift in mutant imbalance in metastasis (Chan-Seng-Yue et al., 2020). In primary tumours, there was no clear pattern between transcriptomic state and mutant *KRAS* imbalance. However, in metastatic disease, tumours with major KRAS imbalance were significantly enriched for Basal states (65%), predominantly Basal-2, compared to Balanced tumours (25%; p=0.002; **Fig. S7E**).

To understand the regulatory landscape underlying cell state transition, we analyzed multiome sequencing data from. Following our workflow for scRNA-seq, we identified 7872 neoplastic cells (**Fig. S8A-B**). Neoplastic cells were ordered based on the RNA profiles as above and subsequently analyzed for changes in chromatin accessibility. Overall chromatin accessibility peaked in Mixed cells (Basal-2 and Classical-1). Transition to a Basal-1 state was marked by significant reduction in chromatin accessibility (Wilcoxon rank sum test, p= 4.088e−29) supporting that this is a restricted state compared to the other programs. Motif enrichment analysis showed a distinct regulatory landscape across this continuum. Motifs related to squamous differentiation, such as *TP63*, were progressively enriched toward the Basal-1 state (**Fig. S8C**). By contrast, transition from Classical-1 to Basal-2 was marked by enrichment of a number of pathways related to epithelial regulators (*ELF3*, *RREB1*), stress and plasticity (*TEAD*, *SP1*, *MEIS*, *MAF, GLI2*) and EMT (*SNAI2*, *RUNX2*). Enrichment of *GLI2* is consistent with prior work implicating the Basal state with hedgehog signaling (Adams et al., 2019). Together, these RNA and chromatin features indicate that Basal-2 is a permissive state that enables transitions between cell states and explains both the Hybrid phenotype and the Basal continuum.

We examined whether separating the Basal states has implications for experimental models. Patient-derived organoids (PDOs) are widely used in pancreatic cancer research (Boj et al., 2015). To characterize our new signatures in these models, we performed RNA-seq on 164 PDOs (**Fig. S7F**). We first classified PDOs based on the Basal/Classical signatures, which identified 3 subgroups (Basal, Hybrid, Classical). Applying our new signatures showed that PDO models previously classified as Basal largely correspond to the Basal-1 program. Interestingly, PDOs classified as Hybrid using the Basal/Classical framework primarily derived from the Basal-2 program, indicating that this category reflects misclassification rather than a true intermediate state. Classical PDOs with a ‘weak’ signature corresponded to the Classical-2 state. Although PDOs are known to drift towards Classical in the culture (Raghavan et al., 2021), the Basal-2 program remains well represented and is often misclassified as Hybrid using previous signatures. Thus, the revised signatures improve the resolution of PDO classification and demonstrate that Basal-1 and Basal-2 represent distinct epithelial states in experimental model systems.

### iii. Classical-2 program mimics a rare neuroendocrine-like progenitor state

In 2020, we identified a second Classical program that we labelled Classical-B. However, however its overlap with the previous Classical-A program limited interpretation of this program (**Fig. S4A**). The revised Classical-2 signature addresses this problem. We first examined the gene composition of the Classical-2 program (**Table S5**). It was characterized by expression of core neuroendocrine genes (*SYP, SNAP25, PCSK1, CPE*), transcription factors central to endocrine lineage (*NKX2-2, ARX, FOXA2*), and ion channel genes related to hormone secretion (*CACNA1H, CACNB2, KCNJ11*). This expression pattern is consistent with endocrine-like state. Importantly, this program does not correspond to pancreatic endocrine cells (islets). Malignant Classical-2 cells lacked expression of all canonical islet hormones, including insulin, glucagon, pancreatic polypeptide, and somatostatin (**Fig S9A**). Also, this program expressed genes associated with epithelial injury and a secretory phenotype (*REG4*, *TFF3*, *CDX1*, *NOTUM*, *DPEP1;* **Table S5**). Pathway analysis of this program revealed enrichment for peptide hormone processing, and developmental patterning pathways (**Fig. S5**). Importantly, the gene composition of the Classical-2 and Classical-1 are highly divergent confirming these are distinct Classical states.

We used the human single cell atlas to further characterize this program. Classical-2 program was enriched in enterocyte progenitor populations in the colon, goblet cells from both the colon and small intestine, and neuronal cell populations in the brain (**Fig. 2E**). Enrichment in enterocyte progenitors is consistent with an immature lineage state compared to the differentiated gastrointestinal state that defines the Classical-1 program. The Classical-2 was significant enriched in neuronal cells. This observation prompted us to examine whether Classical-2 overlaps with the neural-like progenitor (NRP) and neuroendocrine-like (NEN) malignant programs described by Hwang *et al* (Hwang et al., 2022). Expression of both NEN and NRP were weakly positively correlated with the Classical-2 program (**Fig. S9B**). This partial concordance may reflect differences in how the programs were derived (bulk versus single cell profiling). Prior work in murine models has demonstrated that pancreatic injury and oncogenic *KRAS* signalling can induce neuroendocrine-like metaplastic states in the mouse pancreas (Ma et al., 2022; Salas-Escabillas et al., 2025). However, this program has been difficult to resolve in human pancreatic cancer. Our expanded cohort further supports a neuroendocrine-like malignant plasticity state in human tumours.

## Pancreatic cancer transcription states recur in other malignancies

Next, we sought to understand how these transcription states relate to other malignancies. We leveraged a pan-cancer single-cell atlas from Gavish *et al*. (Gavish et al., 2023) that identified 5,547 malignant transcriptional programs from 77 single-cell studies from 19 unique tumour types such as gastrointestinal, lung, breast, genitourinary, brain, head and neck, skin, bone and blood (full list in **Table S6**). Pancreatic ductal adenocarcinoma (‘PDAC’) was part of their study, which served as an internal benchmark for these analyses. Because these were single cell derived, we scored these pan-cancer malignant programs in our single cell dataset (only neoplastic cells). The heatmap in **Fig. 4A** summarizes the enrichment of these pan-cancer programs (normalized score >1.5 was threshold used). We took the following steps to ensure the observed enrichment were valid. First, individual tumours are composed of multiple malignant programs due to intratumoural heterogeneity. Thus, different malignant programs from a single cancer type may align to different transcription states. To account for this, we assessed enrichment both at the level of cancer-type and at the level of individual malignant programs within each cancer type. An example of this is shown for lung squamous cell carcinoma (LSCC) and colorectal cancer (CRC) in **Fig. S10** and further discussed below. Second, to avoid biases from unequal tumour representation in the atlas, we adjusted for the number of tumours from each cancer type. This was done using Fisher’s exact test with Bonferroni correction (adjusted alpha=1.35e-3). Third, to ensure that the enrichment analysis was not from single genes due to amplification biases in scRNA-seq, the top-ranking genes (typically 3–20 per program, depending on program size and gene overlap) were projected as a dot plot (**Fig. S10**). This was followed by pseudo-bulk differential expression analysis to gene-level p-values which were subsequently combined using the Cauchy method (Liu and Xie, 2020) (**Fig. S10**).

**Figure 4:**
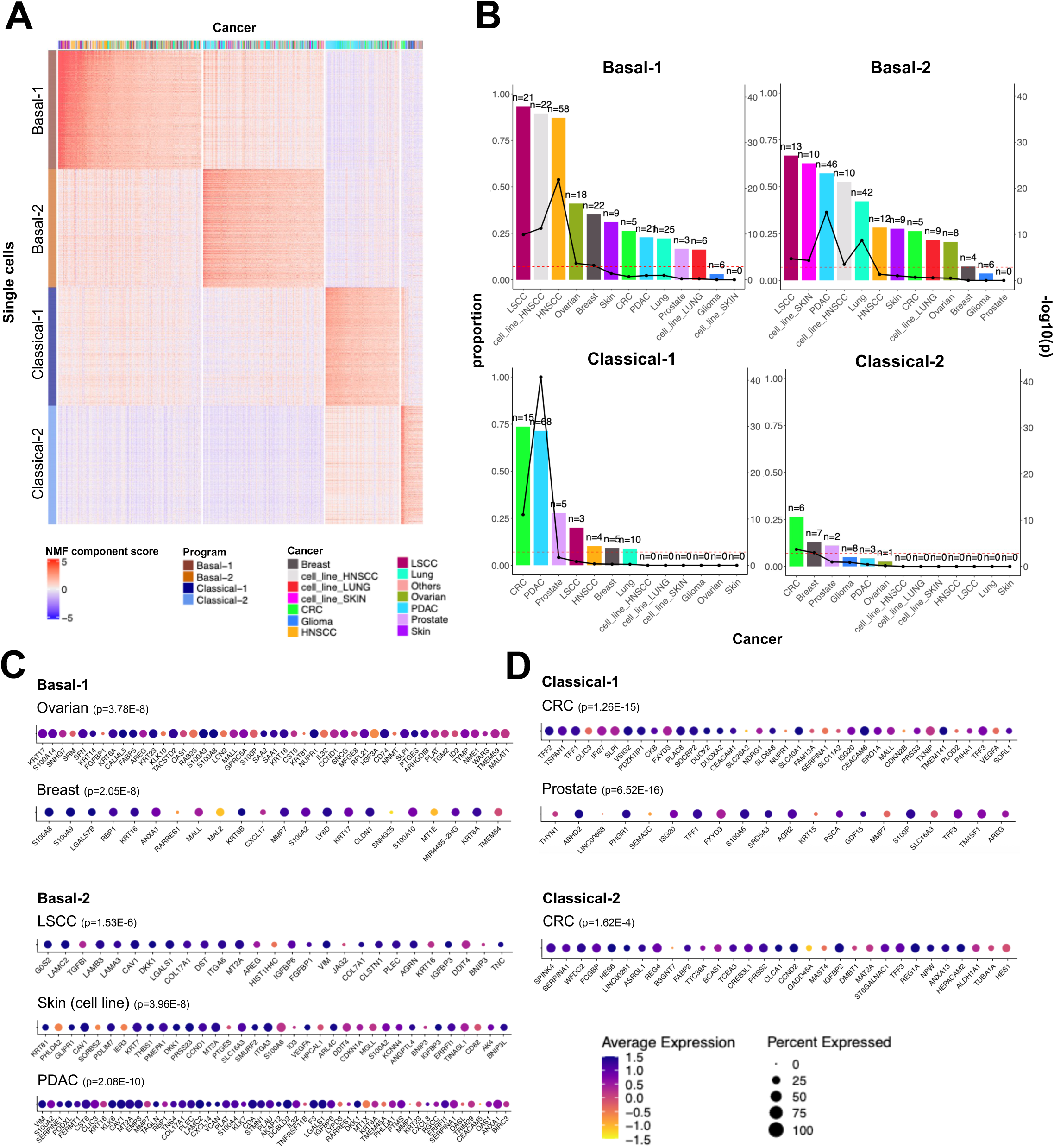
Pancreatic cancer transcriptional programs in the context of pan-cancer. (a) Heatmap showing pan-cancer malignant transcription program scores in the top 500 cells within each pancreatic cancer state (Basal-1, Basal-2, Classical-1, Classical-2) of the single cell cohort. HNSCC head and neck squamous cell carcinoma; CRC colorectal cancer; LSCC lung squamous cell carcinoma; PDAC pancreatic ductal adenocarcinoma. (b) Combined bar and line plots of cancer sample percentage (left-side y-axis) and significance of enrichment (right-side y-axis) for each cancer type for Basal-1, Basal-2, Classical-1, and Classical-2. Bar plots show the percentage of tumour samples from a given cancer type harboring malignant programs aligned with each pancreatic cancer state in (a). The number of malignant programs from each cancer type aligned with each pancreatic state is shown on top of each bar. Line plots show the - log10(p-value) for the cancer type enrichment within each pancreatic cancer state. P-values were determined by one-sided Fisher’s exact test with Bonferroni correction. (c) Dot plots of top-ranking genes for ovarian and breast cancers aligned with Basal-1 state (top). Dot plots of top-ranking genes for LSCC, skin (cell line) and pancreatic ductal adenocarcinoma (PDAC) aligned with Basal-2 state (bottom). Dot size indicates percentage of cells expressing the genes while dot colour indicates average expression from yellow (lowest) to blue (highest). P-values were determined from gene-level differential expression p-values (Wilcoxon rank sum test) combined using the Cauchy method. (d) Dot plots of top-ranking genes for colorectal cancer (CRC) and prostate cancers aligned with Classical-1 state (top). Dot plot of top-ranking genes for CRC aligned with Classical-2 state (bottom)

The Basal-1 state showed strongest enrichment for malignant programs from LSCC (p=1.39e-10, Fisher’s exact test with Bonferroni correction) and head and neck squamous cell carcinoma (HNSCC) (p=1.25e-22) (**Fig. 4B**). Unexpectedly, we also observed enrichment in ovarian (p=2.55e-4) and breast cancers (p=7.03e-4). Inspection at the gene level showed that the ovarian and breast cancer programs were driven by canonical squamous and basal epithelial cell markers (ex. *KRT6A/B, KRT14, KRT17, S100A2/8/9*; **Fig. 4C , top**). This is consistent with reports that show rare subsets of high-grade serous ovarian cancer exhibit squamoid differentiation (Sun et al., 2021; Tamura et al., 2024). In breast cancer, the enrichment align with the basal-like phenotype associated with triple-negative disease (Foulkes et al., 2010; Perou et al., 2000). The Basal-2 state showed strong enrichment for malignant programs derived from LSCC (p=1.98e-5), Skin (p=4.52e-5) PDAC (p=1.65e-15), and HNSCC (p=3.26e-4) (**Fig. 4B**). Unlike Basal-1, this enrichment was not driven by squamous or basal cell genes but instead by EMT- and invasion-associated programs (**Fig. S10**, top; *VIM, LAMC2, TGFB1*) (**Fig. 4C**, bottom). Notably, enrichment of Skin, PDAC and HNSCC malignant programs in Basal-2 single cells was consistently driven by EMT-associated genes (**Fig. 4C**, bottom). This indicates that Basal-2 represents a conserved EMT program shared across other cancers rather than a tissue specific lineage state. Together, these data further support that the two Basal programs are biologically distinct.

Classical-1 cells showed a striking enrichment in malignant programs from CRC (p=1.12e-11) and PDAC (p=2.09e-41) (**Fig. 4B**). The enrichment for CRC was represented by core Classical-1 genes (*TFF1*, *TFF3*, *AGR2*, and *MMP7*; **Fig. 4D , top**). This continues to show that Classical-1 program is not simply a gastric state. Prostate cancer malignant program trended towards enrichment in Classical-1 cells (p=1.97e-2). Although this did not meet our statistical threshold (p=1.35e-3), the enrichment was driven by canonical Classical-1 genes (**Fig. 4D , top**; e.g., *AGR2, TFF1, TFF3*). We mention this here because it aligns to reports showing that a third of castration-resistant prostate cancers acquire a gastrointestinal lineage state due to activation of key foregut transcription factors such as *HNF4G* and *HNF1A* (Shukla et al., 2017). Together, this supports the interpretation that the Classical-1 program is not a pancreatic lineage state but a conserved gastrointestinal epithelial program recurrent in other malignancies.

Classical-2 showed the lowest enrichment in the pan-cancer atlas. This is likely because neuroendocrine tumours (NETs) are underrepresented in the pan-cancer atlas. Nevertheless, we did observe a significant enrichment of CRC malignant programs (p = 2.11e-4; **Fig. 4B**). Importantly, this CRC program was distinct from the CRC program enriched in Classical-1 cells (**Fig. 4E-F**). At the gene level, this CRC program was composed of genes related to intestinal and enterocyte differentiation (*REG4*, *TFF3*, *SPINK4*, *REG1A*, *FABP2*, *CLCA1*, *ANXA13*, *FCGBP*, *SERPINA1*, *DMBT1*). Unlike Classical-1, PDAC malignant programs did not strongly map to Classical-2 cells and show that this distinct state from the Classical-1 program. Collectively, these analyses show that pancreatic cancer transcriptional states are broadly observed as malignant lineage programs in a diverse range of cancers.

## Transcriptional plasticity and genomic evolution are largely uncoupled during tumour progression

As pancreatic cancer is characterized by extensive chromosomal instability (Mullen et al., 2025; Notta et al., 2016), we investigated how genomic evolution impacts transcriptional dynamics. We first quantified transcriptional plasticity in individual tumours by measuring the degree of intratumoural heterogeneity in our scRNA-seq dataset. To do this, we calculated the proportion of cells belonging to each transcriptional state. We then clustered tumours based on their cell state composition. Using this approach, we identified four subgroups (**Fig. 5A-C**): (1) tumours dominated by Basal-1 and Basal-2 cell states (“Basal-dominant”; n = 4); (2) tumours with extensive heterogeneity harbouring all transcription states (“High diversity”; n = 8); (3) tumours with intermediate heterogeneity (“Moderate diversity”; n = 9); and (4) tumours dominated by Classical-1 and Classical-2 states (“Classical-dominant”; n = 23). The Classical-dominant group was significantly enriched in primary tumours, whereas metastatic tumours were enriched in Moderate/High and Basal-dominant subgroups (**Fig. S11A**; p = 0.0199, Fisher’s exact test).

**Figure 5:**
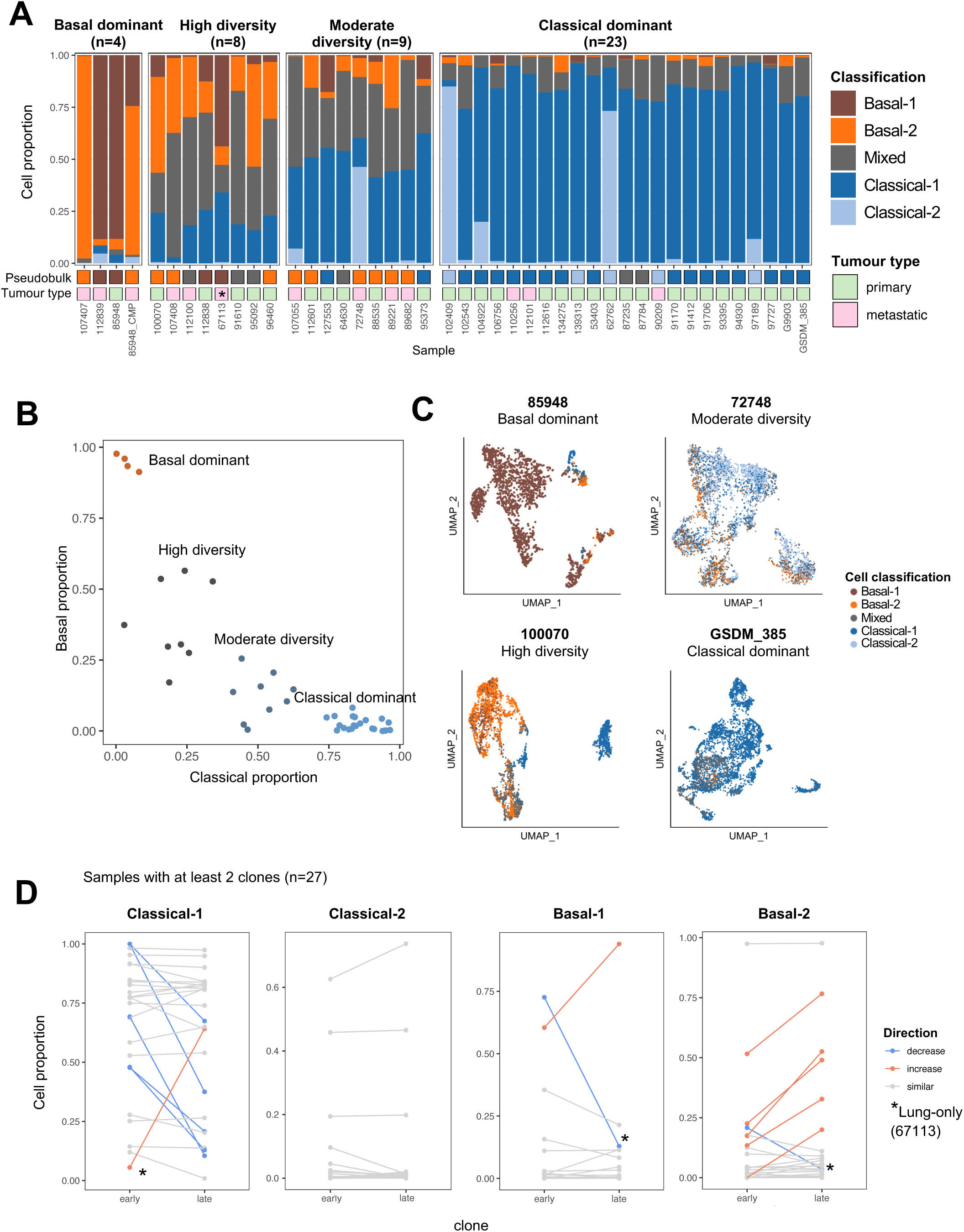
Characterization of single cell transcriptomic and genomic variation. (a) Classification proportions of malignant cells from each sample across the scRNA-seq cohort, separated into diversity groups. Each bar represents a sample and is annotated with the pseudobulk classification and tumour type as primary (pink) or metastatic (green). An asterisk (*) highlights the one lung metastasis. Samples with fewer than 50 malignant cells were omitted, as well as the neuroendocrine tumour. (b) Scatterplot of Basal and Classical program proportions across the malignant cells of the samples of the scRNA-seq cohort, coloured and labelled by k-means diversity grouping. Each dot represents a sample with at least 50 malignant cells as shown in (a). (c) UMAP plots of representative samples from each subgroup, cells coloured by cell classification. (d) Classical-1, Classical-2, Basal-1 and Basal-2 cell proportion between early and late clones in samples with at least 2 clones (n=27). Samples are connected by a line, and lines are coloured by direction of change in proportion, where “similar” in grey indicates a change of less than 10% of total proportion. Asterisks (*) highlight the one lung metastasis.

Next, we examined the relationship between transcriptional heterogeneity and genomic instability. Copy number burden and the number of genetic clones were inferred from scRNA-seq (as shown in **Fig. S1C**) and were not found to be significantly different across the transcriptional subgroups above (**Fig. S11B**). Primary tumours showed a trend toward greater clonal diversity than metastatic tumours (**Fig. S11C**). To validate this, we analyzed our WGS dataset. Consistent with the scRNA-seq results, primary tumours exhibited significantly greater clonal diversity than metastatic tumours (**Fig S11D**), which reflects the genetic bottleneck of metastatic dissemination (a detailed analysis is presented in Fang et al., 2025, BioRxiv), and is consistent with previous work (Makohon-Moore et al., 2017). Importantly, the copy number inference from scRNA-seq is concordant with WGS and confirms that scRNA-seq can capture the genetic architecture of tumours. Overall, this supports that genetic diversity decreases during tumour progression, whereas transcriptional diversity increases.

We then analyzed transcriptional dynamics of individual genetic clones. Among the 47 patients profiled by scRNA-seq, 27 contained at least two genetically distinct clones. Within individual tumours, clones were ordered according to their copy number burden with clones harbouring fewer alterations labelled ‘early’ and those with more alterations labelled ‘late’. In most tumours (78%, n = 21), the transcriptional state of early and late clones remained the same despite ongoing evolution (**Fig. 5D (grey) and Fig. S12A**). Consistent with this, we did not identify any recurrent genomic alteration that associated with a particular transcriptional program. In a minority of cases, (22%, n = 6), transcriptional states shifted during clonal evolution. For example, patient 100070 harboured five genetic clones (C1-C5; **Fig. S12B-D**). C1 harboured a near-diploid genome resembling a normal cell. This clone harboured a small number of copy number alterations, including loss of heterozygosity at chr17p containing *TP53* (**Fig. S12D inset**), which is consistent with an early stage of pancreatic cancer evolution. The transcriptional state of C1 was Classical-1. The subsequent aneuploid clones (C2-C5) diverged into two transcriptional lineages: one that maintained Classical-1 program (C2), while the other showed increased proportions of Mixed, Basal-1 and Basal-2 states (C3-C5). This indicates that Basal programs can emerge clonally from an earlier Classical-1 precursor. In 5 of 6 patients, where the transcription state shifted during clonal evolution, the transition was from Classical-1 to Basal-2. These findings are consistent with phylogenetic analyses of pancreatic cancer (Hayashi et al., 2020) and with recent evidence that EMT-associated programs are linked to genomic instability in this disease (Perelli et al., 2025). However, there was one exception where a late clone shifted to Classical-1 state (**Fig. 5D**, asterisk). Interestingly, this was from a patient with lung-only metastasis, a rare clinical presentation associated with favourable prognosis. Overall, these data show that transcriptional states are largely maintained during clonal evolution. Importantly, this supports a model in which transcriptional plasticity and genomic evolution represent largely independent axes of tumour progression.

## Classical-1 and Classical-2 states emerge as part of injury response in mouse models

The above observations raised questions regarding how the transcriptional programs arise as they are mostly uncoupled from genomic evolution. Because epithelial cell plasticity can emerge from tissue injury and oncogenic stress, we examined whether these human transcriptional states reflect conserved epithelial reprogramming events seen in mouse models. We first analyzed a well-established cerulein injury model of pancreatic cancer. In this model, injury alone is sufficient to generate a spectrum of metaplastic epithelial states, including pyloric-like mucinous cells, tuft-like cells, and enteroendocrine-like populations (Ma et al., 2022). Projection of the human transcriptional programs onto this mouse single cell data showed enrichment of the Classical-1 and Classical-2 programs (**Fig. 6A** and **Fig. S13A**). Basal-1 and Basal-2 were not detected in the injury model. Classical-1 program mapped to cells denoted as ‘Ductal/Mucin’ and described as pyloric-like by the authors (Ma et al., 2022). By contrast, the Classical-2 program aligned with enteroendocrine-like (EEC) epithelial cells consistent with its neuroendocrine features. This mouse dataset was generating using lineage tracing (*Ptf1a-CreER*; *Rosa LSL-EYFP*) and demonstrate that both Classical states are related to the exocrine lineage. Importantly, this supports that the Classical-2 program is not an artefact of contaminating endocrine cells but reflects an injury state from exocrine tissue. Thus, the human Classical states mirror conserved injury responses in mouse models and align to our scRNA-seq analysis of chronic pancreatitis patients.

**Figure 6:**
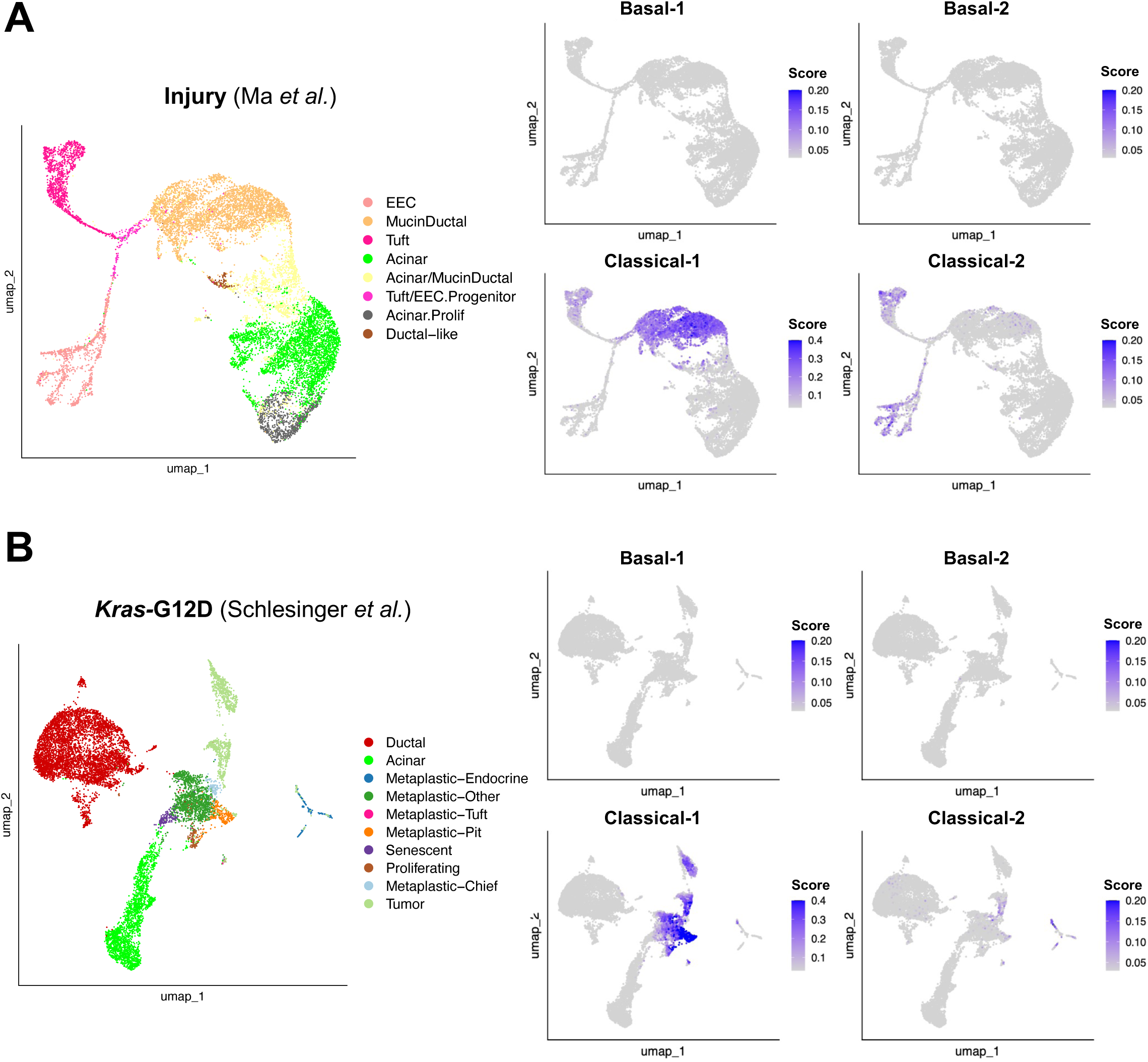
Expression of Classical-1 and Classical-2 programs in early mouse models. (a) UMAP plots of cell types (left) and programs scores (right) across populations in the pancreatitis *Kras*-WT dataset from Ma et al. Converted human-mouse orthologs were scored using the penalized module score, where grey indicates low expression of the program and blue indicates high expression of the program. (b) UMAP plots of cell types (left) and programs scores (right) across populations in the *Kras*-G12D mutant dataset from Schlesinger et al. Converted human-mouse orthologs were scored using the penalized module score, where grey indicates low expression of the program and blue indicates high expression of the program.

We next examined these programs in the context of oncogenic stress. Similar to the above analysis, oncogenic *Kras* activation induces a spectrum of metaplastic states such as duct-like, gastric-like, tuft-like, neuroendocrine-like, and senescent populations. These pre-malignant states arise from chromatin remodelling events (Alonso-Curbelo et al., 2021; Schlesinger et al., 2020). We overlaid our human transcription states onto mouse scRNA-seq data from the inducible *Kras*-G12D model (Ptf1a-CreER; *Kras*-G12D-tdTomato) (**Fig. 6B** and **Fig. S13B**). The Classical-1 program was detected in gastric pit- and chief-like epithelial populations, whereas Classical-2 was enriched in neuroendocrine-like epithelial cells. Similar to the injury model, Basal-1 and Basal-2 were not detected among these metaplastic populations. These analyses suggest that pancreatic cancer transcriptional states arise from epithelial plasticity programs related to cell injury and oncogenic stress.

## Classical-1 program initiates before onset of aneuploidy in human tumours

The above analyses were derived from mouse models. Next, we investigated when the Classical transcriptional states arise in human tumours. To do this, we leveraged our scRNA-seq, which enables tracking of both transcriptional states and aneuploidy in single cells. Because aneuploidy marks malignant transformation, distinguishing neoplastic from non-neoplastic epithelial cells provides an opportunity to examine when the transcription states arise relative to aneuploidy. In our scRNA-seq dataset, 84,239 of 161,363 cells (n=48 patients) were classified as neoplastic based on aneuploidy and lineage markers (**Fig. 7A** and **Fig. S1B**). We next focused on the remaining 77,124 non-neoplastic cells. Of these, 14,345 cells (19%) were identified as epithelial based on lineage marker expression and were composed of acinar, ductal, and endocrine populations (**Fig. 7B** and **Fig. S1B**). We compared transcriptional states of neoplastic and non-neoplastic epithelial cells.

**Figure 7:**
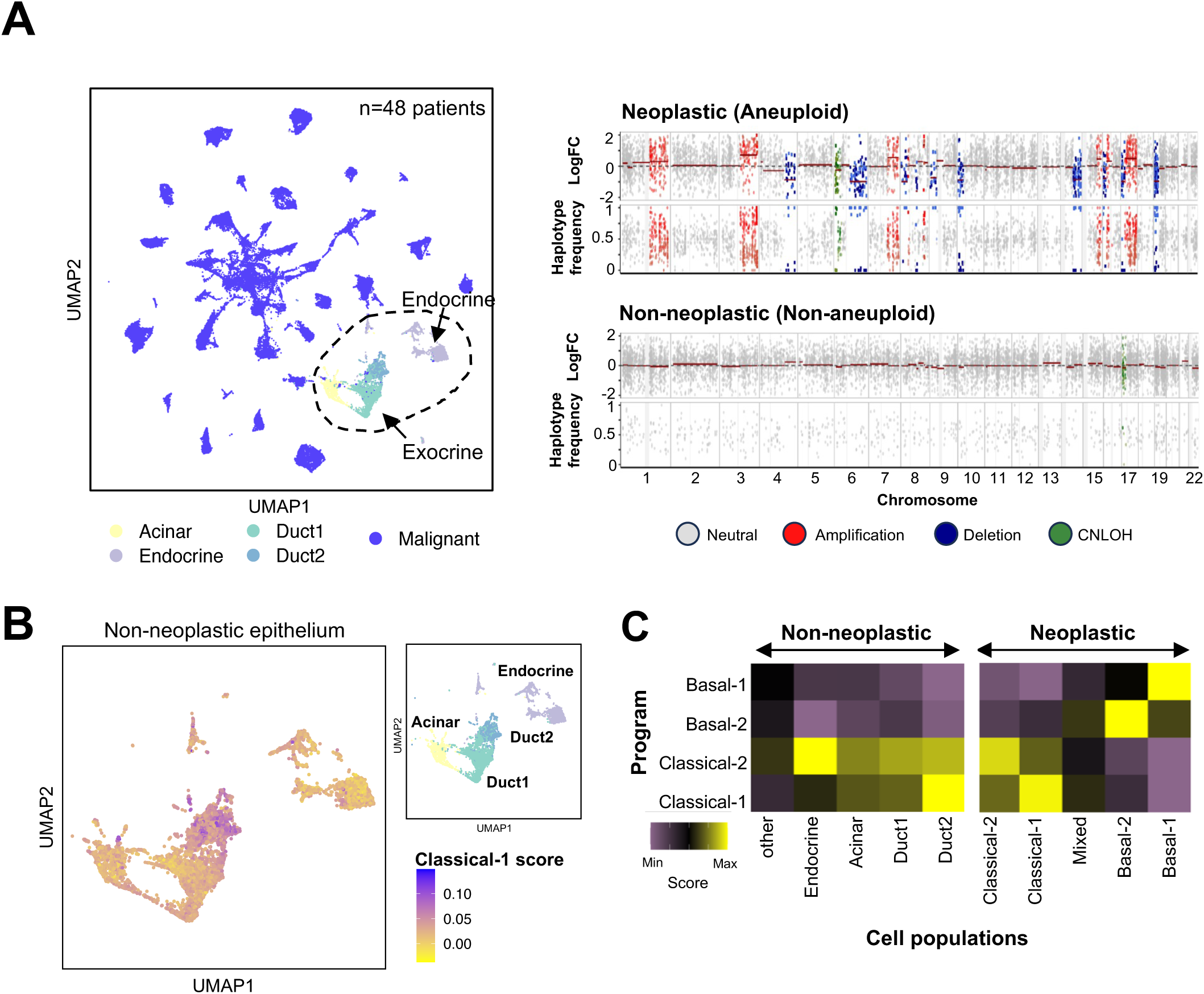
Tumour programs emerge at different times relative to the onset of aneuploidy. (a) UMAP plot of all pancreatic epithelial lineage cells coloured by cell type. Acinar, Endocrine, Duct1 and Duct2 represent non-neoplastic stromal cells (left). Malignant cells represent neoplastic tumour cells. Haplotype-aware inferred copy number plot illustrating differences in RNA log fold change (“LogFC”) and haplotype (allele) frequency between representative neoplastic and non-neoplastic populations across the genome (right). Regions are coloured by predicted copy number status. (b) Subset of UMAP plot highlighting the Acinar, Endocrine, and Duct1 and Duct2 populations. UMAP plots show Classical-1 score (left) and cell types (right). (c) Mean program score across non-neoplastic and neoplastic cell populations, scaled by program. Non-neoplastic populations are separated by cell type, with “other” indicating non- epithelial lineage cells such as fibroblasts and immune cells. Neoplastic populations are separated by cell classification.

Basal-1 and Basal-2 programs were almost exclusively detected in neoplastic cells, consistent with their association with progression (**Fig. 7C**). However, there was one exception. In one case (primary tumour 97727), we identified a rare population of epithelial cells with activation of a basal keratin program and no evidence of aneuploidy (**Fig. S14A-B**). These rare cells expressed basal cell markers including *TP63, KRT5/6A, KRT14, KRT17,* and *S100A2* (**Fig. S14C**). This is consistent with report of rare delta TP63-positive basal cells in the human pancreas and associated with injury (Martens et al., 2022). Outside of this case, Basal-1 and Basal-2 programs were overwhelmingly neoplastic cells indicating they are associated with transformation (**Fig. 7C**). The Classical-2 program was detected in both neoplastic and non-neoplastic epithelial compartments. However, gene-level inspection showed that the signal in non-neoplastic cells was driven by the beta cell program (*INS, PCSK1, PCSK2, CPE, PAX6, NEUROD1, NKX2-2*) and did not reflect the Classical-2 state found in tumours (**Fig. S15A**). As the Classical-2 program is rare, we could not identify a non-neoplastic precursor associated with this program.

The Classical-1 program was detected in both non-neoplastic and neoplastic epithelial cells. Within the non-neoplastic compartment, we identified two ductal populations that expressed canonical duct markers (**Fig S3 and Table S5**), which we annotated ‘Duct1’ and ‘Duct2’ (**Fig. 7B**). Notably, Duct2 cells were strongly enriched for the Classical-1 program comparable to the level in Classical-1 neoplastic cells (**Fig. 7C**). To define the identity of Duct1 and Duct2 cell populations, we performed differential expression analysis (**Fig. S15B**). Duct1 cells retained strong expression of typical pancreatic duct markers such as *CFTR*, *SLC4A4*, and *DCDC2.* By contrast, Duct2 cells showed significant reduction in these ductal identity markers indicating partial loss of ductal identity. Differential expression showed that Duct2 had activated gastrointestinal markers (*MUC5B*, *TFF1*, *TFF3*, *AGR2* and *CRISP3*) and independently recapitulated the Classical-1 program (**Fig. S15B**). This phenotype mirrored the injury-associated epithelial state observed in chronic pancreatitis. Consistent with this, the Duct2 program was strongly enriched in chronic pancreatitis patients (**Fig. S15B**). Together, these data indicate that Duct2 represents an injury-associated state rather than a normal epithelial population. The detection of these Duct2 cells provided us an opportunity to investigate the cellular origins of the Classical-1 program in non-neoplastic human cells.

We sought to validate if the program of Duct2 cells can be found in an independent dataset. To do this, we analyzed a spatial transcriptomics dataset generated by Carpenter *et al*. (Carpenter et al., 2023) that included both pancreatic cancer specimens and normal donor pancreas. Using the GeoMx NanoString platform, morphologically defined regions of interest such as acinar tissue, normal ducts, ADM, PanIN, and tumour regions, were profiled. Notably, the Duct2 program was detected in normal ducts from both cancer-free donors and tumour-adjacent tissue. PanIN lesions also showed consistent enrichment of the Duct2 program, whereas ADM regions were more heterogeneous (**Fig. S16**). Together, these findings indicate that Duct2 cells are present in normal human pancreas and not just associated with inflammatory changes in tumours. Overall, these findings indicate that the Classical-1 program, the main transcriptional state of pancreatic cancer, originates as a pre-malignant state prior to the onset of aneuploidy.

## Classical-1 program arises prior to *KRAS* mutation

Given that the Classical-1 program arises before aneuploidy, defining when it emerges relative to *KRAS* mutation is critical. Addressing this requires genotype and phenotype of the same cells. Standard droplet-based scRNA-seq (10x Genomics) lack sufficient coverage at the 5’ end of *KRAS*, where hotspot mutations reside. To overcome this, we adapted a genotyping of transcriptomes protocol by Landau and colleagues (Nam et al., 2019) to amplify exon 2 of *KRAS* (G12X) from full-length scRNA-seq cDNA. This enabled assignment of *KRAS* mutation status to individual cells via cell barcodes. This approach was applied to 16 donors, yielding *KRAS* mutation hotspot coverage in 22,625 cells (**Fig. 8A**). In cases with matched tumour WGS, the detected *KRAS* mutant alleles were fully concordant (**Fig. 8C**). Using this assay, *KRAS* mutations were readily detected in neoplastic cells (**Fig. 8B**). However, *KRAS* mutations were absent from Duct2 cells. We closely examined individual donors with abundant neoplastic and non-neoplastic Duct2 cells and found that Duct2 precursor cells did not carry the *KRAS* mutations despite being from the same individual (**Fig. 8D**). This supports that the Classical-1 program is absent in Duct2 cells and is not driven by oncogenic *KRAS* mutations. Moreover, this indicates that Duct2 cells could not be derived from PanIN lesions, which uniformly harbour *KRAS* mutations (also covered later).

**Figure 8:**
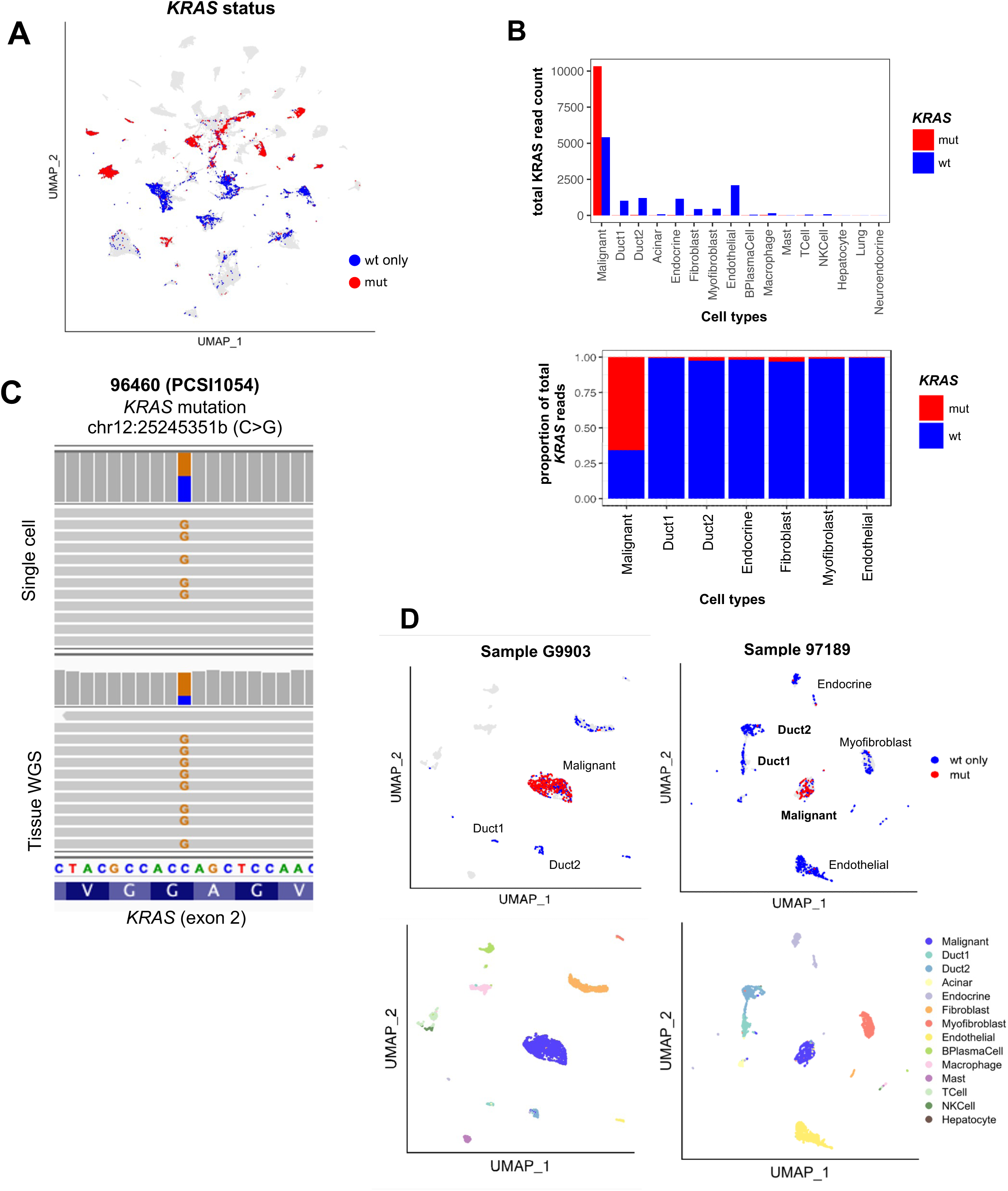
*KRAS* mutation status across tumour and stromal populations. (a) UMAP plot of all cells in the scRNA-seq cohort coloured by KRAS mutation status, where red indicates at least one mutation read found (mut) and blue indicates only wildtype reads (wt only). Cells with unknown *KRAS* status due to lack of sequencing coverage are shown in light grey. (b) Bar plot of total *KRAS* mutation (mut, red) or wildtype (wt, blue) read count by cell type across all *KRAS* PCR amplified samples (top). Proportion of mutant vs wildtype *KRAS* reads found in selected cell types (bottom). Only cell types with at least 250 *KRAS* mutant or wildtype reads are included. (c) Comparison of mutation found between matched bulk RNA-seq and single cell *KRAS* PCR amplified reads from patient 96460 (PCSI1054) in IGV. (d) UMAP plots of all cells in samples from patients G9903 (left) and 97189 (right) coloured by *KRAS* mutation status (top) and cell type (bottom).

The early emergence of some transcription programs could influence mutation trajectory of tumours. We did a side-by-side comparison of the transcription states emerging in the cerulein injury and Kras-*G12D* mouse models. The *Kras-G12D* model showed marked skewing towards the Classical-1 program, whereas the cerulein injury model was enriched for Classical-2 (Fisher’s exact test, p = 6.55e-68; **Fig. S17A–B**). We examined this relationship in human tumours. Among Classical-1 tumours, 96% harboured *KRAS* mutations (182/189), with G12D being the most frequent allele (**Fig. S17C**). By contrast, only 66% Classical-2 tumours harboured *KRAS* mutations (33/50), and G12D was present in only 18% of cases (9/50; Chi-squared test, p=2.38e-18). Notably, 44% (22/50) of Classical-2 tumours were *KRAS* wildtype. The lack of *KRAS* mutations in the Classical-2 was not due to cellularity concerns (**Fig. S6D**) and to ensure this was not the case, we also performed a manual review of all *KRAS* wildtype genomes. Overall, these data show that the *KRAS* allele status is significantly different between Classical-1 and Classical-2 tumours (Chi-squared test, p=9.01e-15). This remains an association but support a model where early changes in state may influence mutational trajectory of tumours.

## Cellular origins of the Classical-1 program

Analysis of the Carpenter *et al*. data showed that the Duct2/Classical-1 program can be detected in multiple non-neoplastic tissue contexts, including normal ducts, PanIN, and variably in ADM. However, those data did not allow us to define the cellular identity of this transcriptional state. To investigate this *in situ*, we leveraged a large-scale imaging mass cytometry (IMC) dataset (Shakfa et al., 2025, BioRxiv). This study profiled immune-stromal interactions in pancreatic cancer using an antibody panel that can distinguish Basal and Classical tumour cells. This study focuses on the broad Basal and Classical states but several markers in the IMC panel map to the transcription programs used in this study. Basal markers such as TP63 (Basal-1) and KRT5 (Basal-1), and CAV1 (Basal-2) captured both Basal states. Classical markers such as GATA6, CLDN3, AGR2, and TFF1 correspond to the Classical-1 program. Markers of the Classical-2 program were not defined at the time of study design. The IMC panel was applied to serial sections of a tissue microarray (TMA) derived from 221 resected patients (∼4 cores per case), yielding over 800 multiplexed images spanning tumour and adjacent tissue. Images were segmented into more than 10 million single cells to enable high-resolution spatial characterization of epithelial cell states in intact tissue. This protein-based approach also serves to validate our RNA-based states.

As part of validation of this approach, we first examined whether two Basal states exist spatially. Among cells classified as Basal in the TMA, one population expressed markers of the Basal-1 program (KRT5+TP63+), whereas a second population lacked these markers but strongly expressed CAV1, a Basal-2 marker (**Fig. 9A**). CAV1 is a membrane scaffolding protein associated with EMT and invasion (Goetz et al., 2011). TP63+KRT5+CAV1- cells comprised 23% of Basal cells, whereas CAV1+TP63-KRT5- cells comprised 20% in this TMA cohort. The remaining cells were heterogenous for Basal markers (ex. S100A2). This confirms that Basal-1 and Basal-2 are distinct Basal states.

**Figure 9:**
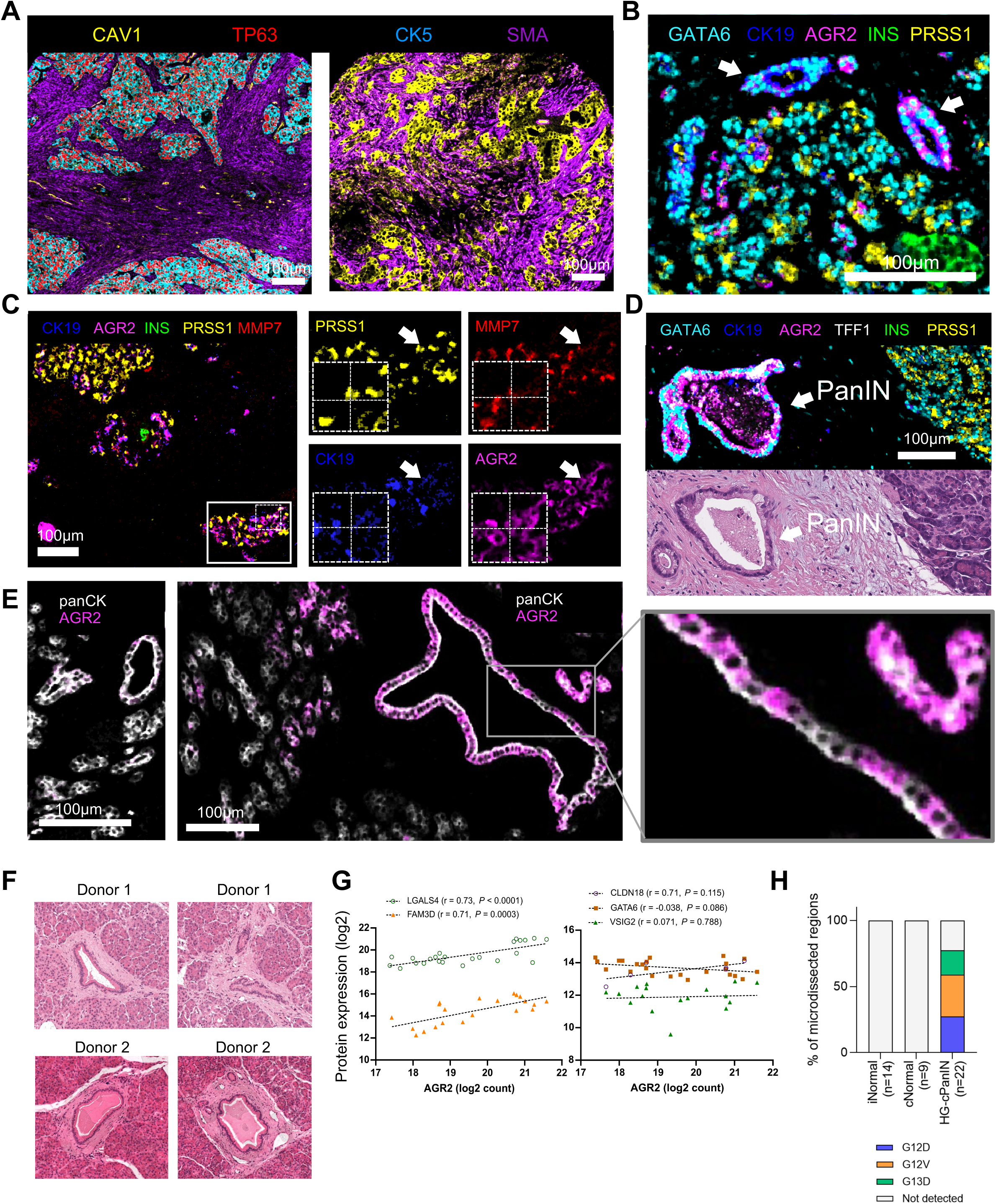
Spatial characterization of Classical-1+ epithelial cells. (a) Imaging mass cytometry (IMC) of pancreatic tumours identifying two spatially distinct Basal populations. Basal-1 cells express TP63 and KRT5 (left panel), whereas Basal-2 cells lack these markers but express CAV1 (right panel). SMA expression highlights fibroblasts. (b) IMC of tumour-adjacent regions showing interlobular ducts (CK19+) with and without expression of AGR2 expression (see arrows). (c) Representative ADM regions showing expression of CK19, PRSS1, AGR2 and MMP7 (left panel). Zoomed in region of ADM (right panel). (d) Co-expression of multiple Classical-1 markers (AGR2, GATA6, TFF1) in PanIN. (e) Example of AGR2 negative ductal cells (left) as a control (left panel). Representative image of a large duct showing mosaic expression of AGR2 (middle panel). Zoomed in inset of middle panel (right panel). All scale bars are 100µm. (f) Images of normal ductal structures from cancer-free donors that were dissected for Deep visual proteomics (DVP). (g) Correlation plots of Classical-1 associated proteins (LGALS4, FAM3D, CLDN18, GATA6, and VSIG2) with AGR2 expression in normal ducts from cancer-free donors. (h) Detection of KRAS mutant peptides in microdissected regions.

Next, we investigated Classical-1+ cells. We focused on tumour-adjacent regions in the TMA that retained normal pancreas architecture, as these would be enriched in non-neoplastic cells. Eighty-four TMA cores met these criteria. Within these regions, the majority of cells were classified as acinar (∼45%), islet (∼20%), and ductal (∼12.5%) by morphology and markers (**Fig. S18A-B**). Unsupervised clustering based on protein markers yielded 30 cell clusters that collapsed into subsets of acinar, ductal, endocrine, stromal, and residual tumour cells. Tumour cells in adjacent regions represented <10% of cells and could be readily identified (**Fig. S18C**). Within cell clusters enriched for duct cells, two populations were evident (**Fig. S18B;** purple and pink categories). These two ductal cell populations expressed similar levels of core epithelial and pancreatic lineage markers (ECAD, CLDN3, PDX1, FOXA2, KRT19; **Fig. S18D**) but separated based on AGR2 expression. AGR2 is a core gene of the Classical-1 program and was also markedly upregulated in Classical-1+ cells in the Carpenter *et al*. dataset. AGR2+ cells also upregulated MMP7 found in Duct2 cells, and co-expressed GATA6, a transcription factor of the Classical program (**Fig. 9B** and **Fig. S18D**) (Martinelli et al., 2017). AGR2+ ductal cells accounted for 7.5% of cells within tumour adjacent regions (**Fig. S18A**).

We examined the spatial organization of these AGR2+ cells. AGR2+ cells were detected in regions with morphological features of ADM, characterized by epithelial structures in acinar-rich areas expressing markers such as PanCK and KRT19 (**Fig. 9C**). PanIN lesions, which are downstream of ADM, showed stronger expression of AGR2, GATA6, and also expressed TFF1, an additional gastrointestinal marker (**Fig. 9D**). Thus, the Classical-1 program is more firmly established in PanIN lesions, consistent with Carpenter *et al*. research findings. Unexpectedly, robust AGR2 expression was also observed in interlobular ducts and in larger morphologically intact ducts that are unrelated to ADM or PanIN (**Fig. 9B, F**; control AGR2-negative ducts shown in **Fig. 9E**, left panel). Within ducts, AGR2 expression was frequently mosaic, indicating that AGR2 activation occurs focally among individual cells in a duct without disrupting morphology (**Fig. 9E**, right). AGR2+ cells showed marked upregulation (4.8-fold) of MMP7 (**Fig. 9C** and **Fig. S18E**) compared to AGR2-negative ductal cells, consistent with the notion that activation of the Classical-1 program is related to stress or cell injury. These findings show that activation of the Classical-1 program is not restricted to metaplasia or precursor lesions but is also unexpectedly present in morphologically intact ducts.

The above observations were made from tumour-adjacent tissue where field cancerization effects cannot be ruled out. To address this, we analyzed an independent dataset from pancreas from cancer-free organ donors and patients. This dataset was generated using Deep Visual Proteomics (DVP), an integrated spatial proteomic approach that combines pathology-guided region selection with high-sensitivity mass spectrometry (Min et al., 2025). The study profiled 115 histologically annotated regions (normal ducts, precursor lesions, and invasive carcinoma) from 20 individuals. Small regions containing approximately 100 phenotypically matched cells were isolated from formalin-fixed paraffin-embedded tissue and subjected to quantitative proteomic profiling (**Fig. 9F**). AGR2 expression was detectable by proteomics in normal ducts from cancer-free donors (**Fig. 9G**) and was modestly higher in normal ducts from patients (**Fig. S18E**). These data recapitulate the IMC and Carpenter *et al*. observations. This shows that AGR2 expression can be detected in normal pancreatic ducts from cancer-free donors and patients. Proteomic profiling allowed us to study additional Classical-1 markers. Several proteins, including LGALS4 and FAM3D, correlated with AGR2 expression in normal ducts (**Fig. 9F**), which supports broader activation of the Classical-1 program. However, other Classical-1 markers, such as CLDN18, GATA6, and VSIG2, were not found to correlate with AGR2 expression. IMC data showed that AGR2+ ducts express little to no TFF1 (**Fig. S18B**), and given that precursor lesions (PanIN) show strong activation of this gastrointestinal marker, suggests that the Classical-1 program is only partly activated in this setting. This partial activation likely explains why AGR2+ ductal cells retain normal duct morphology and do not resemble precursor lesions.

A key advantage of the DVP approach is its ability to detect *KRAS* hotspot mutations as tryptic peptides. While KRAS mutant peptides (ex. G12D/G12V/G13D) were readily detected in high-grade PanIN, KRAS mutant peptides were not detected in AGR2 expressing normal ducts from cancer-free donors or patients (**Fig. 9H**). This mirrors our single-cell mutation analysis showing that the Classical-1 program is established prior to *KRAS* mutations. Collectively, these data establish the Classical-1 program as an early epithelial reprogramming event that arises in multiple contexts including metaplasia but also in morphologically intact ducts. This work highlights mouse studies from Ferreira *et al*. (2017) that showed that Agr2+ ducts give rise to Classical tumours as an alternate cellular route to the development of Classical tumours.

## Discussion

This study advances our understanding of epithelial cell plasticity in human pancreatic cancer. For a decade, the Basal-Classical framework has served as a critical guide in how we view transcriptional heterogeneity in this disease and has also been an important prognostic tool for patients. Our integrated analysis shows that the biology of the transcriptional states of this cancer is more complex. While prior single-cell studies have suggested this, their results have been difficult to apply directly to patient tumours. The scale of our cohort combined with tumour enrichment has allowed us to comprehensively study the transcriptional architecture of these tumours. Through a series of analyses, our work shows that despite originating from the pancreas, the transcriptional states of these tumours are not cell-of-origin identities. Perhaps this suggests that the original programs that regulate acinar and ductal cells are recalcitrant to transformation and require reprogramming. Importantly, we show that the transcriptional states of this disease recur in other tissues and malignancies.

Much of our focus was on the Classical-1 program as this is the main transcriptional state of pancreatic cancer. This should not be confused for a simple gastric program but reflects a conserved pan-gastrointestinal epithelial state. The enrichment of this program in chronic pancreatitis, and the fact that it aligns to reactive biliary cell populations, as well as its detection in cerulein mouse models indicate that this program is a reprogramming event due to cell injury. Why this particular program is activated is unknown, but given it is associated with mucin production, this suggests that it represents a protective response and may reflect a memory of the foregut origins of this tissue. An important advance from our previous work is the resolution of the Classical-2 program. Much remains to be studied about this rare program; however, this signature represents a rare neuroendocrine-like epithelial state that has been found in mouse models. Regarding the Basal programs, our data show that this is not a single continuous program. Basal-1 is a squamous state, whereas Basal-2 a plasticity-associated state that mediates transitions between epithelial programs. This distinction is important as adeno-to-squamous transition in lung drives resistance to KRAS inhibition (Tong et al., 2024), whereas, in pancreatic cancer, reactivation of partial EMT is linked to resistance to KRAS inhibition (Dilly et al., 2024). Thus, distinguishing these states could be important as these therapies emerge in the clinic.

An important contribution of this work is the timing of the emergence of the Classical-1 program. Using aneuploidy as a marker of transformation, we show that the Classical-1 program is present in non-neoplastic epithelial cells that we labelled Duct2. This is consistent with recent studies demonstrating that pancreatic ductal epithelium is not homogeneous, but includes a distinct ONECUT2-positive population associated with metaplasia and support the existence of pre-malignant epithelial intermediates within the ductal compartment (Arcila-Barrera et al., 2026; Schlesinger et al., 2020). Single-cell mutational analysis of KRAS showed that these cells lack activating mutations in KRAS. Together with the spatial proteomic data, this supports that this important transcriptional state in pancreatic cancer is perhaps a key plasticity state change that occurs before KRAS mutations. This is in line with work by David and colleagues showing that KRAS mutations help to ‘lock in’ a particular disease state rather than initiating a disease state (Li et al., 2021). Our data show that Classical-1 tumours are strongly associated with KRAS mutation, particularly the G12D allele. We interpret this cautiously, but this may support a model where epithelial reprogramming provides the right cellular context for KRAS mutations to persist.

We provide spatial and protein-level validation of the Classical-1 program using multiplexed IMC and spatial proteomics. AGR2, a core marker of the Classical-1 program, serves as a useful marker to track this program in tissue. Our data support that ADM or morphologically normal ducts only partly activate the Classical-1 program. This is important as this may explain why mucin has not accumulated sufficiently in these cells to cause morphologic disruption. At the transcriptional level, this is consistent with Duct2 cells showing restricted activation of secretory programs, including SPDEF, AGR2, and MUC5B, but lacking the broader mucin repertoire observed in malignant states, indicating a primed rather than fully differentiated epithelial state. This is important as this may explain why mucin has not accumulated sufficiently in these cells to cause morphologic disruption. Importantly, these findings are consistent in both tumour-adjacent patient tissue and cancer-free donors, indicating that our results do not derive from a cancerization effect of the tumour. Overall, our study represents a fundamental advance in the biological understanding of the transcriptional states of pancreatic cancer. These states are not simple subtype labels but reflect conserved epithelial plasticity programs that arise outside of the pancreas lineage and prior to oncogenic transformation. By integrating genomic, transcriptomic, and spatial data, we show that tumour cell state is established before mutation and is largely independent of subsequent genomic evolution. These findings support a model in which epithelial plasticity defines a permissive cellular context that precedes and shapes tumour initiation. This framework provides a new way to understand tumour heterogeneity and suggests that early cell state changes, rather than genetic alterations alone, may represent key targets for early detection and therapeutic intervention.

## Methods

### Patient tumour specimens

All patients provided written informed consent allowing the molecular characterization of their tumor samples and follow-up on their clinical information under the International Cancer Genome Consortium (ICGC) protocol. Patient samples were mostly accrued at Princess Margaret Cancer Centre at the University Health Network (Toronto, Canada). Resectable tumors were obtained from the UHN Biospecimens Program, and advanced tumors were obtained from the COMPASS trial (no. NCT02750657). As part of previous studies (Aung et al., 2018; Connor et al., 2019, 2017; Notta et al., 2016), some resectable tumors were also obtained from Sunnybrook Health Sciences Centre (Toronto), Kingston General Hospital (Kingston), McGill University (Montreal), Mayo Clinic (Rochester), Massachusetts General Hospital (Boston) and Sheba Medical Centre (Tel Aviv). Approval for the study was obtained through the University Health Network Research Ethics Board (nos. 15-9596, 13-6377, 18-5116, 20-5594, 21-5648 and 32517). Patient cohort data are summarized in Supplementary Tables 1 and 2. This study complies with all relevant ethical regulations. Tumour samples were obtained and processed as described previously (Connor et al., 2017; Notta et al., 2016). Briefly, resectable tumours were obtained from surgical specimens and advanced tumour cores were obtained by image-guided percutaneous core needle biopsy.

### Cell enrichment using laser capture microdissection for bulk sequencing

Fresh tumors were embedded in optimal-cutting-temperature (OCT) compound and snap-frozen in liquid nitrogen. Frozen biospecimens for bulk RNA-sequencing and whole genome sequencing (n = 490 from 464 patients) underwent laser capture microdissection (LCM) to enrich for tumour cells as described previously (Connor et al., 2017; Notta et al., 2016).

### Bulk RNA sequencing

RNA was extracted from LCM tissue using the PicoPure RNA Isolation Kit (ThermoFisher Scientific) and quantified using the Qubit dsRNA High Sensitivity Kit (Invitrogen). Quality was assessed using the RNA ScreenTape Assay on the 2200 TapeStation Nucleic Acid System (Agilent Technologies).

RNA with RIN (RNA integrity number) > 7 was used to prepare sequencing libraries using the TruSeq RNA Access Library Sample prep kit (Illumina) or the TruSeq Stranded Total RNA Library Prep Gold kit (Illumina) according to the manufacturer’s instructions. Libraries were then quantified using the KAPA Illumina Library Quantification Kit (Roche) according to the manufacturer’s protocol. Paired-end sequencing was carried out on the Illumina HiSeq 2500 platform (2x126 cycles) or the NovaSeq 6000 (2x151 or 2x101 cycles).

RNA sequencing (RNA-Seq) was conducted at the Ontario Institute of Cancer Research following the protocol outlined by O’Kane et al. (O’Kane et al., 2020). Sequencing reads were aligned to the human reference genome (hg38) and transcriptome (Ensembl v100) using STAR v.2.7.4a.

Duplicate reads were marked with Picard v.2.21.4. Gene expression levels were quantified in transcripts per million (TPM) using the stringtie package v.2.0.6, and log2-transformed expression values were used for subsequent analyses.

### Whole-genome sequencing (WGS) and analysis

DNA was extracted from LCM tissue using the Gentra Puregene Tissue kit components (Qiagen). The Gentra Puregene Blood Kit (Qiagen) was used to extract DNA from buffy coat. DNA was then quantified using the Qubit dsDNA High Sensitivity Kit (Invitrogen). Sequencing libraries were prepared using either the NEBNext DNA Sample Prep Master Mix Set (New England Biolabs), the Nextera DNA Sample Prep Kit (Illumina) or the KAPA Library Preparation Kits (Roche), following the manufacturer’s instructions. Libraries were quantified on the Illumina Eco Real-Time PCR Instrument using KAPA Illumina Library Quantification Kits (Roche) according to the manufacturers protocol. Paired-end sequencing was carried out on the Illumina HiSeq 2000/2500 platform (2x101 or 2x126 cycles) or the NovaSeq 6000 (2x151 or 2x101 cycles) targeting a collapsed coverage of 50X and 35X for tumour and normal samples, respectively.

Sequencing reads were aligned to the human reference genome (hg38) using Burrows-Wheeler Aligner (BWA, v0.7.17). Germline variant calling was performed using the Genome Analysis Toolkit (GATK, v4.1.2). Somatic single nucleotide variants (SNVs) were identified by the intersection of calls from Strelka2 v2.9.10 and MuTect2 v4.1.2. Indels were called based on consensus from at least two of four tools: Strelka2, MuTect2, SVABA v134, and DELLY2 v0.8.1. Structural variants (SVs) were similarly defined by consensus from at least two of the following: SVABA, DELLY2, and Manta v1.6.0. Copy number alterations, tumor cellularity, and ploidy were estimated using HMMcopy v0.1.1 and CELLULOID, an internally developed algorithm described in Notta et al. (2016). CELLULOID infers allele-specific copy number states and estimates tumor purity and ploidy by fitting copy number profiles to cellular mixtures. KRAS imbalance status was defined using the mutant and wildtype copy numbers of KRAS as in Knox et al (2025). Balanced KRAS status was defined as equal mutant to wildtype copies. Tumours with 3 or more copies of mutant KRAS than wildtype were called major KRAS imbalance tumours, and fewer than 3 as minor.

The numbers of subclone clusters across the WGS cohort were determined as described in Fang et al. (BioRxiv, 2025), excluding samples with cellularity < 30%. Briefly, mutant copies of SNVs were calculated correcting for cellularity and ploidy. To identify the subclonal clusters, the distribution of mutant copies was clustered using mclust v6.1 and the best model was selected using a combination of Bayesian Information Criterion, Integrated Complete Likelihood and median uncertainty for each sample. For the initial identification of clusters, SNVs with a depth of less than 30 were treated as noise, but once clusters were identified, they were included in the remaining analysis. Subclonal clusters were defined as clusters with the median mutant copy count below 0.75.

### Non-negative matrix factorization (NMF)

Non-negative matrix factorization (NMF) was leveraged to extract transcriptional program signatures. Transcripts per million (TPM) data were utilized for this analysis. Initial filtering retained genes expressed in more than 25% of the samples. Further filtering included only protein-coding genes with a minimum length of 500 nucleotides. Small nucleolar RNA (snoRNA) and ribosomal RNA (rRNA) were excluded from the dataset. Duplicate genes were identified and removed to ensure data integrity. K-means clustering was performed, and further genes were discarded to improve stability. The remaining data were renormalized to TPM. NMF was executed 200 times for each rank from 2 to 20 on both the cohort and randomized data using a high-performance computing cluster using R 4.2.1 and NMF v.0.26. The ranks were evaluated based on their ability to extract relevant biological signals as well as tumour signatures described by Chan-Seng-Yue et al. (2020). Additionally, we compared ranks with those previously identified from Chan-Seng-Yue et al. Standardized weights of genes from each previous programs (BasalA, BasalB, ClassicA, ClassicB, fibroblast, liver, immune and pancreas) were compared against the other genes in the same component from NMF W matrix using the Wilcoxon rank sum test. The lowest rank at which all previous programs were identified distinctly, rank 15, was selected for downstream analysis. For selection of top NMF genes, a standardized W matrix was generated by first normalizing each column of the weight (W) matrix by its column sum, followed by scaling each row by the median gene expression. This approach makes weights comparable across both components (columns) and genes (rows). Genes with the highest weights in each component were then selected as the top NMF genes and used for downstream analyses. Specific interrogation of genes enriched in the signatures revealed that the Classical-2 component (enriched in ClassicB) featured several acinar genes as top genes and the Classical-2 component from rank 16 was extracted instead.

### Hierarchical and consensus clustering

Hierarchical clustering was applied to the weight matrices from each run, and the results were analyzed to identify consistent clusters. An average weight matrix was calculated and consensus clustering using ConsensusClusterPlus v.1.60.0 was performed to ensure the stability of the identified clusters.

### Single-sample classifier

Starting with cluster annotations derived from hierarchical clustering, we performed gene filtering informed by the sparsity inherent to our scRNA-seq cohort to enhance classifier generalizability. We then applied a multi-level bootstrap-aggregated K-top scoring pairs (K-TSP) framework to robustly identify gene pair comparisons capable of distinguishing between subtyping classifications. Expression matrices underwent normalization—either through relative median scaling, log-transformation adjusted by median values, or remained untransformed—as appropriate to the data context. For each binary comparison between annotated classes, we employed a hierarchical bootstrapping approach consisting of nine iterative layers, each comprising up to 1,000 independent bootstrap replicates. During each iteration, subsets of samples (50%) and gene features (2,000 genes) were randomly or probabilistically selected based on previous iteration outcomes to train K-TSP classifiers using the R library switchBox v1.38.0, with k-values ranging from 3 to 50. Classifiers generated at each bootstrap step were immediately validated on the full dataset, and the frequency of gene pairs selected across replicates was aggregated to determine their relative predictive importance. These aggregated gene pair frequencies guided subsequent iterations and were ultimately used to construct a binary tree classifier to systematically assign subtype classifications.

### Differentially expressed genes and gene set enrichment

Transcripts per million (TPM) data were processed using Limma v.3.60.2 to filter (filterbyexp with min.count=1 and minimum number of samples based on the model program definitions), normalize and calculate differential expression. Differential expression was calculated comparing each tumour program to the remaining tumour program (excluding Hybrid tumours).

Each program-specific rank file was analyzed using Gene Set Enrichment Analysis (GSEA, v4.3.2) (Subramanian et al., 2005), PreRanked with parameters set to 2000 gene-set permutations and gene-sets size between 15 and 500. Genes were ranked according to the negative log of their p-value times the sign of the differential expression indicating the magnitude and direction of differential expression. The gene sets included for the GSEA analyses were obtained from MsigDB-c2, NCI, Biocarta, IOB, Netpath, HumanCyc, Reactome, Panther, WikiPathways and Gene Ontology (GO) databases excluding inferred from electronic annotation terms, updated August 1, 2024 (http://download.baderlab.org/EM_Genesets/). An enrichment map (version 3.4.0 of Enrichment Map software; Merico et al., 2010; Reimand et al., 2019) was generated using Cytoscape v3.10.2 using only enriched gene-sets with threshold FDR <0.01. Similarity between gene sets was filtered by combined coefficient (constant = 0.5) greater than 0.375.

### Generating signature-scoring differential expressed gene sets

Program scores were generated using differentially expressed genes from each bulk RNA-seq program signature with annotations derived from the single-sample classifier. Differentially expressed genes were filtered for adjusted p-values less than 0.01 and log fold change greater than 1. Pi value (Xiao et al., 2014) was calculated using pi_value = logFC * -1 * log10(adjusted p-value) and the top 100 genes by pi-value were utilized for signature scoring after removing genes with disproportionate expression in the liver and pancreas (e.g. *PRSS2*, *REG1A*) that were deemed to be contamination and were filtered out.

### Fresh tumour processing for single cell sequencing and organoid initiation

Tumours from freshly resected tumors and fresh core biopsies were chopped finely using a razor blade in 1 ml cold dissociation medium (IMDM/2% fetal bovine serum (FBS) with 2X collagenase/hyaluronidase mix (Stemcell Technologies) and 10 mg ml–1 DNaseI (Millipore)) on ice-cold 10 cm tissue culture Petri dishes. Final dissociation volume was brought up to 10 ml and the sample was placed on a rotator at 4 °C for overnight incubation. The following day, the sample was passed through a 100-μm nylon mesh, washed with 30 ml of IMDM/2% FBS (Gibco), centrifuged and cell pellet was resuspended IMDM/2% FBS for viably freezing or downstream applications.

### Single cell RNA sequencing processing

Post tumour processing, tumour cells were enriched through depletion of CD45+/CD90+/GlyA+ populations using MS columns (Miltenyi Biotec) following the manufacturer’s instructions. After enrichment, cells were resuspended in PBS/0.5% BSA and used for single-cell RNA library preparation using the Chromium Single Cell 3’ v2 kit or Chromium Next GEM Single Cell 3’ v3.1 Kit (10X Genomics) following the manufacturer’s instructions. Libraries were quantified using the Qubit dsDNA High Sensitivity Assay (Invitrogen) and quality was assessed using the LabChip GXII Touch DNA High Sensitivity kit (Perkin Elmer). Cluster generation and sequencing was performed on the Illumina HiSeq 2500 platform using HiSeq PE Cluster Kit v4 and HiSeq SBS Kit v4 (Illumina) for High Output mode, or HiSeq PE Rapid Cluster Kit v2 and HiSeq Rapid SBS Kit v2 (Illumina) for Rapid Run mode or NovaSeq 6000 targeting 40,000 reads per cell.

### Single cell RNA-seq data analysis

Single cell RNA-seq data were aligned to hg38 using CellRanger (v3.0.1, v3.1.0; 10X Genomics). Quality filtering, normalization and dimensionality reduction were performed in python v3.7.9 with scanpy v1.8.1. Cells with high mitochondrial content (>15%) and fewer than 400 genes expressed were removed. Doublets were predicted using scrublet v0.2.3. Samples were merged and counts normalized to counts per 10000 (scanpy.pp.normalize_total). Using the SAM algorithm v0.8.1 (Tarashansky et al., 2019), the merged dataset was clustered with leiden clustering (res 0.8, 50 neighbours), and underwent principal component analysis and UMAP dimensionality reduction. Clustering, principal component analysis and UMAP dimensionality reduction were repeated on individual samples and celltype-specfic subsets of the merged dataset. Processed data were saved as anndata h5ad files and converted to Seurat v5.2.1 (Hao et al., 2024) RDS files using hdf5r v1.3.12 in R v4.3.0 for other subsequent analysis.

### Cell-typing and gene set scoring

Cells were scored for marker genes as described in Tirosh et al. (2016) and Satija et al. (2015) (scanpy.tl.score_genes in python, Seurat’s AddModuleScore in R). Briefly, the average expression of the genes of interest are subtracted by the average expression of an expression-bin-sampled reference set of genes. Cell types were assigned to each cluster at the merged dataset level using the highest scoring cell type marker genes. Gene sets and marker gene lists can be found in Table S5. Malignant cell types were further confirmed using haplotype-aware inferred copy number tool Numbat v1.4.2 (Gao et al., 2023). Numbat was run on individual single cell RNA-seq samples using input counts and BAMs from Cellranger (run as described previously) with default parameters, using the default HCA data as reference. Additionally, Duct1 and Duct2 gene sets (Cui Zhou et al., 2022) were scored in duct cells and each duct cell was assigned to the group by highest score. Cell type information was used to confirm findings from the bulk NMF and DE analysis by scoring for the NMF and DE genesets using AddModuleScore as described above.

### Single cell classification and continuum analysis

SingleR v2.4.1 (Aran et al., 2019) was used to project tumour classification labels onto malignant single cells. Using the 490 bulk tumour cohort with Hybrid tumours removed as reference, SingleR was run using default parameters on the single cell cohort. Cell classification and scores were determined as output by SingleR. Classical-1 and Basal-2 cells were further assigned a Mixed classification if the absolute difference between the Classical-1 and Basal-2 scores was less than 0.02. Principal component analysis was run on the scores output by SingleR and PC1 was used as the cell program trajectory.

Malignant cells were divided into 100 nearly equal bins for further analysis. To validate the trajectory, Slingshot v2.10.0 (Street et al., 2018) was used to run pseudotime analysis on malignant cells of individual scRNA-seq samples in principal component space, clustered with kmeans (n=10). The generated trajectory was rooted by the cluster with the highest proportion of cells found to be earliest phylogenetically by Numbat analysis. Individual tumours with at least 50 malignant cells and excluding the neuroendocrine tumour (n=44/48) were grouped by malignant cell classification proportion by plotting Classical cell proportion against Basal cell proportion and clustering with k-means (n=4).

### Comparison of programs with Gavish et al

Pan-cancer malignant transcription programs from Gavish et al. (2023) were scored in our neoplastic single cell dataset using AddModuleScore from Seurat v5.1.0.

Cancer type enrichment analysis for each pancreatic cancer state was assessed based on the proportion of tumours from a given cancer type that harbor malignant programs aligned with that state, as shown in the heatmap. Statistical significance was evaluated using a one-sided Fisher’s exact test, with Bonferroni correction applied to control for multiple testing (adjusted alpha=1.35e-3).

All cancer type enrichment results were further validated at the gene level. For each enriched cancer type within a given pancreatic cancer state, we selected top-ranking genes (typically 3–20 per program, depending on program size and gene overlap) and projected them into our single-cell dataset. Pseudobulk differential expression analysis was then performed for each individual gene using FindMarkers in combination with MAST v1.28.0. Gene-level p-values (Wilcoxon rank sum test, output by FindMarkers) were subsequently combined using the Cauchy combination method (ACAT v0.91; Liu and Xie, 2020) in R.

### KRAS exon 2 hotspot amplification

KRAS exon 2 hotspot amplification was adapted from Nam et al. (2019). Universal forward PCR primer was designed with the Illumina P5 and partial Read 1 sequencing primer sequence. Reverse PCR primers were designed with 20bp spanning *KRAS* Exon 1 and Exon 2 boundary, Illumina Read 2 sequencing primer sequence, 8bp unique sample index, and P7 sequence. PCR was performed using 0.5ng of cDNA (produced from the Chromium Single Cell 3’ v2 or Chromium Next GEM Single Cell v3.1 protocol), 3% DMSO and 1X Phusion Master Mix (New England Biolabs) for 30 PCR cycles with annealing temperature of 65°C. PCR products were subjected to 0.7X SPRIselect (Beckman Coulter) bead cleanup and were quantified using Qubit dsDNA High Sensitivity Assay (Invitrogen). Samples were then pooled and sequenced on MiSeq using Micro v2 kit (Illumina) to generate reads spanning Cell Barcode and UMI (Read 1) and *KRAS* G12 hotspot (Read 2).

KRAS reads were aligned to hg38 using CellRanger v3.1.0. Variant numbers were counted using Vartrix v1.1.3 (10X Genomics) using commonly mutated KRAS variants in VCF as reference. Sample-specific KRAS variants were identified and counted per UMI. Bam files were viewed using Integrative Genomics Viewer (IGV) v2.4.16.

### Penalized Module Score

Standard methods for scRNA-seq gene set (module) scoring may yield inflated scores driven disproportionately by a few highly expressed genes. For modules where broad expression of genes was important, we developed PenalizedModuleScore, an extension to Seurat’s AddModuleScore.

PenalizedModuleScore applies a penalty factor to inflated scores arising from uncoordinated gene expression within modules, allowing a module score that better reflects expression of all genes within a desired module.

The penalization procedure is implemented as follows: for each cell and each gene set (module), we calculate the fraction of genes whose expression exceeds the mean expression of matched control genes. Specifically, for cell 𝑐, given module genes 𝑀 and matched control genes 𝐶, we determine the fraction 𝑓_!_as:

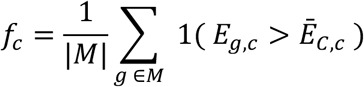

where *E_g,c_* is the expression level of gene *g* in cell *c*, ̄𝐸*_c,c_* is the mean expression of the control gene set *C* in cell *c*, and 1( · ) is an indicator function.

This fraction *f_c_* is transformed into a penalty factor 𝑝*_c_* using a sigmoid function:

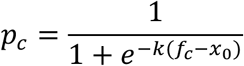

Here, 𝑘 controls the steepness of the sigmoid curve, determining how sharply the penalty transitions (default *k* = 25), and *x*_0_ represents the midpoint, the fraction at which the penalty equals 0.5 (default *x*_0_ = 0.25). Consequently, cells with coordinated and broad expression across module genes experience minimal penalty, whereas those dominated by a small subset of highly expressed genes are significantly penalized.

The final penalized module score per cell is computed by scaling the original module score (the difference between module and control scores obtained via AddModuleScore) by the penalty factor 𝑝*_c_*:

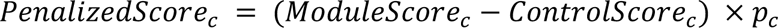

### Reference scRNA-seq datasets

To validate and contextualize our program signatures, we incorporated publicly available single-cell RNA sequencing datasets spanning various developmental and disease states. These included datasets from mouse models of pancreatic injury and oncogenesis; human datasets encompassing fetal, neonatal, and adult pancreatic tissue; as well as human datasets spanning multiple other organs.

Specifically, we utilized:

- A murine pancreatitis dataset from Ma et al. (2022) profiling acinar-to-ductal metaplasia (ADM) following injury; Normalized and scaled before performing PCA and UMAP reductions.
- A murine oncogenic KRAS^G12D^ dataset from Schlesinger et al. (2020), capturing preinvasive lesions and acinar cell heterogeneity; Subsetted to the epithelial compartment, normalized and scaled before performing PCA and UMAP reductions.
- The Human Transcriptome Cell Atlas (HTCA) dataset from Pan et al. (2023), comprising single-cell transcriptomes across human organs; Immune cell types along with cells from adrenal gland, blood, kidney, muscle, ovary, spleen and thymus were not used. Data was subsetted and renormalized before performing PCA and UMAP reductions.
- A human fetal pancreas dataset from Migliorini et al. (2024)
- And a single-nucleus and in situ RNA-sequencing dataset from Tosti et al. (2021), profiling both neonatal and adult human pancreatic tissues, including samples from the chronic pancreatitis setting.

Datasets were downloaded from the respective public repositories. Any additional processing was conducted using Seurat v5.1.0. Scores were generated using penalized module scoring on gene sets. Differentially expressed genes were also obtained from Ma et al. and the epithelial subset in Schlesinger et al. Genes were filtered for up to the top 100 genes by Pi value (as previously described) and then converted to human genes by one-to-one orthologs. The 26 sets of differentially expressed genes, representing the 26 clusters from Schlesinger et al. were additionally filtered by Gini score to remove universally expressed genes. Genes with gini > 0.7 (ineq v0.2-13) within the tumour scRNA-seq dataset were kept for subsequent scoring with PenalizedModuleScore.

### Other transcriptome datasets

Normalized GeoMx data from Carpenter et al. (2023) were downloaded and separated by donor. Genes of interest were plotted after donor-level z-score normalization. Additional single cell RNA-seq data collected across 229 tumours from multiple studies from Loveless et al. (2025) were scored using AddModuleScore for all programs.

### Patient-derived organoid initiation and culture

Cells from dissociated tumour were resuspended in 100% Matrigel (Corning) and embedded onto 24-well plates. Matrigel domes were solidified after a 30-minute incubation at 37 °C. After polymerization, matrigel domes were covered by 0.5 ml conditioned media, and cells were cultured at an environment of 37°C with 5% CO2, which allows the formation of patient-derived organoids (PDOs). During the PDO cultivation, conditioned media was replaced into fresh one every 3 days, and cells were passaged every 10-14 days depending on their growth rate. Conditioned media consists of 50% Advanced DMEM/F-12 media (Gibco), 20% Wnt-3a conditioned media and 30% Human R-Spondin1 conditioned media (Princess Margaret Living Biobank), further supplied with 1x B-27 supplement (Gibco), 2 mM GlutaMax (Gibco), 10 mM HEPES (Gibco), 10 mM nicotinamide (Millipore Sigma), 1.25 mM N-Acetyl-L-cysteine (Millipore Sigma), 10 nM Gastrin I(1-14) (Millipore Sigma), 0.5 µM A 83-01 (Tocris Bioscience), 10 µM Y-27632 (Selleckchem), 1:500 Primocin (InvivoGen), 100 ng/ml recombinant human noggin (Peprotech), 100 ng/ml recombinant human FGF-10 (Peprotech) and 50 ng/ml recombinant human EGF (Peprotech). All the experiments in this study were completed before cells’ passage number reached to 20.

### Patient-derived organoids - bulk RNA-seq

Patient-derived organoids were initiated and cultured as described above. At passage 6 or after, organoids are dissociated using TrypLE (Life Technologies) into single cells, pelleted and frozen at - 80°C. RNA was extracted using the PicoPure RNA Isolation Kit (ThermoFisher Scientific) and quantified using Qubit RNA Assay (Invitrogen). 400ng RNA was subjected to library prep using Illumina Stranded Total RNA Prep Ligation with Ribo-Zero Plus. Sequencing was performed on the NovaSeq 6000 with read length PE150 targeting 50 million reads per sample. Sequencing data were processed identical to bulk tumour RNA-seq, as described previously. Variance stabilizing transformation (VST) was applied to the raw count matrix of 162 PDOs using DESeq2 (v1.42.0). The resulting VST-transformed matrices for the Moffitt gene set and the differential expression (DE) gene set are shown as two separate heatmaps generated by ComplexHeatmap v2.18.0. PDO samples are displayed in the same order across heatmaps, with columns split based on k-means clustering derived from the DE gene set.

### Single cell multiome analysis

8 pancreatic cancer cases from resected tumors were dissociated and enriched for tumour cells using methods described previously. Cells were either processed fresh or were viably frozen and thawed for the assay. Nuclei Isolation and library preparation was performed using the Chromium Next GEM Single Cell Multiome ATAC + Gene Expression assay kit (10X Genomics) according to the manufacturer’s instructions. Sequencing was performed on NovaSeq 6000 targeting an average sequencing depth of 50,000 reads per nucleus for mRNA and 30,000 reads per nucleus for ATAC-seq.

Multiome data were aligned to hg38 using CellRanger ARC v2.0.0 (10X Genomics). Quality filtering, normalization and dimensionality reduction were performed in R with Seurat and Signac v1.14.0 (Stuart et al., 2021). RNA data were filtered to include only cells with low mitochondrial percentage (<20%), gene counts between 200 and 3 Mean Absolute Deviation (MAD) above the median, and total UMI counts within +/-3 MAD of the median. ATAC data were filtered to include only cells with nucleosome signal between the 2^nd^ and 98^th^ percentile, TSS enrichment above the 2^nd^ percentile, peak region fragments between the 2^nd^ and 98^th^ percentile, and a blacklist ratio of less than 5%. UMAP dimensionality reduction was performed jointly with RNA and ATAC data using a weighted nearest neighbour graph. Gene set scoring, cell classifcation and trajectory analysis for RNA data were performed as described previously for tumour scRNA-seq samples. ATAC data were analyzed further with ArchR v1.0.2 (Granja et al., 2021), inheriting trajectory data from the RNA data.

### Detection of cilia by Immunofluorescence

Immunofluorescent analysis of Alpha-Tubulin was conducted on a formalin-fixed, paraffin-embedded tissue microarray, comprising 20 cases of pancreatic cancer resections with 4 cores per case. Each case had paired whole transcriptome data. Antigen retrieval was carried out using a Tris-EDTA buffer at pH 9. For staining, the tissue was incubated overnight at room temperature with Alpha-Tubulin (Acetylated) Recombinant Mouse Monoclonal Antibody (Product # 32-2700) at a concentration of 1 µg/mL in 1% BSA. This was followed by a 1-hour incubation with Alexa Fluor 555 Rabbit Anti-Mouse IgG Secondary Antibody (Product # A21429), diluted 1:200. Nuclei were counterstained with DAPI (Product #62247) and were coverslipped using Fluorescence Mounting Media (Product#S3023). Slides were scanned at 40X magnification using a Zeiss AxioScan fluorescence scanner (AOMF, UHN) and evaluated by a trained pathologist.

### Morphological subtype analysis

Haematoxylin and eosin slides from patients with pancreatic ductal adenocarcinoma, alongside morphological subtype data were obtained from Flores-Figueroa et al. (BioRxiv, 2026). The relative proportion of morphological pattern, estimated as a percentage of total tumour area per tumour slide, was compared with tumour program scores of matching tumours (n=240) using Spearman correlation.

### Imaging mass cytometry

Imaging mass cytometry (IMC) analysis followed Shakfa and Nowlan et al. (BioRxiv, 2025). IMC data were generated from a large pancreatic ductal adenocarcinoma (PDA) cohort comprising 221 resected tumours, assembled into tissue microarrays (TMAs) with four spatially distinct cores per tumour (∼880 cores total). Formalin-fixed, paraffin-embedded sections were stained with a multiplexed panel of metal-conjugated antibodies targeting epithelial, stromal, and immune compartments, as described by Shakfa and Nowlan et al. (BioRxiv, 2025). Antibodies were optimized and validated prior to conjugation, and staining was performed uniformly across all samples. IMC acquisition was performed using Hyperion imaging systems at ∼1 μm resolution, enabling subcellular spatial resolution of protein expression. Laser ablation of tissue followed by time-of-flight mass spectrometry allowed quantitative detection of metal-tagged antibodies. Images were processed using a standardized pipeline, including spillover correction (CATALYST), normalization, and transformation of marker intensities. Single-cell segmentation was performed using Mesmer (DeepCell), integrating nuclear and membrane markers (DNA, pan-cytokeratin, E-cadherin, CD45, SMA) to generate high-confidence cellular masks. Across all images, more than 10 million single cells were segmented and quantified. For the purposes of this analysis, the data was subset to the 84 TMA cores with more exocrine cells than malignant cells, and then further gated for cells positive for E-cadherin, panCK, Insulin or trypsin. This subset was then clustered using PhenoGraph and annotated into epithelial populations based on canonical marker expression.

### Deep visual proteomics and spatially resolved mass spectrometry

Deep visual proteomics (DVP) analysis followed Min et al. (BioRxiv, 2025). DVP was used to link tissue morphology with spatially resolved protein expression in pancreatic cancer specimens. Whole-slide H&E images were processed using an artificial intelligence-based pipeline to identify morphologically distinct regions of interest (ROIs). Tissue images were divided into high-resolution tiles (224 × 224 pixels at 0.24 μm/pixel), generating over 1.6 million tiles across the cohort. Each tile was embedded using a vision transformer-based foundation model (H-optimus-0), which captures histological features in a high-dimensional representation. Unsupervised k-means clustering was then applied to group tiles into morphologically similar regions, enabling both global pattern discovery across samples and identification of intra-sample heterogeneity. All regions were reviewed by expert pathologists. Selected ROIs were subjected to laser capture microdissection (LCM) guided by the AI-defined morphology. Approximately 100 phenotypically matched cells were isolated per region using a gravity-assisted laser microdissection system. This approach enabled precise sampling of spatially defined epithelial structures, including normal ducts, precursor lesions, and tumour regions.

Microdissected samples were processed using an automated workflow for protein extraction and digestion, followed by ultra-sensitive liquid chromatography–mass spectrometry (LC-MS/MS). Peptides were analyzed using an Orbitrap Astral mass spectrometer coupled with an Evosep One liquid chromatography system under standardized acquisition conditions. Protein identification and quantification were performed using the DIA-NN software suite, enabling reproducible and high-sensitivity detection of proteins from low-input samples. To specifically assess oncogenic mutations at the protein level, a custom spectral library was generated to detect common KRAS variants (including G12, G13, Q61, and A146 substitutions). This enabled direct detection of mutant KRAS peptides in spatially defined regions. Protein abundance data were log2-normalized and analyzed in R. Differential protein expression across spatial regions was assessed, and results were integrated with single-cell RNA-seq data to link protein-level findings with transcriptional states. This multimodal framework enabled validation of epithelial programs, including the Classical-1 program, at the protein level and within defined tissue architectures. Together, this approach provided spatially resolved, quantitative proteomic profiling of epithelial states across normal ducts, precursor lesions, and tumour regions, enabling direct comparison of transcriptional programs with their corresponding protein expression in situ.

## Data and code availability

Processed data will be publicly available on GEO and raw data will be available through EGA (<u>EGAS00001002543</u>). Processed single cell RNA-seq data will also be accessible in the form of an interactive cell browser through the UCSC cell browser (https://cells.ucsc.edu/). Code including the bulk and single cell classifiers will be made openly available on Github.

## Acknowledgements

We would like to thank all members of the Notta Laboratory and PanCuRx program at the Ontario Institute for Cancer Research (OICR) for their advice, discussion and review of the manuscripts. We thank R. Chandwani and F. Real for critical review of this manuscript. We are grateful for the participation of patients and their families in this study, as well as the contributions of the Princess Margaret Cancer Biobank, COMPASS, and Pathology Research Program Laboratory (UHN) teams. We thank the OICR Drug Discovery team for their assistance with the KRAS inhibitor experiment, Dr. Elvin Wagenblast for sharing the CRISPR protocol, and The Advanced Optical Microscopy Facility (UHN).

This study was conducted with the support of OICR (PanCuRx Translational Research Initiative) through funding provided by the Government of Ontario, the Wallace McCain Centre for Pancreatic Cancer supported by the Princess Margaret Cancer Foundation, the Terry Fox Research Institute, the Canadian Cancer Society Research Institute, the Canadian Institutes of Health Research, VFoundation, Ontario Ministry of Economic Development and the Pancreatic Cancer Canada Foundation. The study was also supported by charitable donations from the Canadian Friends of the Hebrew University (A. U. Soyka). We acknowledge the contributions of members of OICR’s Diagnostic Development (Tissue Portal; oicr.on.ca/programs/diagnostic-development/), Genomics (genomics.oicr.on.ca) and PanCuRx programs for sample management, genomic and transcriptomic sequencing, data analysis.

F. Notta is supported by the Gattuso-Slaight Personalized Cancer Medicine Fund, funding from the OICR, the Canadian Institutes of Health Research (no. 388785) and the Cancer Research Society (no. 23383). S. Gallinger is the recipient of an Investigator Award and Senior or Clinician-Scientist Awards from OICR. S. Ge was supported by scholarships from the Canadian Institutes of Health Research (CGS-M), the Ontario Graduate Scholarship, and the Princess Margaret Hospital Foundation Graduate Fellowships In Cancer Research.

## Contributions

S.Ge. and F.Notta led data analysis and interpretation. J.M.W., A.D., A.E., I.L., S.R., S.H., D.B., A.B., S.H., E.S.T., R.C.G., G.M.O’K., D.T., J.J.K., S.Gallinger, and F.Notta contributed to pathology, sample acquisition and clinical annotation. K.N., E.F.F., and Y.Z. processed tumour and organoid samples. J.X., G.H.J., A.Z., and M.C-S-Y. processed bulk RNA-seq and WGS data. P.T. and G.H.J. analyzed tumour bulk RNA-seq data. R.I. and V.V. assisted with tumour bulk RNA-seq analysis. S.Ge and J.X. analyzed single cell tumour RNA-seq and organoid RNA-seq data. P.T., S. Ge, A.Migliorini, S.A.-B. and N.S. analyzed external RNA-seq data under the supervision of M.C.N., O.P., and F.Notta. P.K. and S. Ge analyzed single cell multiome data under the supervision of F.G. and F.Notta. Imaging analyses were performed by E.F.F., A.E., Y.F., F.Nowlan., and J.M. under the supervision of A.Mund, A.Maitra, H.W.J., and F.Notta. S.Ge, P.T. and F.Notta wrote the manuscript. J.X., F.Nowlan, J.M., K.N., E.F.F., M.C.N., O.P., and S.Gallinger reviewed and edited the manuscript.

## Competing Interest Statement

No conflicts of interest are reported.

---

**Figure S1:** Cell type identification in scRNA-seq, related to Figure 1. (a) UMAP plots of all filtered cells from scRNAseq of human tumours. Cells are coloured by sample (left), malignant (aneuploid) status (middle), and cell type (right). Malignant cells are additionally separated by Moffitt subtype in the cell type UMAP plot. (b) Dot plot showing expression of marker genes across different cell types in the cohort. Dot size represents percentage of cells expressing the marker gene while dot colour indicates average gene expression of the marker gene in each cell type population. (c) Exemplar inferred copy number plot showing inferred copy number status (by Numbat) grouped by chromosome and across cells of sample 96460. Inferred amplifications (AMP) are indicated in red, deletions (DEL) by blue, and copy neutral loss of heterogeneity (CNLoH) by green. Cells are grouped by non-aneuploid (normal) cells and by aneuploid (malignant) clones. Additional cell-level information about program score and classification are annotated on the left.

**Figure S2:** NMF rank determination, related to Figure 1. (a) Component number by negative log p-value (Wilcoxon rank sum test) across all components across ranks up to rank 16. Each component was compared against reference signatures from Chan-Seng-Yue et al., as indicated by dot colour. Tumour programs were extracted from rank 15 and 16, the lowest ranks at which the reference signatures were distinctly enriched. (b) Comparison of the Classical-2 enriched component across rank 15 and 16, showing top genes by normalized NMF component score. Highlighted genes are marker genes of acinar cells.

**Figure S3:** Comparison of signature gene sets derived from NMF vs differential expression (DE), related to Figure 1. UMAP plots where cells are scored for gene sets from NMF (left) or from differential expression (DE, middle). Dotplots of expression of DE genes across cell types and Moffitt-subtyped malignant cells (right).

**Figure S4:** Comparison of this study with previous classification from Chan-Seng-Yue et al., related to Figure 1. (a) Comparison of heatmaps from Chan-Seng-Yue et al. (top) and this study (bottom). Arrows highlight groups of tumours with low confidence from Chan-Seng-Yue et al. (b) Boxplots comparing consensus probability by consensus clustering based on Chan-Seng-Yue et al. and this study. (c) Distribution of tumour classifications by stage. Number of tumours by stage and classification are indicated.

**Figure S5:** Summary of tumour programs and pancreatic ductal marker expression, related to Figure 2. (a) Mean expression of tumour programs across pancreas cell types from Migliorini et al. (Fetal pancreas) and Tosti et al. (Neonatal pancreas, Adult pancreas and Chronic pancreatitis). (b) Expression of normal pancreatic ductal marker genes from the Human Protein Atlas across cells in the Tosti et al. chronic pancreatitis dataset. (c) EnrichmentMap featuring the gene sets significantly enriched (FDR < 0.01) within each program. Closely related gene sets are joined by an edge, and clusters of gene sets are circled and described by a summarizing title from one of the encircled gene sets.

**Figure S6:** Comparison of tumour programs by Human Transcriptome Cell Atlas, cilia, and cellularity, related to Figures 2 and 3. (a) Dotplot of expression of Classical-1 genes across selected Human Transcriptome Cell Atlas cell types. Cell types were extracted from Classical-1 enriched cell types as shown in Figure 2e and cell types enriched in other tumour programs. Dot size indicates percentage of cells expressing the genes while dot colour indicates average expression from yellow (lowest) to blue (highest). (b) Representative immunofluorescence staining of DAPI (blue) and acetylated tubulin (red, indicative of cilia) for control normal testis and PDA tumours of each of the different tumour programs as determined by matched bulk RNAseq (top). Quantification of highest acetylated tubulin % across tumours with at least 100 histological sections (n=16), grouped by tumour program as determined by matched bulk RNAseq (bottom). (c) Cellularity across tumours by classification. The programs show no significant difference in cellularity (Kruskal-Wallis, p=0.23).

**Figure S7:** Further comparison of Basal programs, related to Figure 3. (a) Scatterplots comparing the relationship between Basal-1, Basal-2, Classical-1 and Classical-2 scores for all malignant cells, excluding the neuroendocrine tumour. Labels in plot corners indicate percentage of cells within each quadrant. Grey boxes outline the quadrant representing positive expression of both program scores per plot. (b) Heatmap of Moffitt and program scores of all malignant cells ordered on the program continuum along PC1 as in Figure 3B (top). Bar plot of cell classification proportions across 100 bins along the program continuum (middle). Scaled expression of EMT, partial EMT, KRAS, Raghavan et al., and cell cycle programs plotted as line plots across 100 bins of the program continuum (bottom). (c) UMAP of malignant cells collected across 229 samples from Loveless et al. Cells are coloured by Basal-1 (left) and Basal-2 (right) program scores, where yellow is the lowest expression of the program and purple the highest (d) Proportion of malignant cells proliferating by cell cycle program score grouped by cell classification. Basal-2 has the highest proportion of proliferating cells by Fisher’s exact test, p = 2.68e-77. (e) Distribution of tumour classifications across KRAS imbalance status for primary (top) and metastatic (bottom) tumours. Tables below respective plots indicate exact sample tumours. Categories are significantly different by Fisher’s exact test. Notably, in the metastatic setting, major KRAS imbalance tumours are enriched in Basal-2. (f) Heatmaps of 164 patient-derived organoids ordered by k-means (n=4) clustering along the program gene sets. Moffitt subtype is annotated above. Variance-stabilized expression of Moffitt genes (top) and program genes (bottom) are shown where red indicated higher relative expression and blue indicates lower relative expression.

**Figure S8:** scMultiome analysis of tumour programs, related to Figure 3. (a) UMAP plots of malignant cells plotted using integrated scATAC-seq and scRNA-seq data from scMultiome analysis. Points are coloured by cell classification (top) and sample (bottom). (b) Heatmap of program scores calculated on RNA data, ordered as a program continuum along PC1 for all malignant scMultiome cells (top). Matched mean accessibility (bottom) across 10 bins along the program continuum (bottom). (c) Bar plot of counts of recurrent significant transcription factor motifs enriched by program (left). Selected transcription factor gene (solid line) and motif (dotted line) enrichment across the program continuum (right).

**Figure S9:** Classical-2 comparison with endocrine and neural-like programs, related to Figure 2 and 3. (a) Dot plot showing expression of markers of the endocrine pancreas in the epithelial cells of the scRNAseq cohort neoplastic and non-neoplastic cell populations. Non-neoplastic cells are separated by cell type while neoplastic cells are separated by tumour program. Dot size indicates percentage of cells expressing the genes while dot colour indicates average expression from grey (lowest) to blue (highest). (b) Correlation between programs from Hwang et al. and programs from this study. Positive correlation is indicated by magenta and negative by green. Magnitude of correlation is indicated by dot size. Neuroendocrine-related programs are highlighted with the red box.

**Figure S10:** Comparison of expression of top pan-cancer programs, related to Figure 4. Dot plots of top-ranking genes from selected differing components of lung squamous cell carcinoma (LSCC) and colorectal cancer (CRC) compared across all Basal-1, Basal-2, Classical-2 and Classical-2 cells. Dot size indicates percentage of cells expressing the genes while dot colour indicates average expression from yellow (lowest) to blue (highest). For assessing significance, we performed differential expression analysis using pseudobulk data constructed from the top 500 cells with the highest scores in each cell state. The overall p-value was calculated by combining gene-level differential expression p-values (Wilcoxon rank sum test) using the Cauchy combination test (see Methods).

**Figure S11:** Transcriptomic and genetic variation in primary and metastatic tumours, related to Figure 5. (a) Mosaic plot of primary and metastasis numbers between the ‘Classical dominant’ group and other groups. Bar widths and heights are proportional to group numbers, numbers as labelled. The ‘Classical dominant’ group is significantly different from the rest by Fisher’s Exact Test, p=0.020. (b) Boxplots of unique copy number variant counts (top) and clone numbers (bottom) per sample separated by diversity group. Neither are significantly different by the Kruskal-Wallis test (p=0.86, p=0.42). (c) Boxplot of unique copy number variant counts by primary or metastatic tumour (top). There is no significant difference in number of unique inferred copy number variants between primary and metastatic tumours. Bar plot of clone numbers by primary and metastatic tumours (bottom). Samples with 3 or more clones are grouped together. While there are more samples with 3+ clones in the primary setting, the difference is not significant (Fisher’s exact test, p=0.17) (d) Bar plot of the number of subclonal clusters by primary and metastatic tumours from the larger whole genome sequencing analysis (WGS) cohort. Primary tumours significantly differ from metastatic tumours (Fisher’s exact test, p=3.4e-5).

**Figure S12:** Inferred copy number and phylogenetic analysis from scRNA-seq, related to Figure 5. (a) Proportion bar plots of each sample as plotted in Figure 5d showing the cell classification proportions of early and late clones (b) Haplotype-aware inferred copy number scatter plot illustrating RNA log fold change (logFC) and allele frequency (pHF) for the normal, stromal cells of sample 100070 across the genome. (c) UMAP plot of malignant cells from sample 100070 coloured by clone number (top) and cell classification (bottom). (e) Haplotype-aware inferred copy number scatter plot illustrating differences in RNA log fold change (logFC) and allele frequency (pHF) between different clones of sample 100070 across the genome. Regions are coloured by predicted copy number status. Chromosome 17, featuring a copy number loss common to all clones, is highlighted with a zoomed-in panel (right). NEU = neutral, CNLoH = copy neutral loss of heterogeneity, DEL = deletion, AMP = amplification.

**Figure S13:** Mouse model marker expression in human tumours, related to Figure 6. (a) Program scores across populations in the pancreatitis *Kras*-WT dataset from Ma et al. Converted human-mouse orthologs were scored using the penalized module score, where white indicates low consistent expression of the program and dark red indicates high consistent expression of the program. (b) Program scores across populations in the *Kras*-G12D mutant dataset from Schlesinger et al. Converted human-mouse orthologs were scored using the penalized module score, where white indicates low consistent expression of the program and dark red indicates high consistent expression of the program.

**Figure S14:** Rare non-aneuploid basal cells, related to Figure 7. (a) UMAP plots of all filtered cells from sample 97727. Cells are coloured by cell type (left), aneuploid probability predicted from inferred copy number analysis (right). Non-aneuploid population of interest is circled in red dotted lines. (b) Inferred copy number plot showing inferred copy number status (by Numbat) grouped by chromosome and across cells of sample 97727. Inferred amplifications (AMP) are indicated in red, deletions (DEL) by blue, and copy neutral loss of heterogeneity (CNLoH) by green. Cells are grouped by non-aneuploid (normal) cells and by aneuploid (malignant) clones. Additional cell-level information about program score and cell type are annotated on the left. The non-aneuploid population of interest with high Basal-1 program score is highlighted with a red rectangle. (b) UMAP plots of expression of genes *TP63*, *KRT5*, *KRT6A*, *KRT14*, *KRT17* and *S100A2* (left). Blue indicates high expression within the sample while yellow indicates no expression. Dotplot of the same genes from the UMAP grouped by cell type (right), highlighting expression of these genes in the non-aneuploid basal cell population unique to this sample (“Basal_97727”).

**Figure S15:** Comparison of transcriptionally similar cell populations by marker genes, related to Figure 7. (c) Heatmap showing expression of differentially expressed genes between normal endocrine (non-neoplastic) pancreas cells and neoplastic Classical-2 cells (b) Heatmap showing expression of differentially expressed genes between the Duct1 and Duct2 populations. Expression of the differentially expression genes are shown in the Duct1 and Duct2 cells of this study (left) in comparison to the Classical-1- (Ductal) and Classical-1+ (MUC5B+ Ductal) populations from Tosti et al (right). Tumour program genes are highlighted.

**Figure S16:** Expression of Duct1 and Duct2 marker genes in donors and patients from Carpenter et al., related to Figure 7. Heatmaps showing expression of Duct1-Duct2 differentially expressed genes as well as acinar marker genes in healthy pancreas donors (a) and patients (b) from the Carpenter et al. GeoMX dataset. Rows in each plot are genes scaled by z-score. Columns are regions grouped by original annotation.

**Figure S17:** Classical-2 is associated with KRAS wildtype, related to Figure 8. (a) Histograms comparing distribution of Classical-1 (top) and Classical-2 (bottom) program scores from the injury *Kras*-WT (orange, Ma et al.) and *Kras*-G12D (purple, Schlesinger et al.) metaplastic single cell datasets. (b) Bar plot of the top Classical-1 and Classical-2 cells, grouped by the originating model. Classical-1 and Classical-2 top cells differ significantly by injury *Kras*-WT or *Kras*-G12D model (Fisher’s exact test, p=6.6e-68) (c) Bar plot of *KRAS* variants for tumours of each program classification. *KRAS* variant distribution is associated with tumour program (Chi-Squared test, p=9.0e-15) and *KRAS* wildtype is specifically associated with Classical-2 (Chi-Squared test, p=2.4e-18).

**Figure S18:** Identification and characterization of Classical-1+ cells in tumour-adjacent pancreas from patients and cancer free donors, related to Figure 9. (a) Cell type composition of tumour-adjacent regions based on IMC, showing predominance of acinar, endocrine, and ductal populations, with minimal tumour contamination. (b) Dot plot of population markers by unsupervised clusters. Clusters of IMC data are further grouped into major epithelial and stromal populations, annotated by the bar to the right of the plot. Colours follow (a) and (c). Two distinct non-cancer ductal populations are highlighted with dashed lines. (c) Representative IMC image highlighting identification of residual tumour cells in tumour adjacent region in IMC data. (d) Protein expression profiles across annotated cell populations. (e) Quantification of AGR2 expression in normal ducts from cancer-free donors and tumour-adjacent tissue using DVP. * p < 0.05; ** p < 0.01; *** p < 0.001; **** p < 0.0001.

## Notes

### Competing Interest Statement

The authors have declared no competing interest.

