## Supplemental Figures for "Extra-lineage tissue programs define the transcription states of human pancreatic cancer"

### Supplemental Figure 1

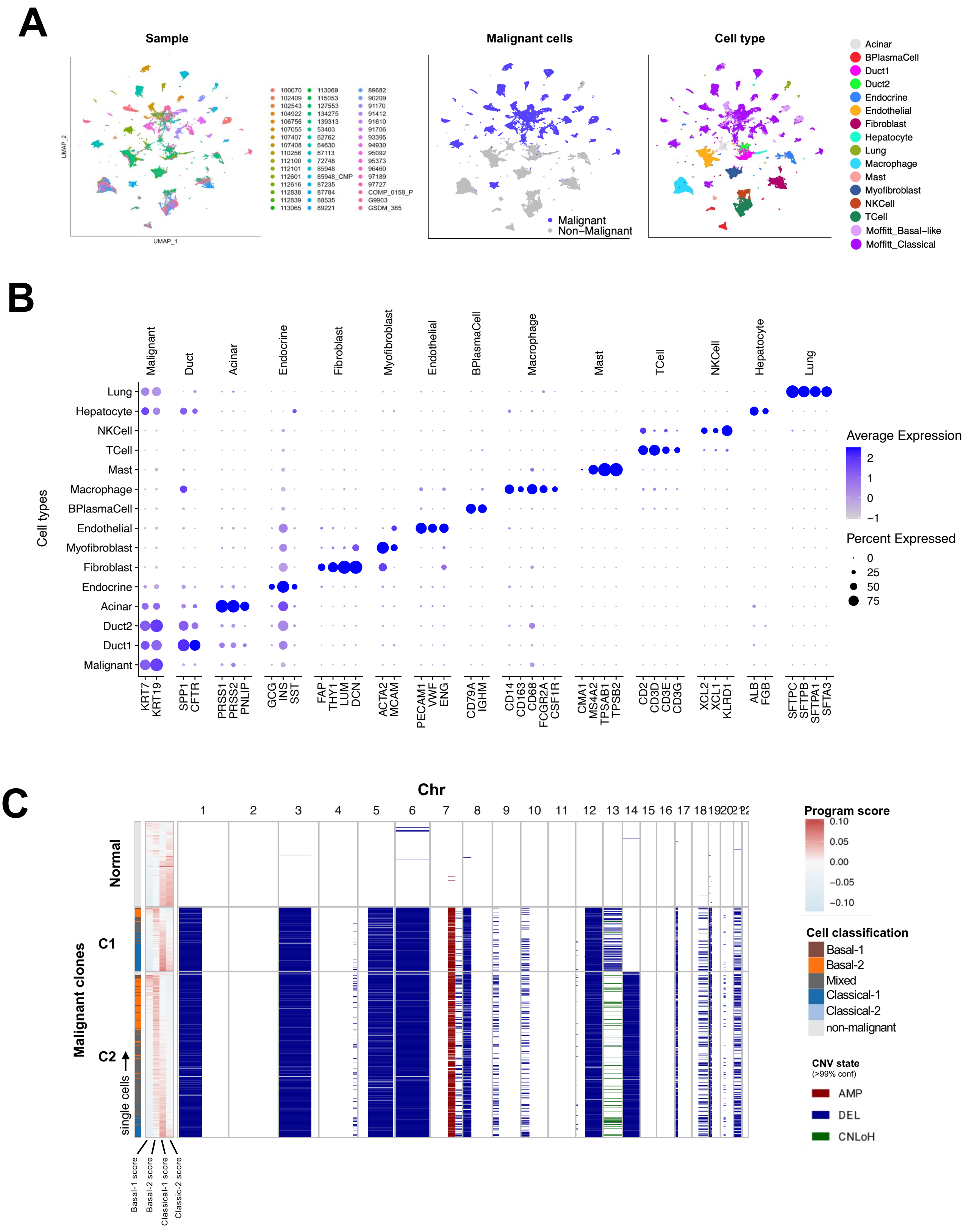

### Supplemental Figure 2

**A**

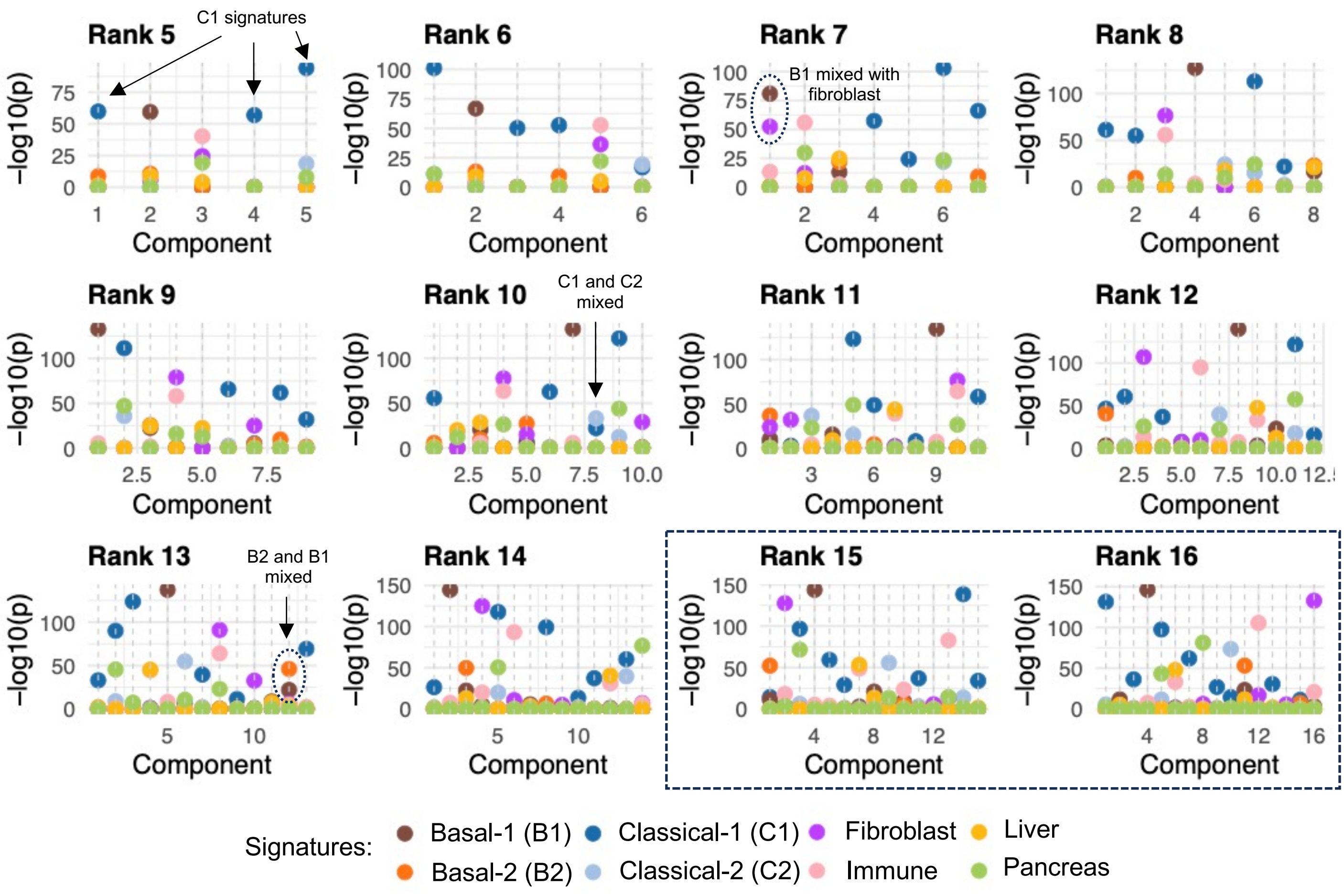


**B**

C2 signature

Rank 15

| | Genes | Score |
|---|---|---|
| 1 | REG3G | 0.99951969 |
| 2 | SPINK4 | 0.99608337 |
| 3 | CLCA1 | 0.99517981 |
| 4 | REG1B | 0.99438793 |
| 5 | MUC2 | 0.99223841 |
| 6 | PRSS3P1 | 0.98808737 |
| 7 | PRSS2 | 0.96015177 |
| 8 | GALNT8 | 0.96012464 |
| 9 | ITLN1 | 0.95821043 |
| 10 | CPA2 | 0.95318364 |
| 11 | CPA1 | 0.94767953 |
| 12 | RBPJL | 0.94575196 |
| 13 | PRSS1 | 0.9266556 |
| 14 | HEPACAM2 | 0.92084298 |
| 15 | ATOH1 | 0.90820234 |
| 16 | CLPS | 0.90476706 |
| 17 | REG1A | 0.89612821 |
| 18 | AQP12B | 0.89491053 |
| 19 | CELA3B | 0.88862096 |
| 20 | CELP | 0.87733344 |
| 21 | CEL | 0.87677097 |
| 22 | MEP1A | 0.86655797 |
| 23 | NXPE4 | 0.86023467 |
| 24 | FCGBP | 0.84186434 |
| 25 | AQP12A | 0.83800325 |
| 26 | PNLIP | 0.83345745 |
| 27 | KIF19 | 0.82677864 |
| 28 | KLHL32 | 0.80934612 |
| 29 | LRRC26 | 0.80891044 |
| 30 | NXPE2 | 0.80818758 |

Rank 16

| | Genes | Score |
|---|---|---|
| 1 | CLCA1 | 0.99837164 |
| 2 | SPINK4 | 0.99803856 |
| 3 | MUC2 | 0.9974501 |
| 4 | ITLN1 | 0.96604604 |
| 5 | COL2A1 | 0.96252904 |
| 6 | GALNT8 | 0.94957582 |
| 7 | ATOH1 | 0.94534805 |
| 8 | FCGBP | 0.88182751 |
| 9 | HEPACAM2 | 0.86841742 |
| 10 | MEP1A | 0.84520643 |
| 11 | CHST5 | 0.83080149 |
| 12 | NXPE2 | 0.82368338 |
| 13 | DAZ4 | 0.81040748 |
| 14 | REG4 | 0.8089723 |
| 15 | KLHL32 | 0.80715722 |
| 16 | NXPE4 | 0.80681708 |
| 17 | FABP2 | 0.80269426 |
| 18 | ITLN2 | 0.80182614 |
| 19 | CDX1 | 0.78715513 |
| 20 | NBPF5P | 0.78351312 |
| 21 | NXPE1 | 0.78253267 |
| 22 | LRRC26 | 0.78008881 |
| 23 | GUCY2C | 0.77912841 |
| 24 | KIF19 | 0.77052877 |
| 25 | NBPF6 | 0.75888643 |
| 26 | GUCA2A | 0.75291202 |
| 27 | NOS2 | 0.7526965 |
| 28 | XPNPEP2 | 0.73282996 |
| 29 | NBPF4 | 0.72712549 |
| 30 | AP000867.4 | 0.7184794 |

### Supplemental Figure 3

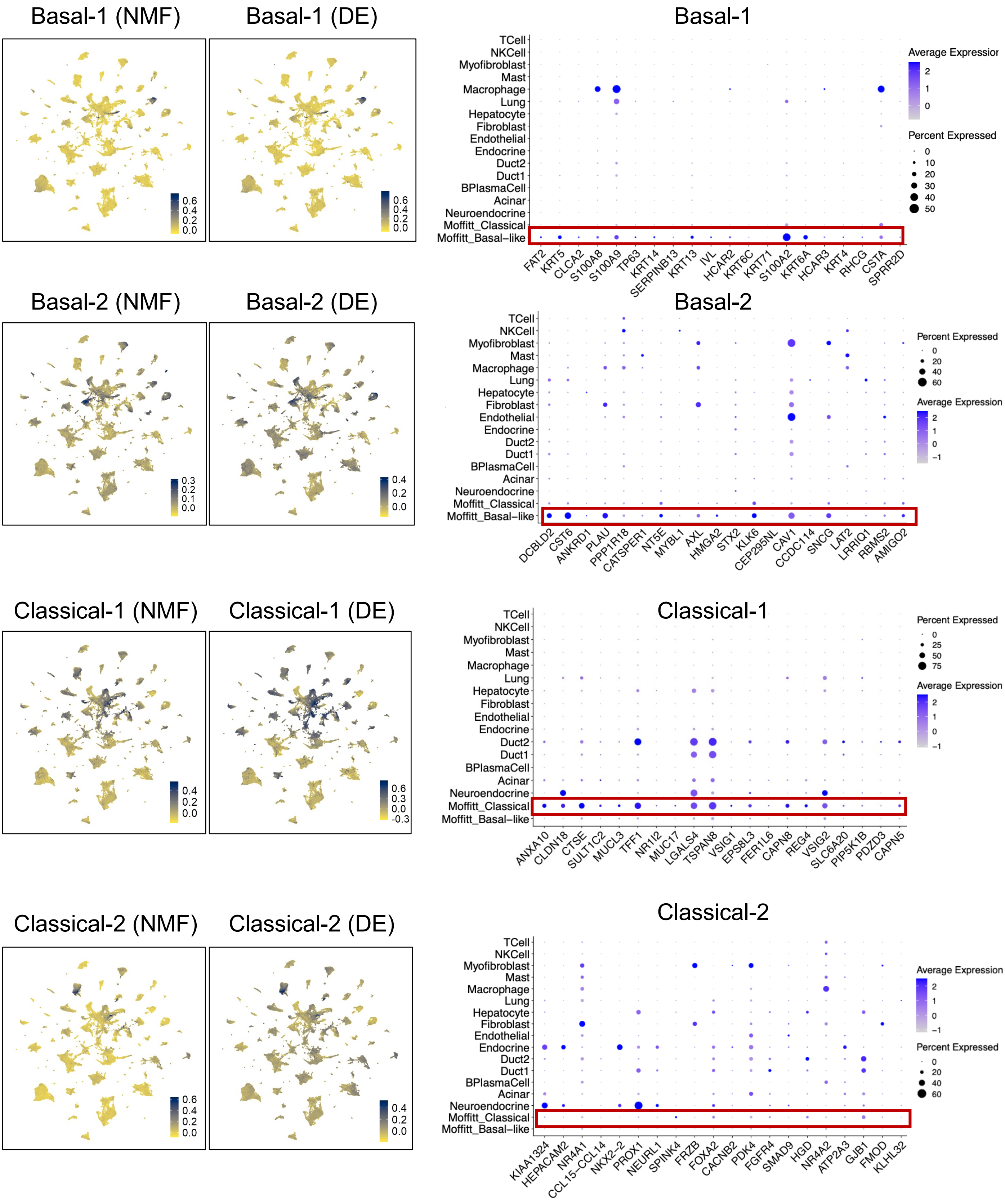

### Supplemental Figure 4

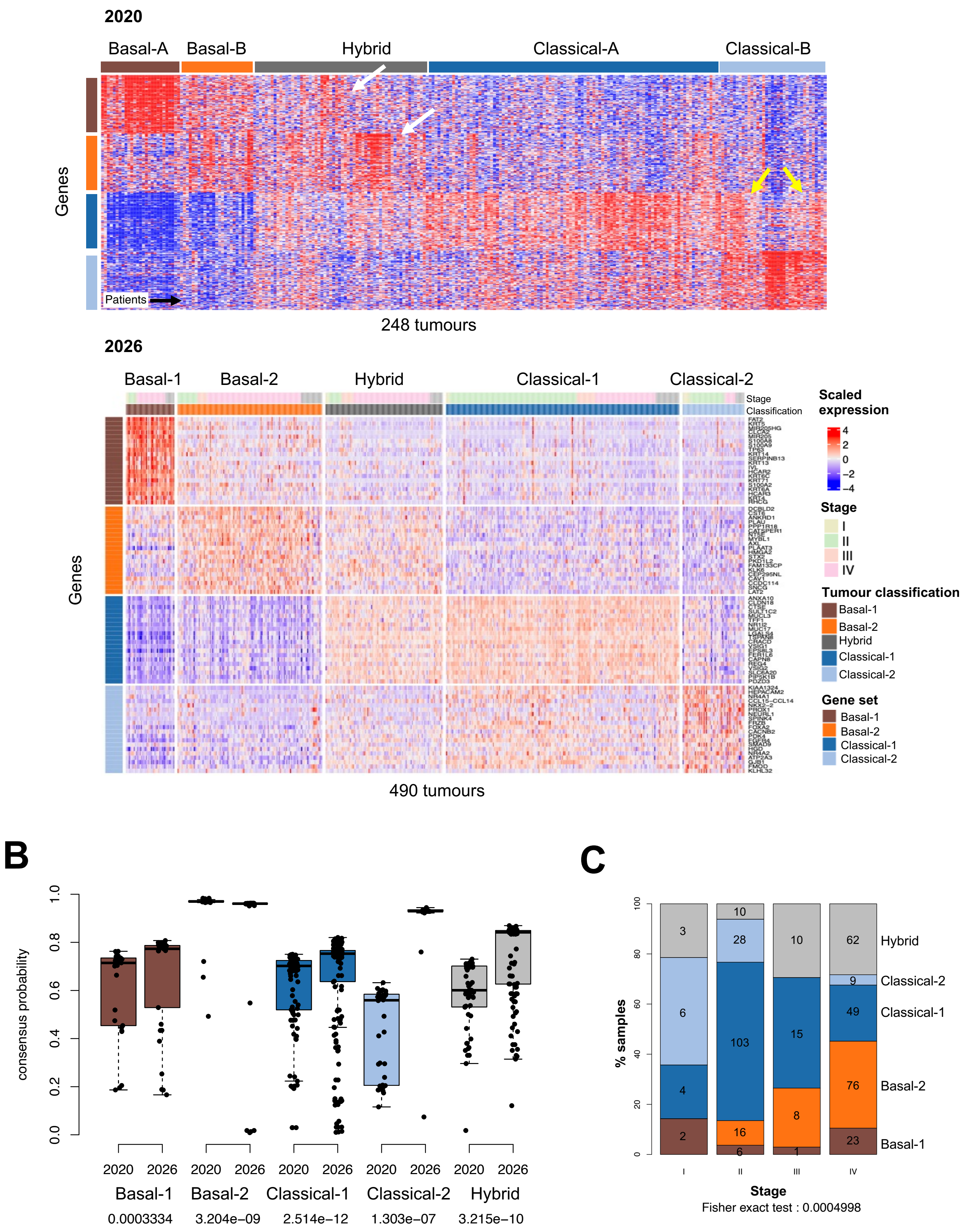

### Supplemental Figure 5

A

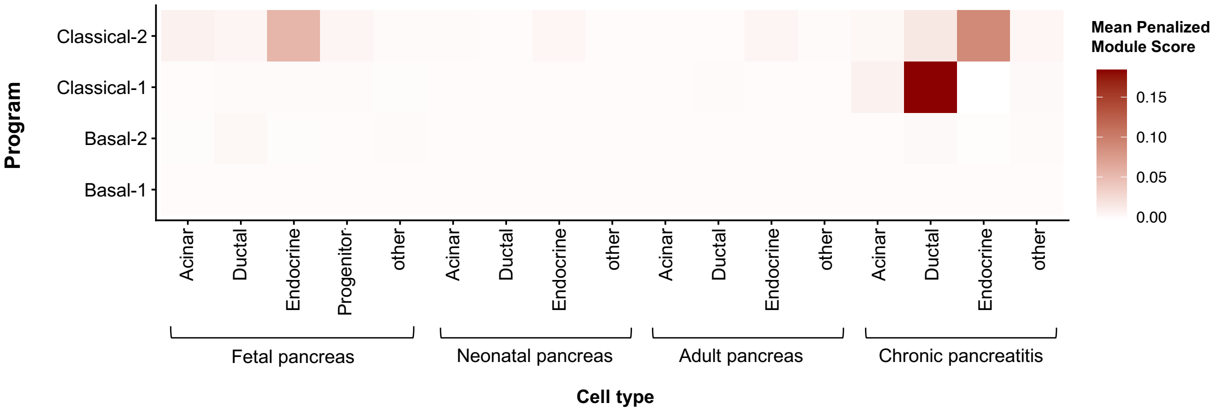


B

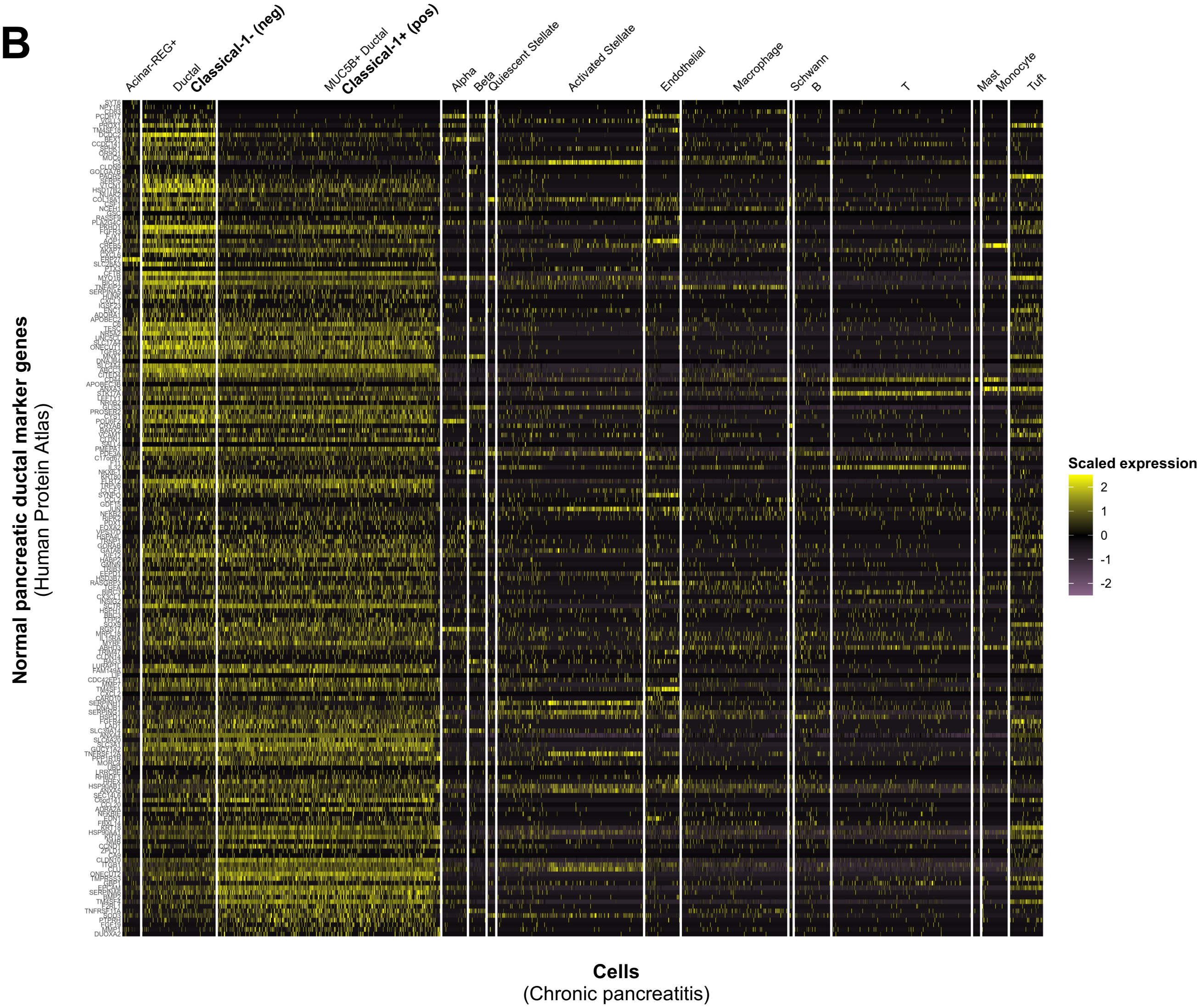

### Supplemental Figure 5 cont'd

**C**

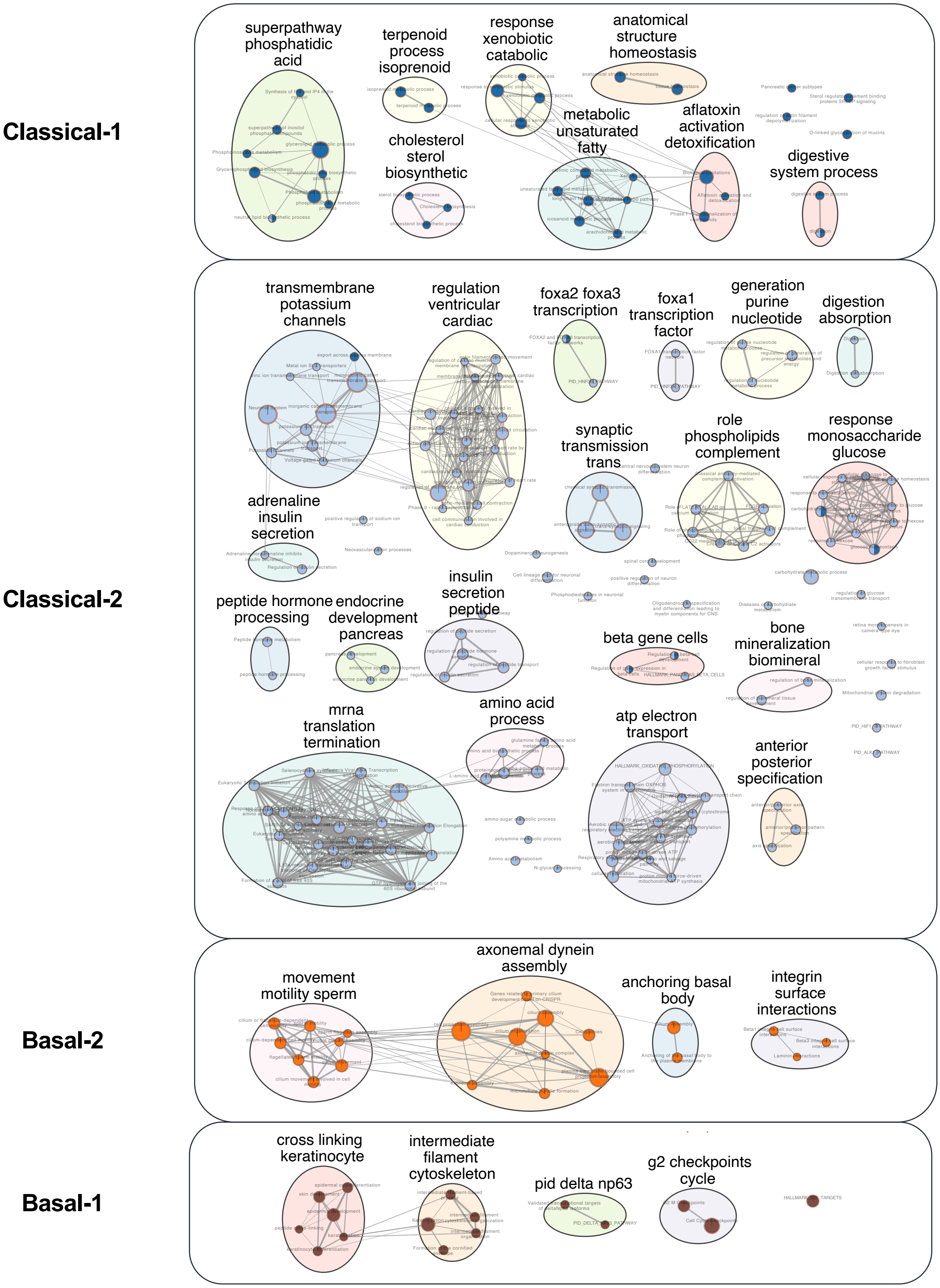

### Supplemental Figure 6

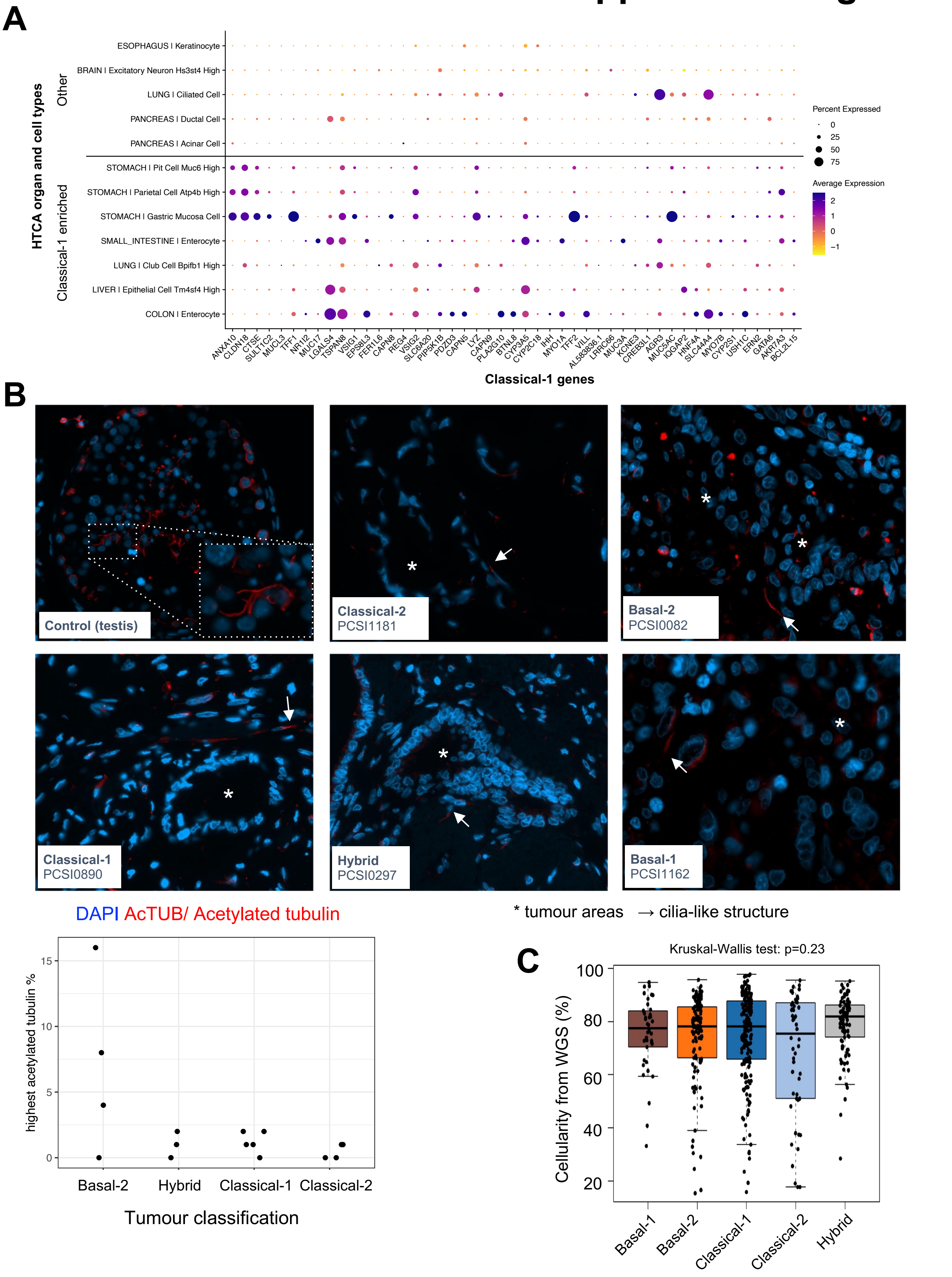

### Supplemental Figure 7

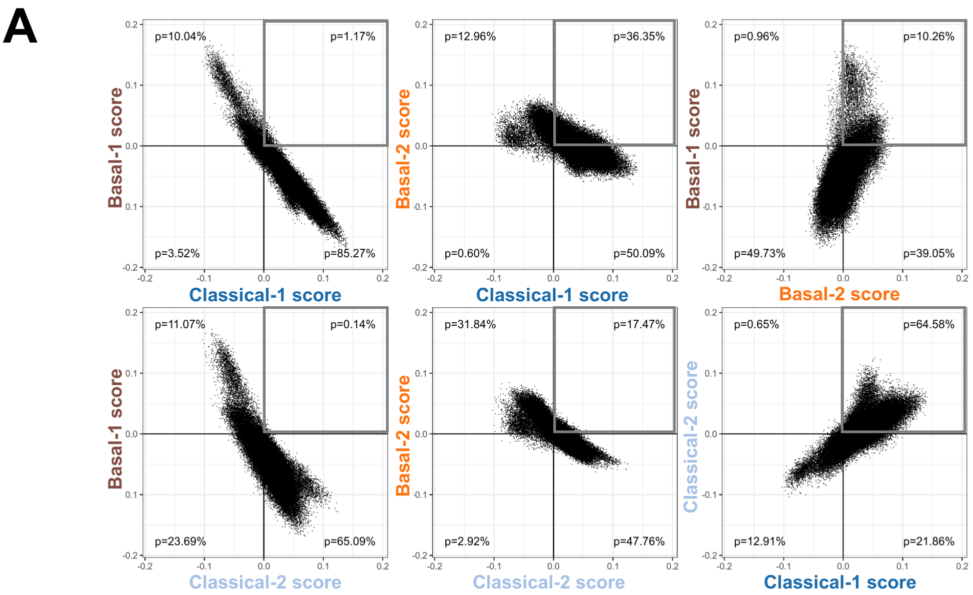


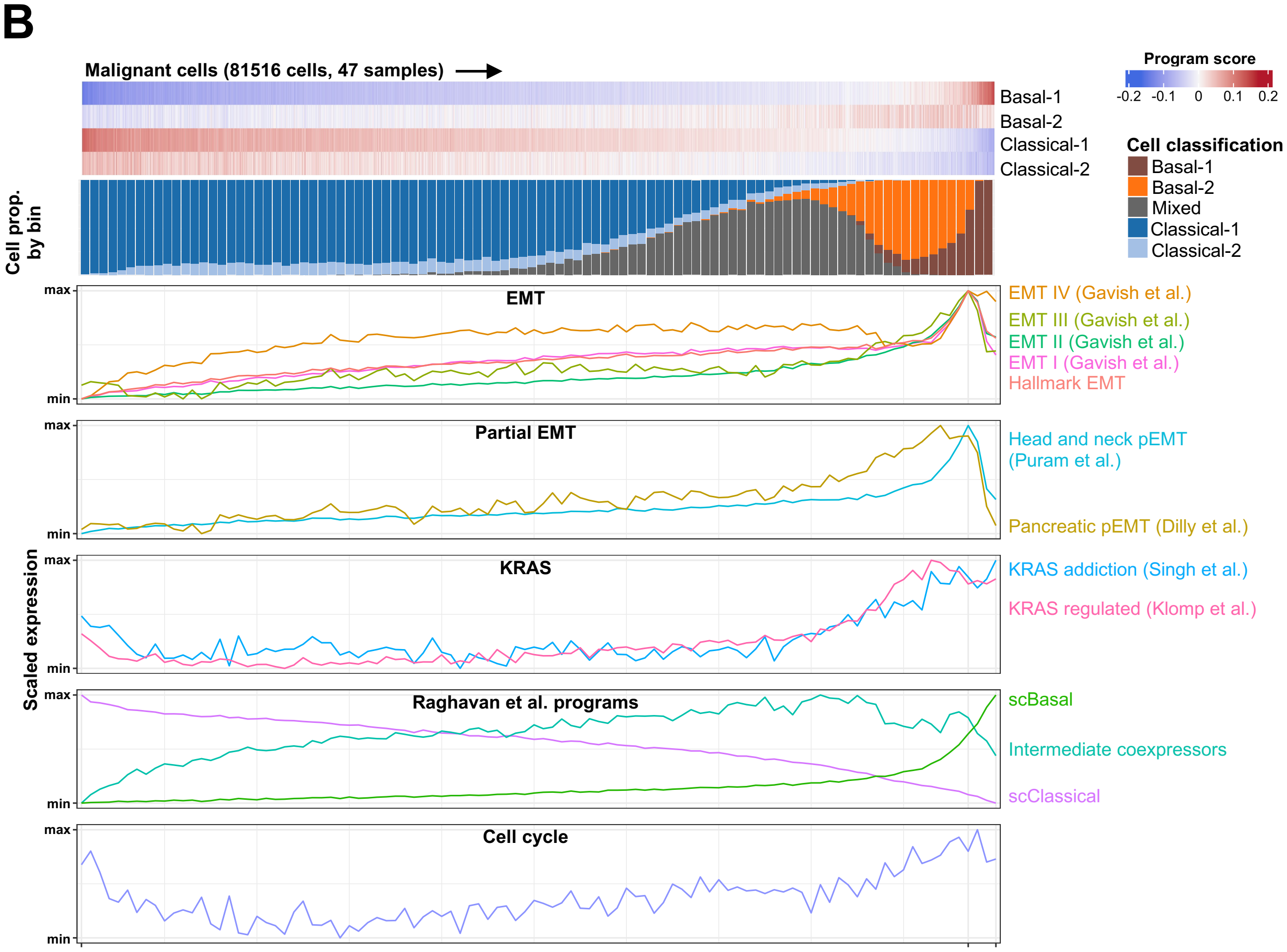

### Supplemental Figure 7 cont'd

**C** Malignant cells from Loveless et al. (229 samples)

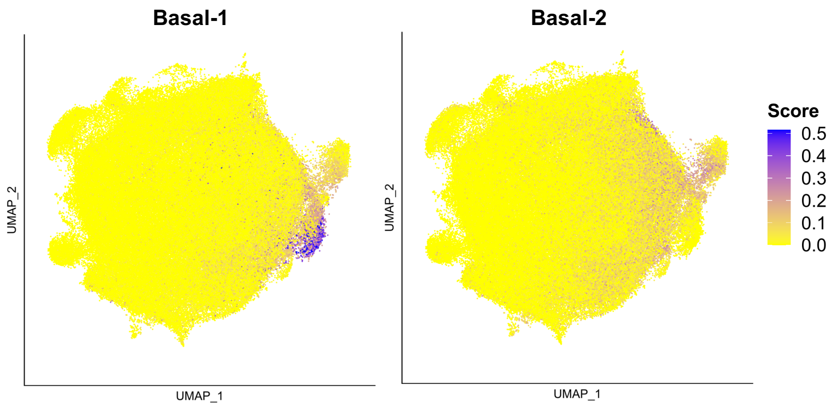


**D**

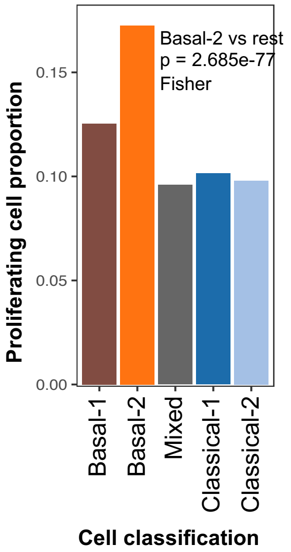


**E**

KRAS Imbalance

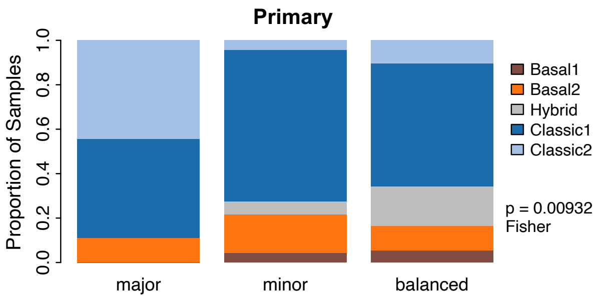


| | Basal1 | Basal2 | Hybrid | Classic1 | Classic2 |
|---|---|---|---|---|---|
| *major* | 0 | 1 | 0 | 4 | 4 |
| *minor* | 3 | 12 | 4 | 47 | 3 |
| *balanced* | 9 | 18 | 29 | 90 | 17 |

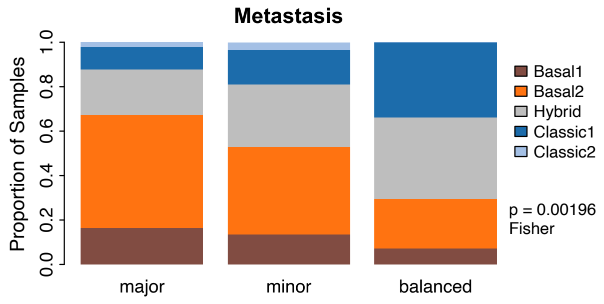


| | Basal1 | Basal2 | Hybrid | Classic1 | Classic2 |
|---|---|---|---|---|---|
| *major* | 8 | 25 | 10 | 5 | 1 |
| *minor* | 12 | 35 | 25 | 14 | 3 |
| *balanced* | 5 | 15 | 25 | 23 | 0 |

**F**

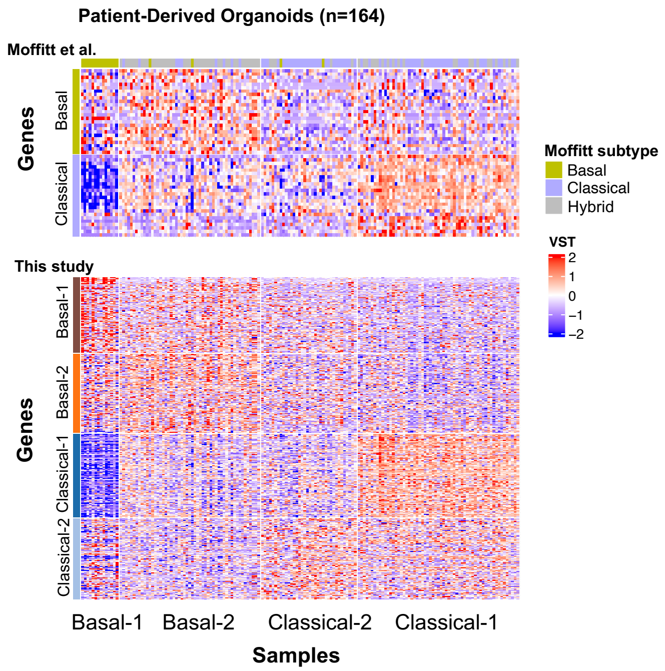

### Supplemental Figure 8

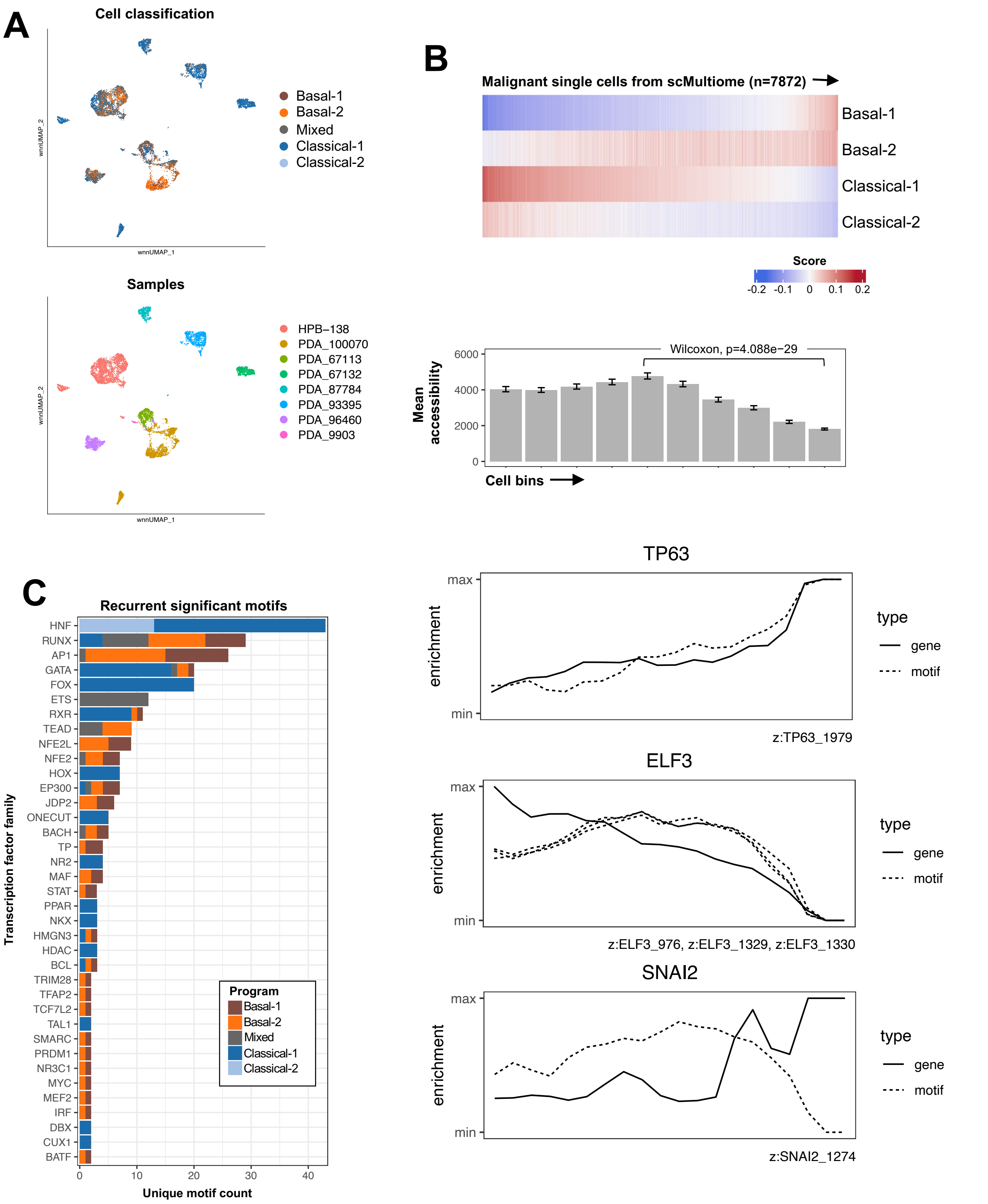

### Supplemental Figure 9

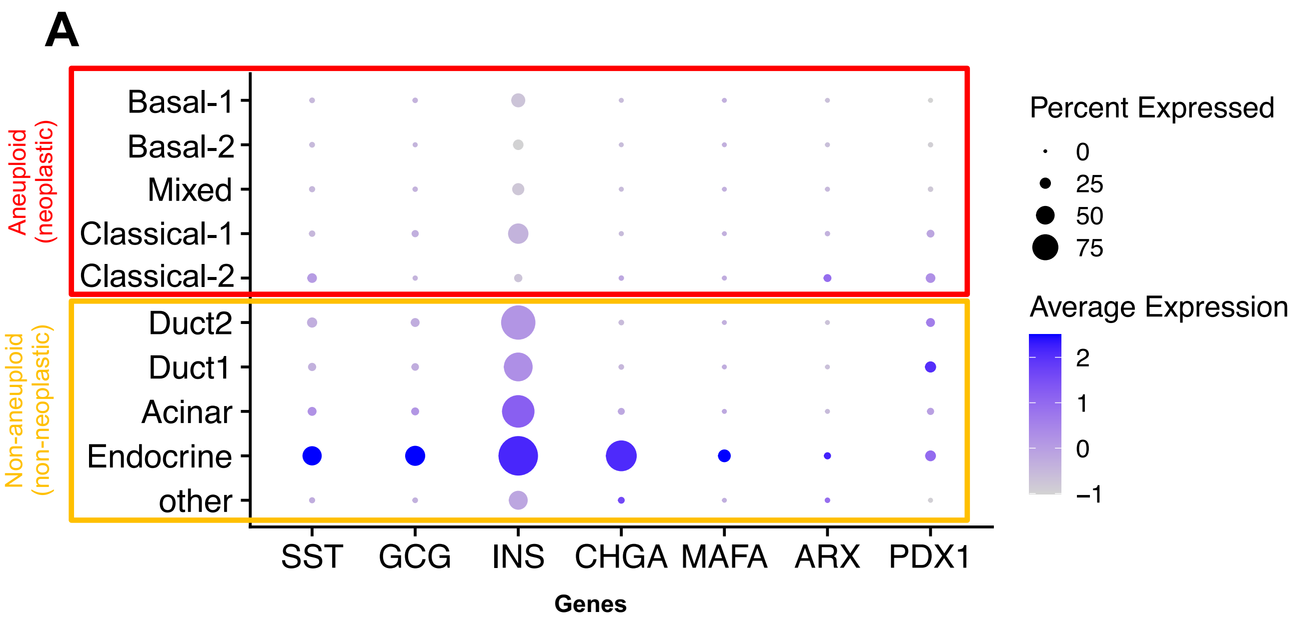


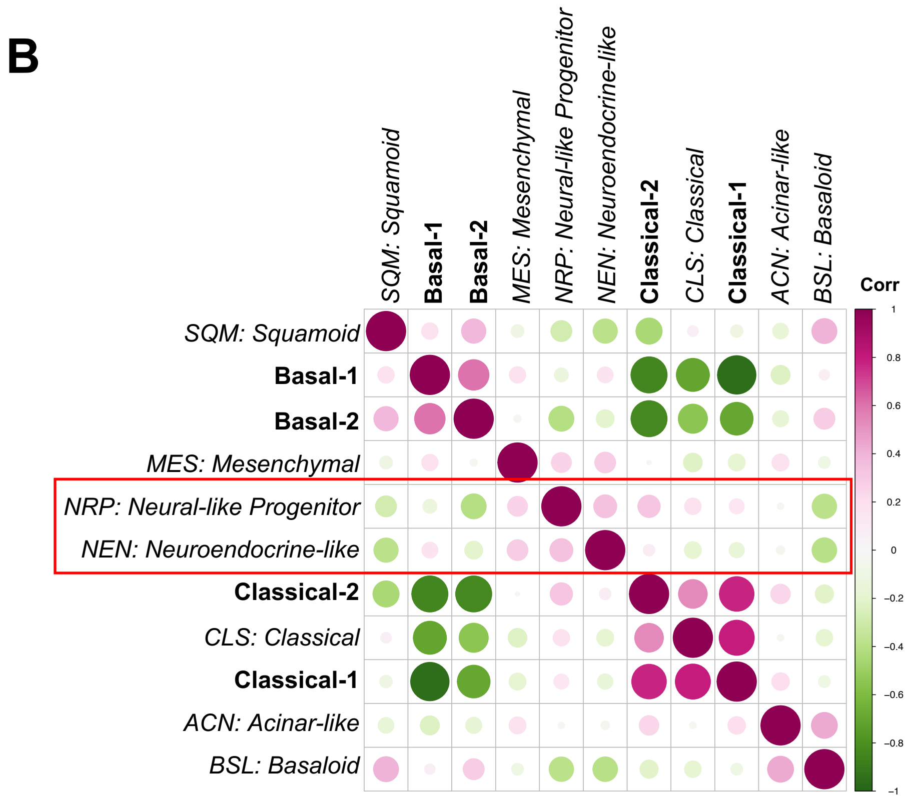

### Supplementary Figure 10

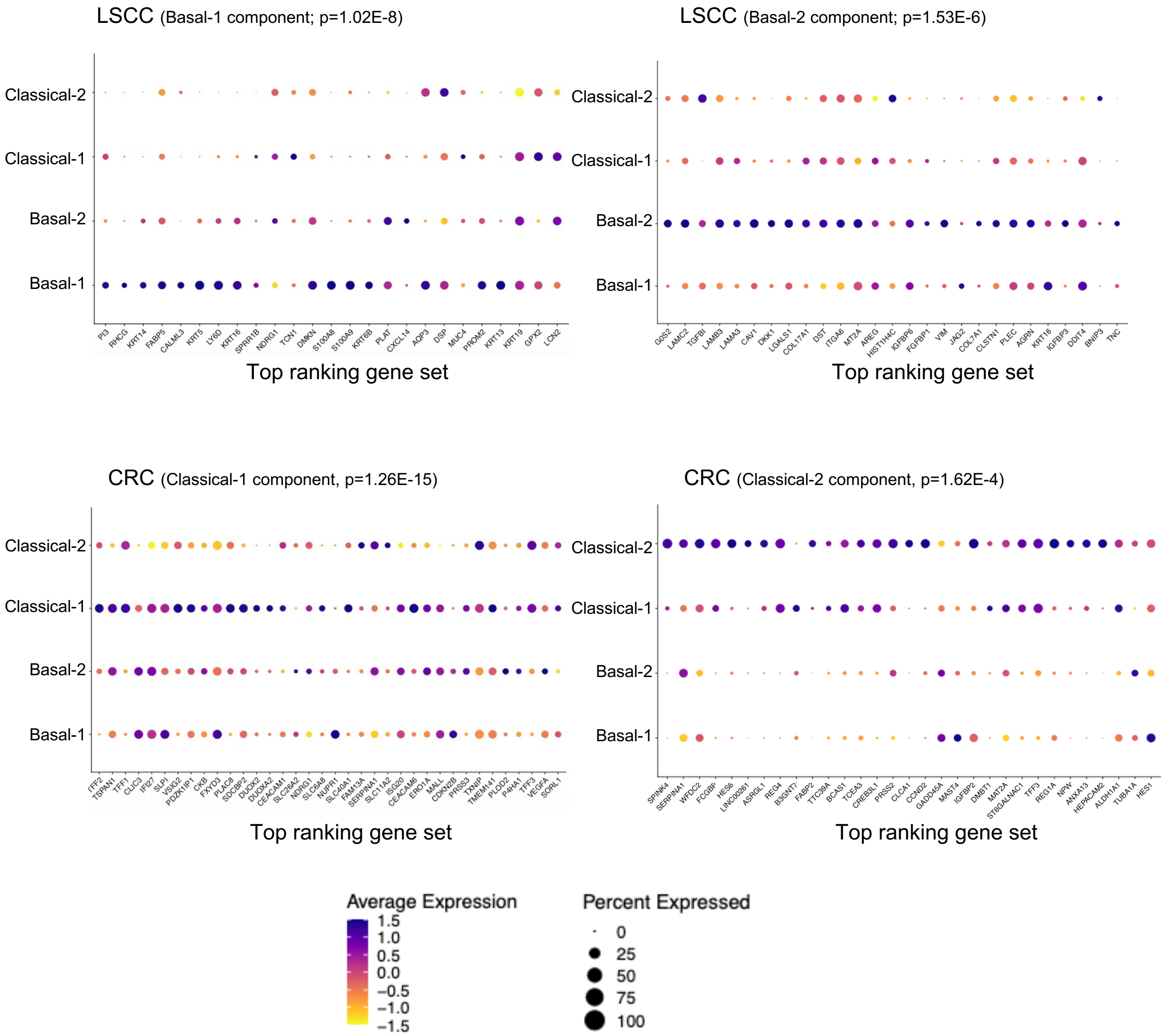

### Supplementary Figure 11

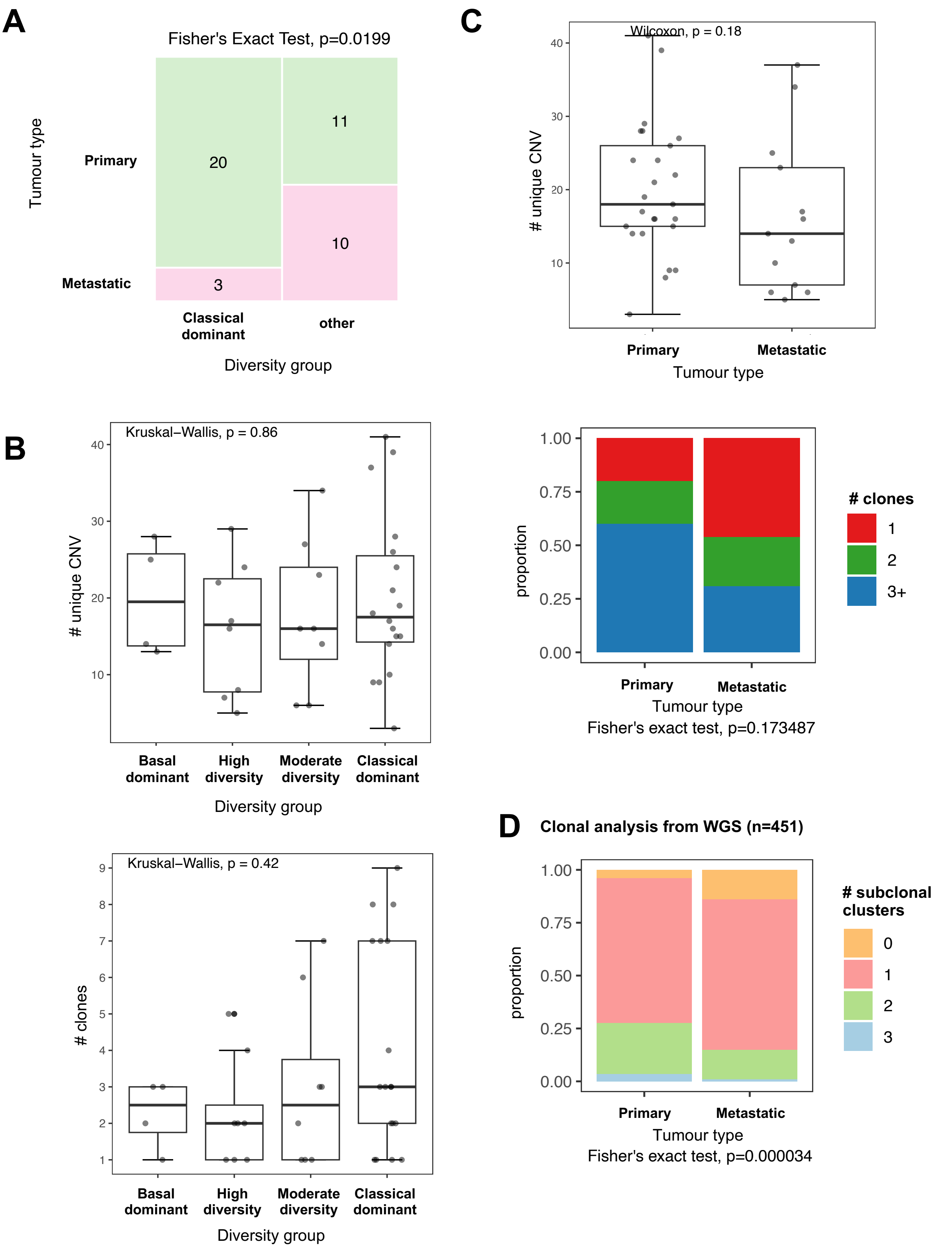

### Supplementary Figure 12

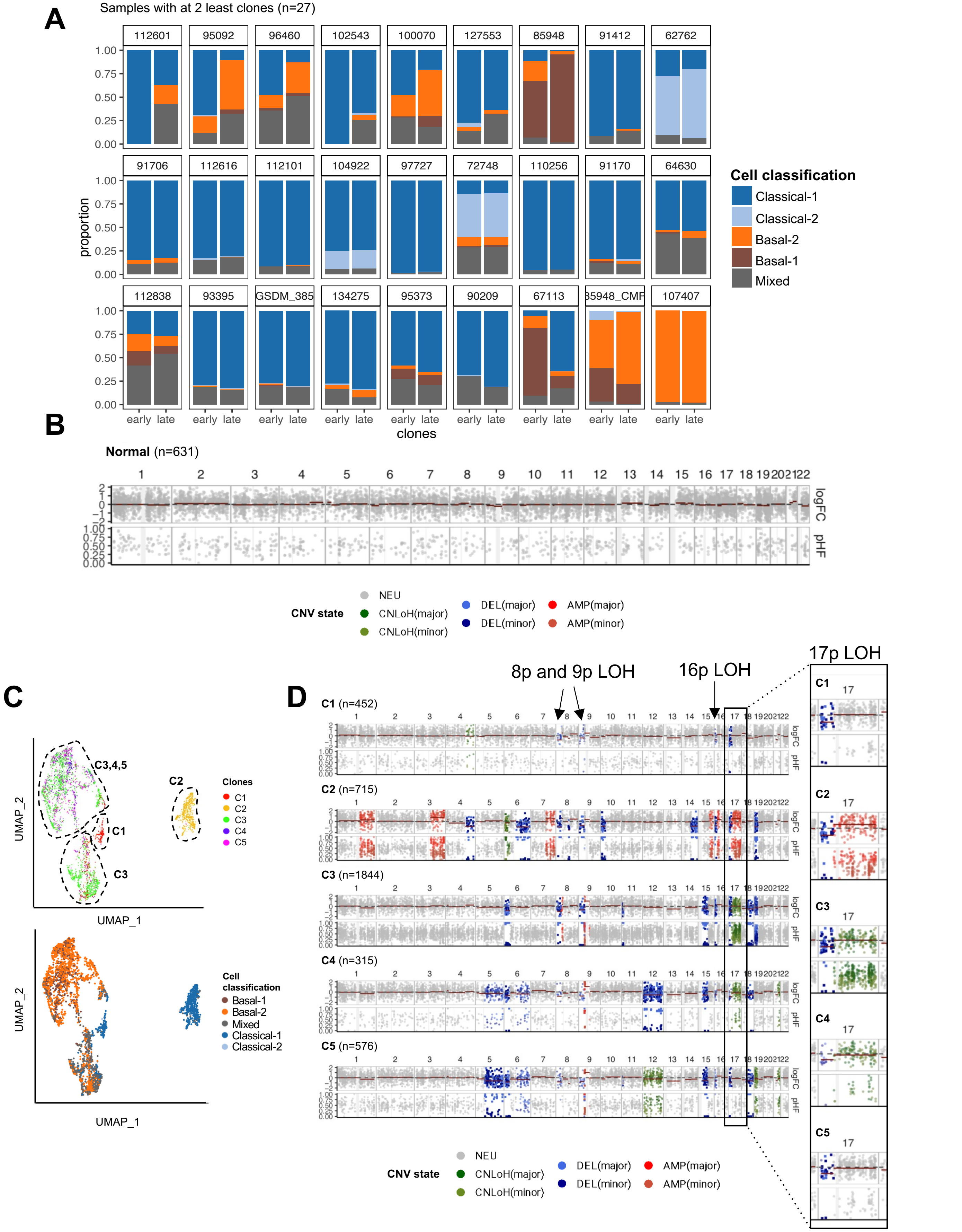

### Supplemental Figure 13

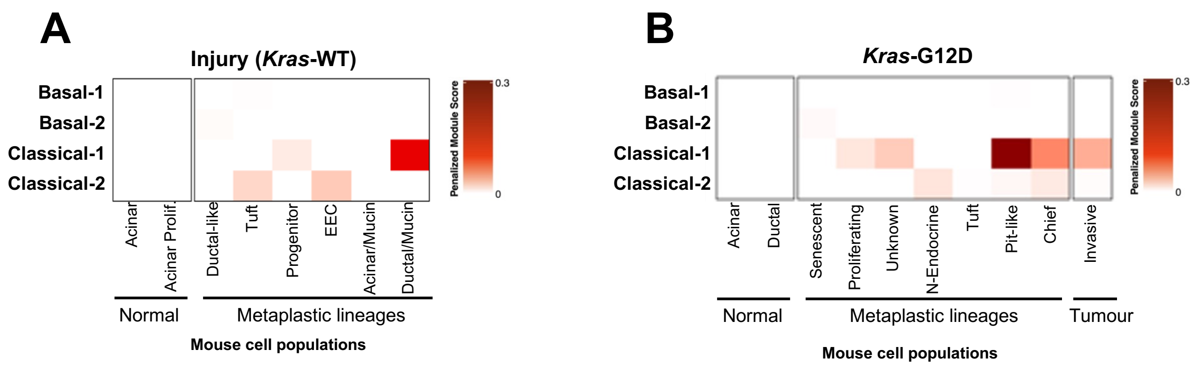

### Supplemental Figure 14

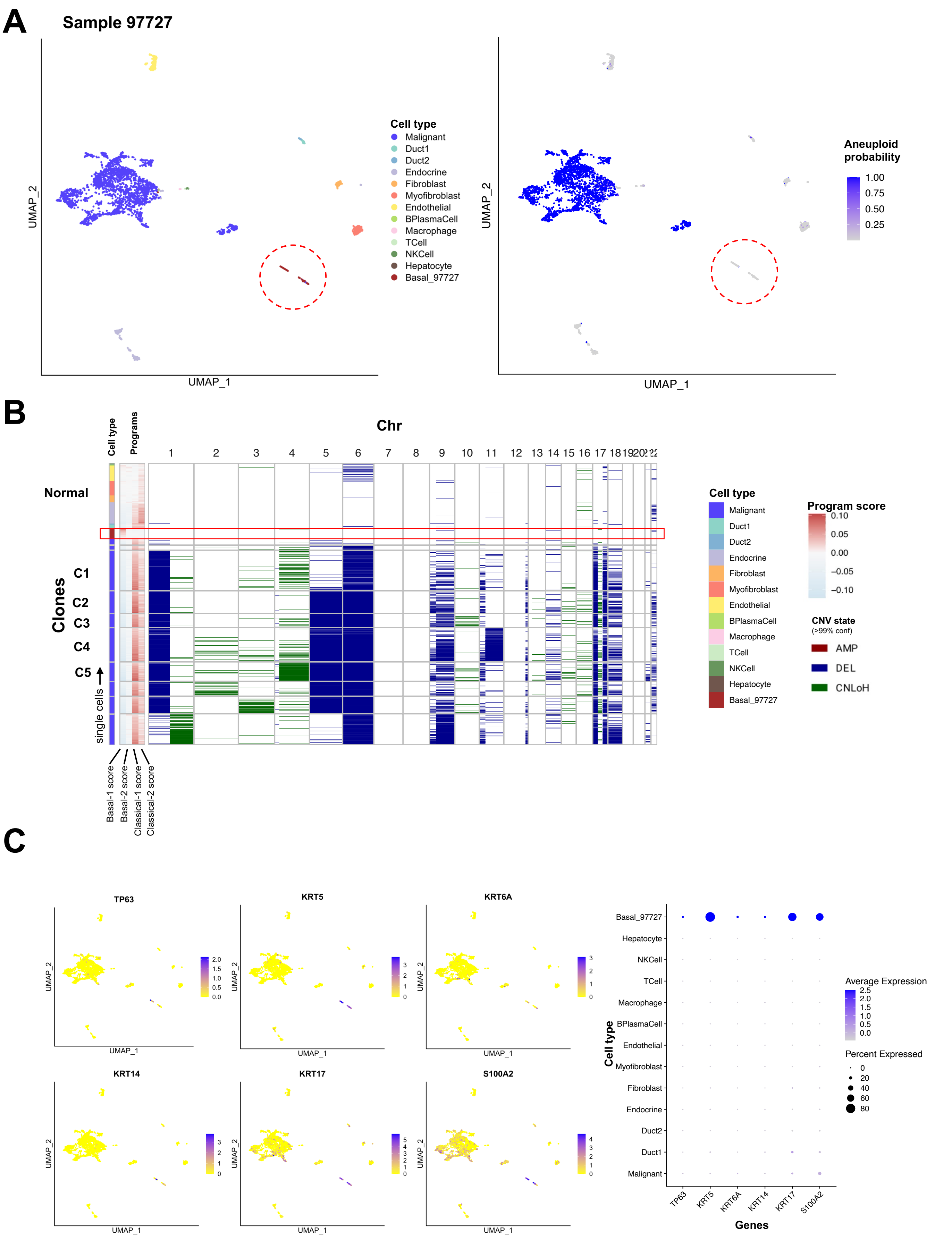

### Supplemental Figure 15

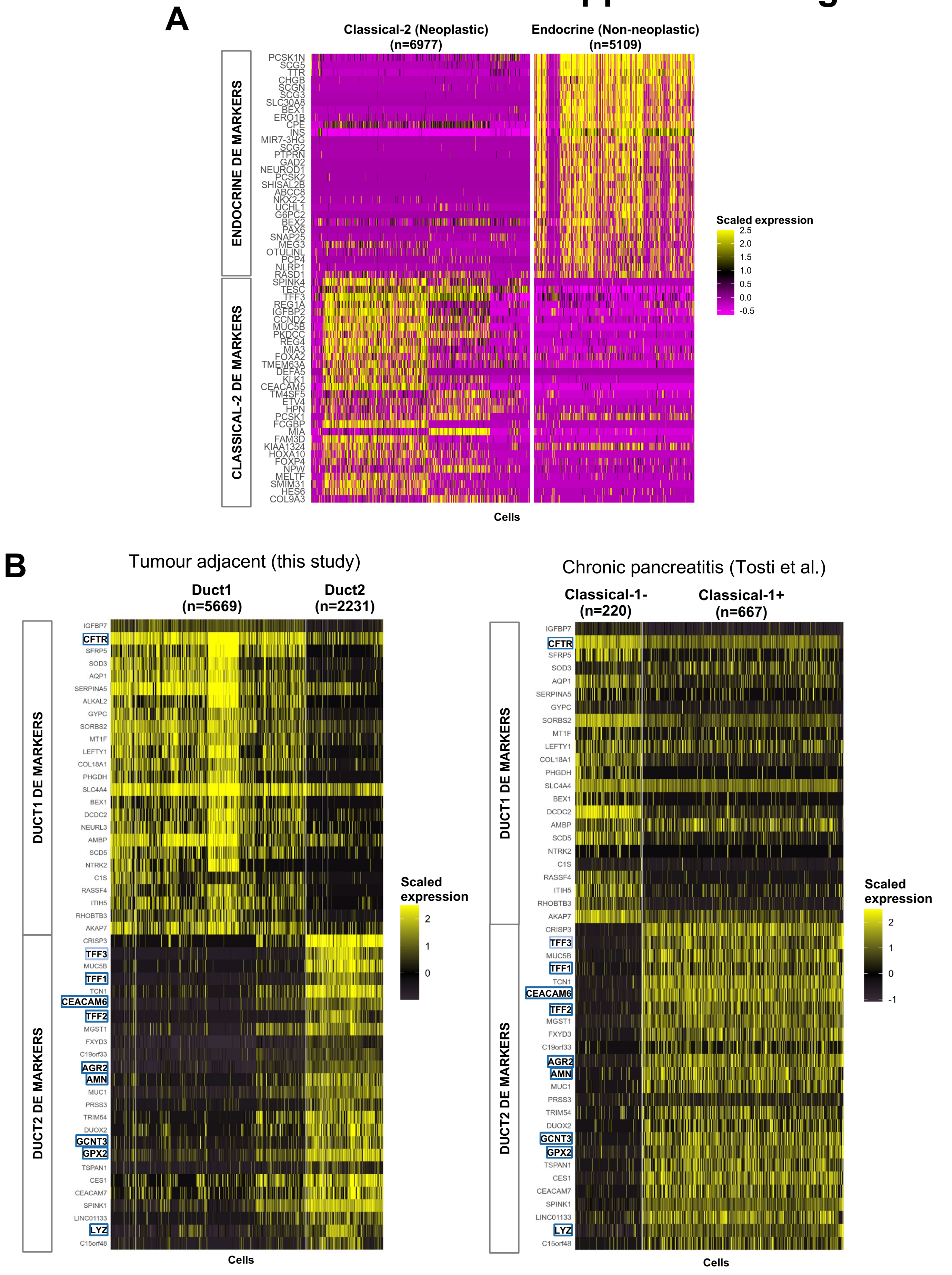

### Supplemental Figure 16

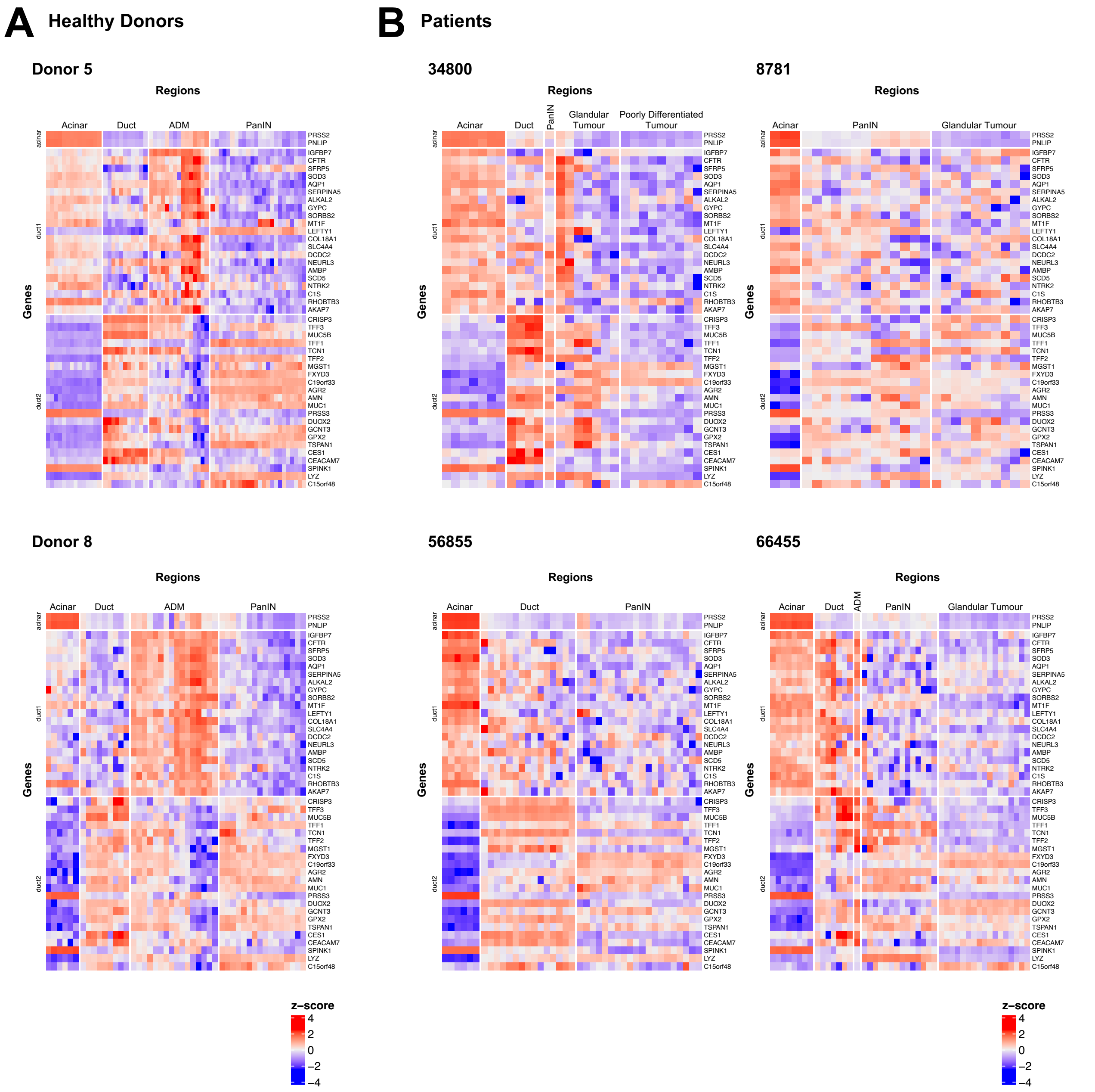

### Supplementary Figure 17

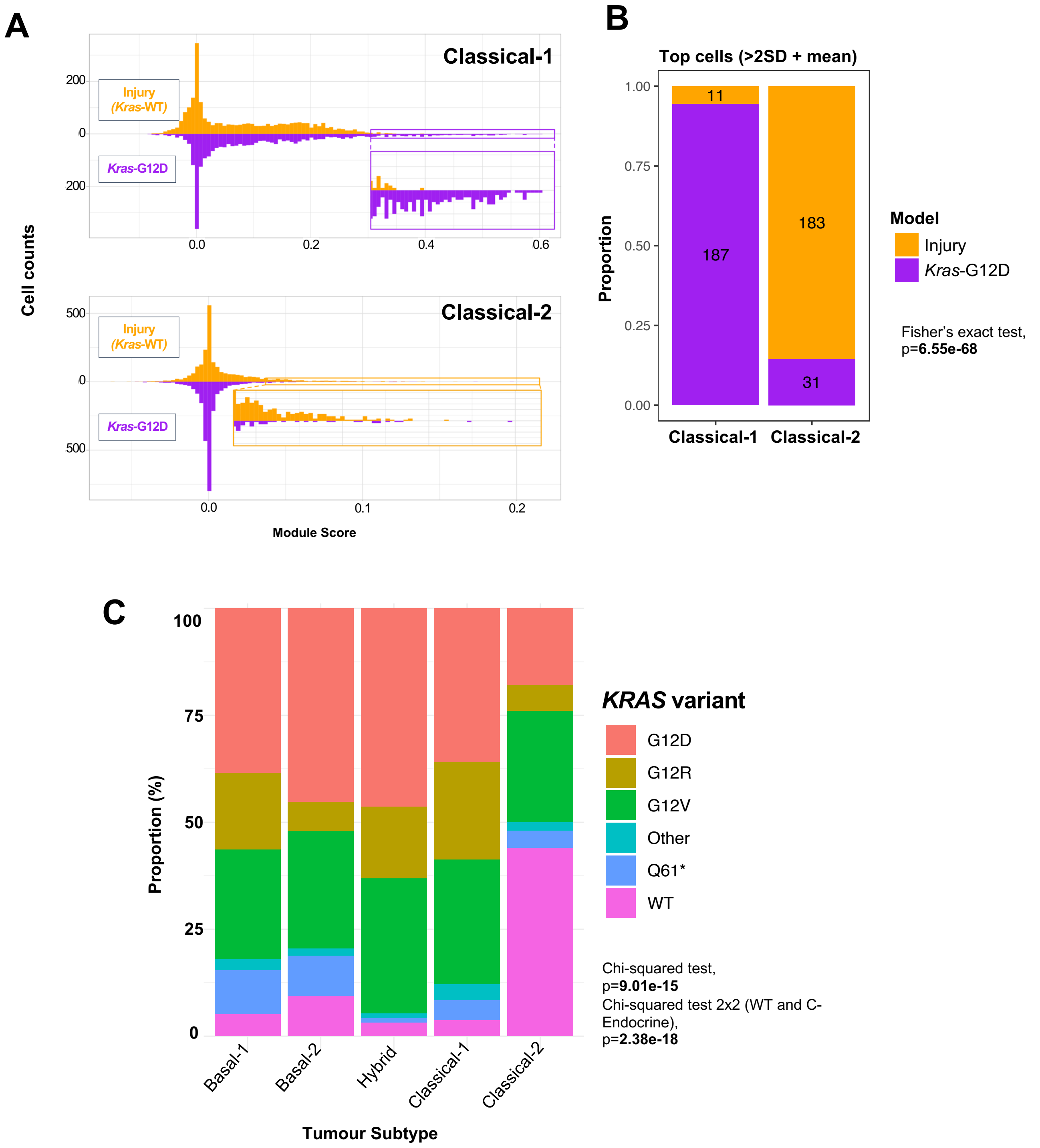

### Supplementary Fig 18

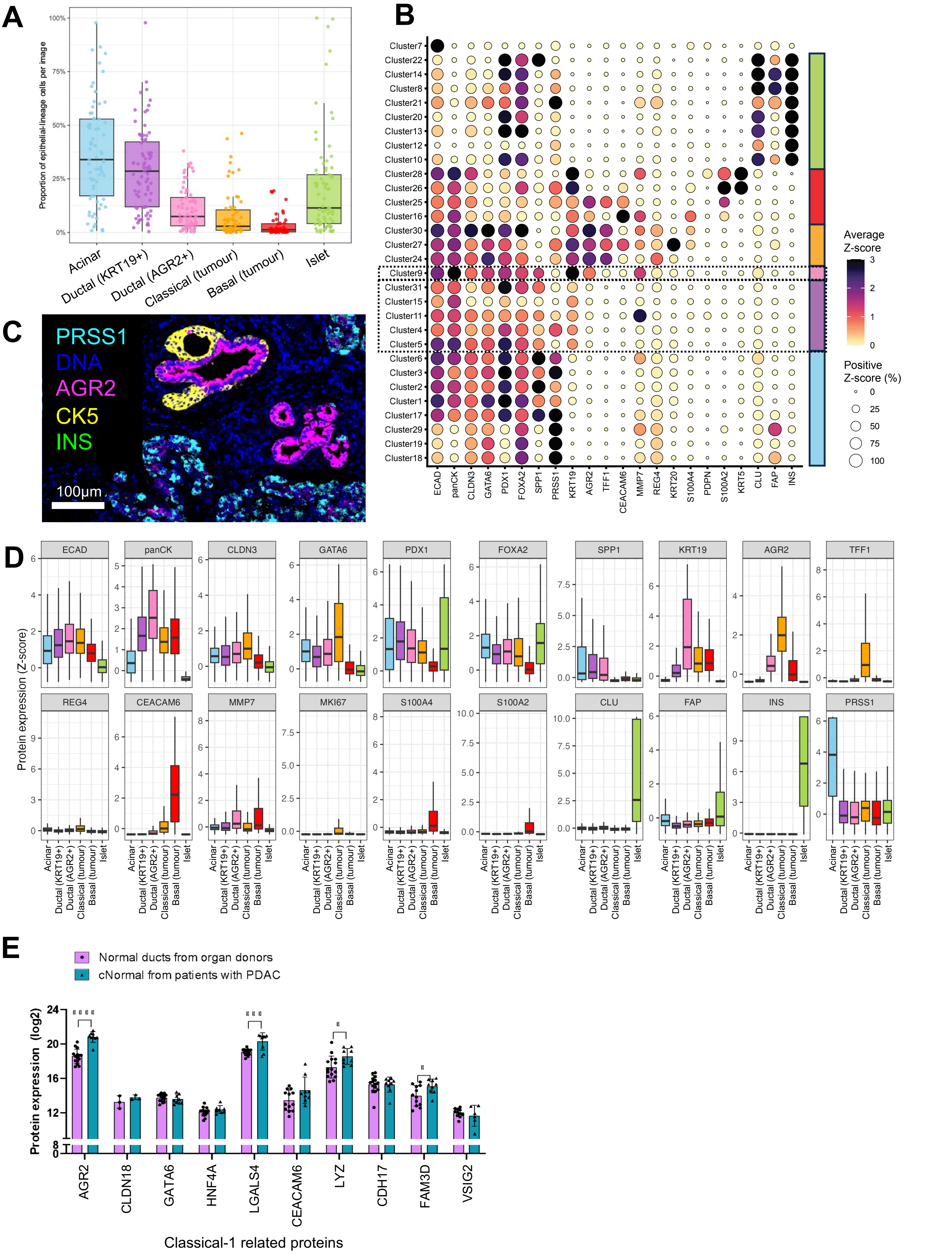
